# Human cells contain myriad excised linear intron RNAs with links to gene regulation and potential utility as biomarkers

**DOI:** 10.1101/2020.09.07.285114

**Authors:** Jun Yao, Hengyi Xu, Elizabeth A. Ferrick-Kiddie, Ryan M. Nottingham, Douglas C. Wu, Manuel Ares, Alan M. Lambowitz

## Abstract

Previous Thermostable Group II Intron Reverse Transcriptase sequencing (TGIRT-seq) found that human plasma contains short (≤300 nt) structured full-length excised linear intron RNAs (FLEXIs) with potential utility as blood-based biomarkers. Here, TGIRT-seq identified >9,000 different FLEXIs in human cells, including relatively abundant FLEXIs with cell-type-specific expression patterns. Analysis of published CLIP-seq datasets identified 126 RNA-binding proteins (RBPs) that bind different FLEXIs, including 53 RBPs that bind ≥30 FLEXIs. These RBPs included splicing factors, transcription factors, a chromatin remodeling protein, cellular growth regulators, and proteins with cytoplasmic functions. Analysis of published datasets identified subsets of these RBPs that bind at FLEXI splice sites and impact alternative splicing or FLEXI host gene mRNA levels. Hierarchical clustering identified 6 subsets of RBPs whose binding sites were co-enriched in different subsets of FLEXIs: AGO1-4 and DICER, including but not limited to annotated agotrons and mirtron pre-miRNAs; DKC1, NOLC1, SMNDC1, and AATF (Apoptosis Antagonizing Transcription Factor), including but not limited to FLEXIs encoding snoRNAs; two sets of alternative splicing factors; and two sets that included RBPs with cytoplasmic functions (*e.g.*, LARP4, PABPC4, METAP2, and ZNF622) together with nuclear regulatory proteins. The subsets of host genes encoding FLEXIs that bind these RBPs were enriched with non-FLEXI other short and long introns that bind the same RBPs, and TGIRT-seq of nuclear and cytoplasmic fractions from 4 cell lines showed cytoplasmic enrichment of FLEXIs with binding sites for RBPs that function in the cytoplasm. Collectively, our findings suggest that different subsets of RBPs bind FLEXIs and other introns to coordinately regulate the expression of functionally related host genes and then co-localize with stably bound FLEXIs to different intracellular locations. The cell-type specific expression of relatively abundant FLEXIs suggests utility as cellular as well as blood-based RNA biomarkers.

## Introduction

Most protein-coding genes in eukaryotes consist of coding regions (exons) separated by noncoding regions (introns), which are removed by RNA splicing to produce mRNAs. RNA splicing is performed by the spliceosome, a large ribonucleoprotein complex that catalyzes transesterification reactions yielding ligated exons and an excised intron lariat RNA, whose 5’ end is linked to a branch-point nucleotide near its 3’ end by a 2’-5’-phosphodiester bond (1). After splicing, this bond is typically hydrolyzed by the debranching enzyme DBR1, and the resulting linear intron RNA is rapidly degraded by exonucleases (2). Several classes of excised intron RNAs persist after excision. These include trimmed lariat RNAs that lack a 3’ tail; circularized intron RNAs whose 5’ and 3’ nucleotides are putatively linked by a 3’-5’ phosphodiester bond; and full-length excised linear intron RNAs (FLEXIs), with members of all 3 classes found to have cellular functions or clinical correlations (3–14). FLEXIs include a group of yeast excised linear intron RNAs that contribute to cell growth regulation by sequestering spliceosomal proteins in stationary phase or under other stress conditions (11, 15). Other FLEXIs include mirtron pre-miRNAs, structured excised intron RNAs that are debranched by DBR1 and processed by DICER into annotated miRNAs, and agotrons, structured excised intron RNAs that bind AGO2 and repress target mRNAs in a miRNA-like manner (16–19).

Previously, we identified 44 different short (≤300 nt) FLEXIs in human plasma by using Thermostable Group II Intron Reverse Transcriptase sequencing (TGIRT-seq), a method that enables continuous end-to-end sequencing reads of structured RNAs (20). Plasma FLEXIs were identified by peak calling as discrete blocks of continuous sequencing reads of excised introns beginning and ending within 3 nucleotides (nt) of annotated splice sites, with relatively large subsets corresponding to annotated agotrons or mirtron pre-miRNAs (20). Almost all of the plasma FLEXIs (>95%) had stable predicted secondary structures (ΔG ≤ -30 kcal/mol), with ∼50% containing a CLIP-seq-identified binding site for one or more RNA-binding proteins (RBPs) that may help protect FLEXIs from plasma RNases. These findings suggested that FLEXIs might have utility as RNA biomarkers in liquid biopsies (20).

Despite the precedents of agotrons and mirtron pre-miRNAs, human FLEXIs have remained largely unexplored. Here, TGIRT-seq of RNAs present in human cell lines identified >9,000 FLEXIs, subsets of which had cell-type specific expression patterns, suggesting utility as cellular RNA biomarker. Analysis of published CLIP-seq datasets identified subsets of RBPs that bind ≥30 FLEXIs that were enriched in different subsets of functionally related host genes with other short and long intron that have binding sites for the same RBPs. The latter findings suggest previously unidentified mechanisms for co-regulation of functionally related gene sets and provide a road map for further analysis of these mechanisms as well as FLEXI-RBP interactions and cellular functions.

## Results

### Identification of FLEXIs in cellular RNA preparations

A search of the human genome (Ensembl GRCh38 Release 93 annotations; http://www.ensembl.org) identified 51,645 short introns (≤300 nt) in 12,020 different genes that could potentially give rise to FLEXIs. To investigate which short introns give rise to FLEXIs, we carried out TGIRT-seq (21, 22) of rRNA-depleted, unfragmented HEK-293T, K-562, and HeLa S3 whole-cell RNAs and Universal Human Reference RNA (UHRR), a commercial mixture of whole-cell RNAs from 10 human cell lines (Fig. 1). TGIRT-seq is well-suited for the identification of FLEXIs as it employs a TGIRT enzyme (GsI-IIC RT aka TGIRT-III) that can efficiently template-switch from an RNA-seq adapter to initiate cDNA synthesis directly at the 3’-nucleotide of target RNAs and then reverse transcribe processively to yield full-length cDNAs that give end-to-end sequencing reads of structured RNAs (23–26). Because this TGIRT enzyme does not efficiently read through the 3’-poly(A) tails of mature mRNAs, most of the reads in the TGIRT-seq datasets of unfragmented whole-cell RNAs corresponded to full-length mature tRNAs and other small non-coding RNAs (sncRNAs; Fig. S1 and S2).

**Figure 1.**
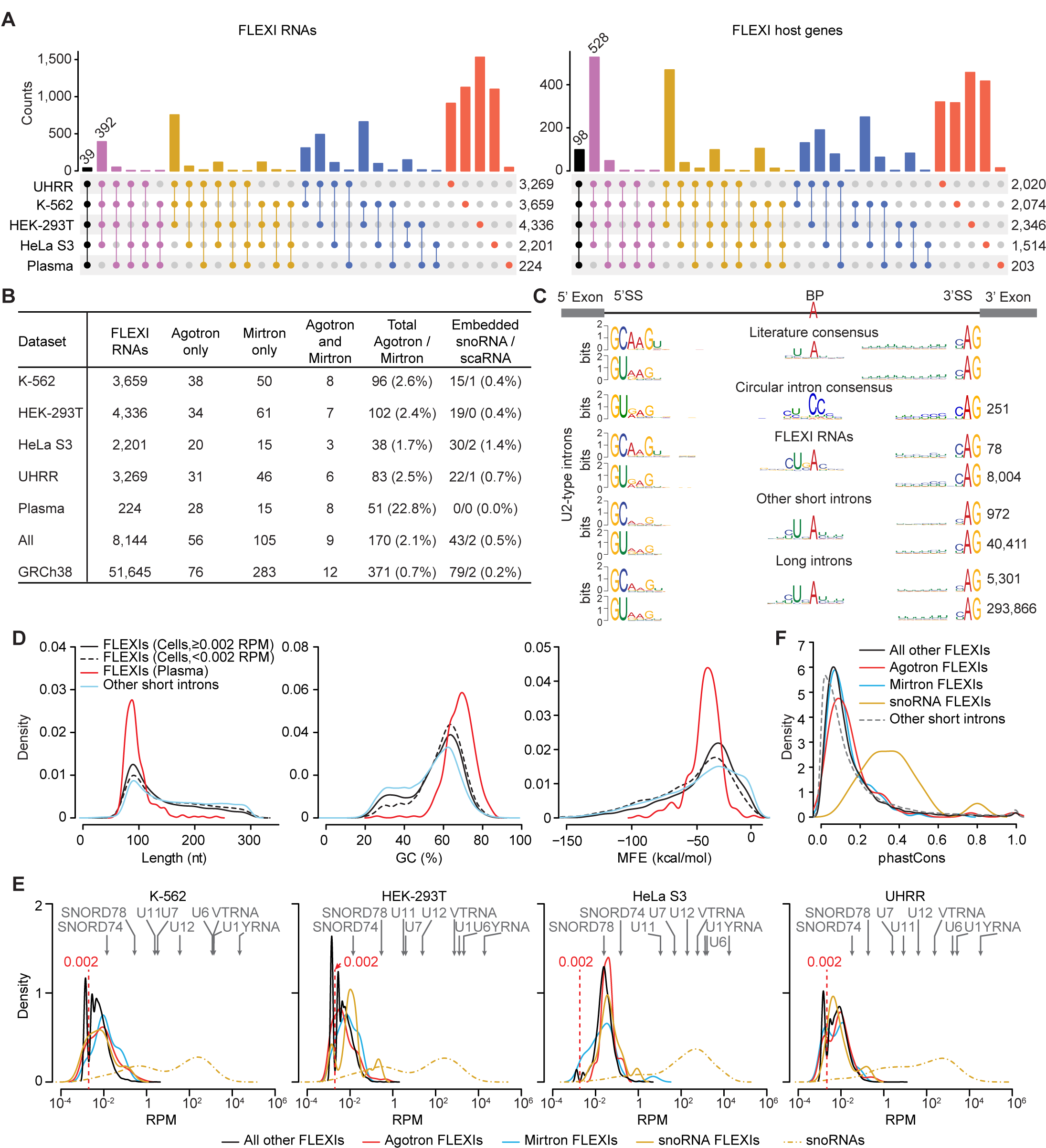
Characteristics of FLEXIs in human cells and plasma. (A) UpSet plots of FLEXI RNAs and their host genes identified by TGIRT-seq of ribo-depleted, unfragmented RNA preparations from the indicated human cell lines and plasma. Different FLEXIs from the same host gene were aggregated into one entry for that gene. (B) Numbers and percentages of different FLEXIs in datasets for each sample type corresponding to an annotated agotron, a pre-miRNA of annotated mirtron, or encoding an annotated snoRNA including subsets of small Cajal bodyspecific snoRNAs (scaRNAs). “All” indicates the total number of different FLEXIs in each cate-gory in this group of samples, and GRCh38 indicates the number of annotated short introns (≤300 nt) in each category in the GRCh38 human genome reference sequence. (C) 5’- and 3’-splice site (5’SS and 3’SS, respectively) and branch-point (BP) consensus sequences of FLEXIs corresponding to GC-AG and GU-AG U2-type spliceosomal introns in a merged dataset for the 4 cellular RNA samples compared to the literature consensus sequences for human U2-type spliceosomal introns (29, 30) and other Ensemble GRCh38-annotated U2-type short (≤300 nt) and long (>300 nt) introns. The number of introns with each consensus sequence is indicated to the right. (D) Density distribution plots of length, GC content, and minimum free energy (MFE) for the most stable RNA secondary structure predicted by RNAfold for more abundant (RPM≥0.002, solid black line) and less abundant (RPM <0.002, dashed black line) FLEXI RNAs in a merged dataset for the 4 cellular RNA samples compared to other GRCh38-annotated short introns (≤300 nt, blue line) in same merged dataset and FLEXIs in plasma (red line). (E) Density distribution plots of the abundance of different categories of FLEXIs in different cellular RNA samples color coded as indicated below the plots. The abundance distribution of annotated snoRNAs (dashed yellow line) and the abundance of annotated sncRNAs (gray arrows above the plots) in the same samples are shown for comparison. The vertical dashed red line in each plot separate low abundance (<0.002 RPM) from higher abundance (≥0.002 RPM) FLEXIs. (F) Density distribution plots of phastCons scores for different categories of FLEXIs in the cellular and plasma RNA samples compared to those of other annotated short introns (≤300 nt). PhastCons scores were calculated by the average score across all intron bases from multiple sequences alignment of 27 primates, including humans, in addition to mouse, dog, and armadillo.

To identify FLEXIs in the whole-cell RNA preparations, we compiled the coordinates of all short introns (≤300 nt) in Ensembl GRCh38 Release 93 annotations into a BED file and searched for intersections in TGIRT-seq datasets for each of the 4 cellular RNA samples. For each cellular RNA sample, the searched TGIRT-seq dataset included merged technical replicates totaling 666 to 768 million mapped paired-end reads, supplemented by datasets for previously published biological replicates to assess reproducibility (Table S1). As done previously for FLEXIs in plasma, we identified FLEXI by continuous intron reads that began and ended within 3 nt of annotated splice sites (20). This definition enabled us to include full-length intron reads with small numbers of non-templated nucleotides added by cellular enzymes to the 3’ end of RNAs (27) or by the TGIRT enzyme to the 3’ end of completed cDNAs (21, 22). The 300-nt size cutoff for more detailed characterization of FLEXIs was based on the length distribution of FLEXIs found previously in human plasma (20) and encompassed 85-92% of the FLEXIs detected in the 4 cellular RNA samples, including the most abundant FLEXIs (Fig. S3A). By using the same read mapping pipeline and mapping reads to an updated human genome reference sequence (GRCh38 Release 93), we identified a total 224 FLEXIs in the previously analyzed plasma samples from healthy individuals (20).

By using these approaches, we identified 8,144 different FLEXIs in the initially analyzed cellular and plasma RNA data originating from 3,743 different protein-coding genes, lncRNA genes, or pseudogenes (collectively denoted FLEXI host genes; Supplementary file). The distribution of FLEXIs and their host genes between the different cellular RNA samples and plasma is shown in UpSet plots in Fig. 1A. Fifty-six of the FLEXIs identified in the 4 cellular RNA samples and plasma corresponded to an annotated agotron and 105 corresponded to a pre-miRNAs of an annotated mirtron (Fig. 1B) (16, 19). Notably, the proportion of FLEXIs corresponding to annotated agotron or mirtron pre-miRNAs was higher in plasma (22.8%) than in cells (1.7-2.6%; p <1×10^-15^ by Fisher’s exact test for all cell types tested; Fig. 1B), suggesting preferential cellular export and/or greater resistance of these classes of FLEXIs to plasma RNases (20). Forty-three of the FLEXIs in the cellular RNA samples encoded a snoRNA, all which were also detected as processed mature snoRNAs in the same samples (Fig. 1B).

In Integrated Genomics Viewer (IGV) alignments of reads mapping to protein-coding genes, cellular FLEXIs stood as blocks of continuous intron reads that began and ended within 3 nt of annotated splice sites (Fig. 2A). The IGV alignments for 4I_JUP, an annotated agotron (19), provided an example of an excised intron RNA for which a high proportion (94%) of the reads satisfied the conservative definition of a FLEXI in whole-cell RNAs from HeLa S3 cells, but had 3’ truncations in the other cellular RNA samples that resulted in only small proportions of intron reads in those samples (2-3%) being counted as FLEXIs (Fig. 2A, top left). FLEXI reads showed no indication of impediments at or near a branch-point nucleotide as might be expected for a lariat RNA (Fig. 2A and B).

**Figure 2.**
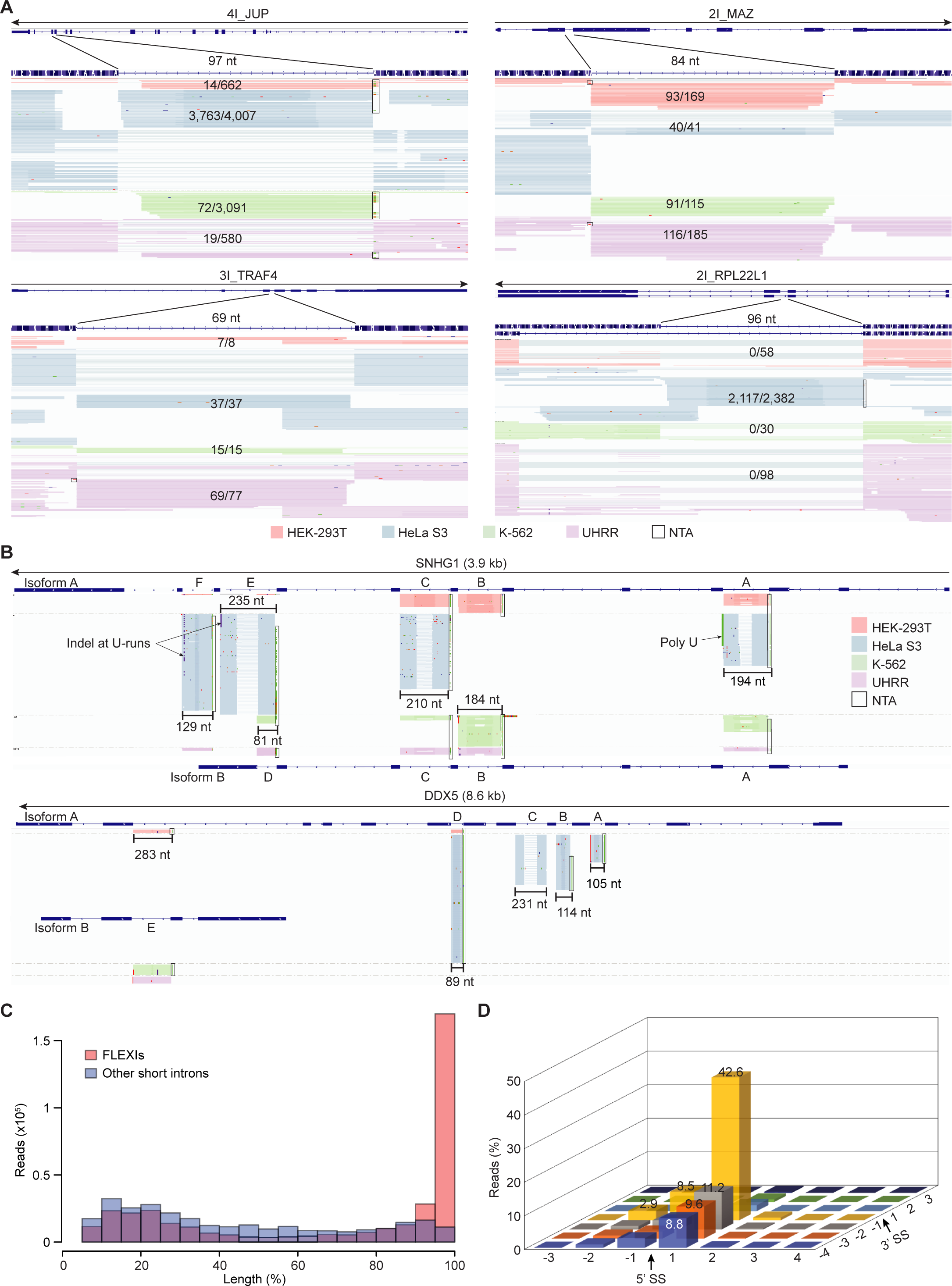
FLEXI RNAs are detected by TGIRT-seq as continuous full-length, end-to-end reads. (A) Integrative Genomics Viewer (IGV) screenshots showing read alignments for FLEXI RNAs in different cellular RNA samples. FLEXIs are named at the top with an arrow indicating 5’ to 3’ orientation and the tracks below showing gene annotations (exons, thick bars; introns, thin lines) followed by alignments of FLEXI reads in different cellular RNA samples. The length of the intron that is the focus of each panel is shown above the read alignments and the fraction of FLEXI reads (continuous reads that began and ended within 3 nt of the annotated 5’- and 3’-splice sites) of that intron is shown within the read alignments. (B) IGV screenshots showing examples of cell-type specific differences in abundance of FLEXI RNAs from the same host gene due to differences in splicing efficiency, alternative splicing, and/or differential stability of FLEXI RNAs. Gene maps for different RNA isoforms generated by alternative splicing of FLEXI RNAs are shown at the bottom with introns from the 5’ to 3’ RNA end named A to F. Reads mapping to exons or non-FLEXI introns were omitted for clarity. In panels A and B, reads for different cellular RNA samples are color coded as shown in the Figure and were down sampled to a maximum of 100. Mismatched nucleotides in boxes at the 5’ end of RNA reads due to non-templated nucleotide addition (NTA) to the 3’ end of cDNAs during TGIRT-seq library preparation. Two FLEXIs in panel B had indels at short runs of 3’ U residues, likely reflecting a previously reported tendency for TGIRT enzyme slippage at such locations (Fig. 2B) (67). (C) Histogram of the length distribution of reads mapping within introns identified as FLEXIs (red) compared to those mapping within other Ensemble GRCh38-annotated short introns (≤300 nt; blue) in a merged dataset for the 4 cellular RNA samples. Percent length was calculated from TGIRT-seq read spans in the indicated intervals normalized to the length of the corresponding intron. FLEXIs encoding embedded snoRNAs/scaRNAs were excluded to avoid contributions from fully or partially processed sno/scaRNAs. (D) Three-dimensional bar graph showing the percentage of FLEXI reads in a merged dataset for the K-562, HEK-293T, HeLa S3, and UHRR cellular RNA samples ending at different positions around intron-exon junctions.

Most FLEXIs (8,082 of 8,144) identified in the 4 cellular RNA samples and plasma had sequence characteristics of major U2-type spliceosomal introns (8,004 with GU-AG splice sites and 78 with GC-AG splice sites); 36 had sequence characteristics of minor U12-type spliceosomal introns (34 with GU-AG and 2 with AU-AC splice sites); and 26 had non-canonical splice sites (*e.g.,* AU-AG and AU-AU; Fig. 1C and S4) (28, 29). The splice-site and branch-point (BP) consensus sequences of the identified FLEXIs were similar to the literature consensus sequences of canonical human introns (28, 30) as well as those of other short and long introns in the same TGIRT-seq datasets, with no indication of enrichment for a CC branch-point sequence characteristic of the major class of trimmed lariat introns (Fig. 1C for U2-type FLEXIs and Fig. S4A for U12-type FLEXIs) (10, 12, 30). None of the identified FLEXIs corresponded to a previously identified trimmed lariat intron (10, 12); was present in databases of circular RNAs generated by backsplicing (circRNADb, http://reprod.njmu.edu.cn/circrnadb; circBase, http://www.circbase.org); or was identified by CIRI2 (31) as having back-spliced circular RNA junctions in TGIRT-seq datasets of chemically fragmented UHRR (32). RT-qPCR assays for six relatively abundant FLEXIs confirmed that they were sensitive to 5’- and 3’-exonuclease digestion, whereas a synthetic FLEXI RNA circularized *in vitro* was resistant to digestion by these nucleases (Fig. S5A and B, Supplementary file).

A histogram of the length distribution of all reads that mapped to FLEXIs in the 4 cellular RNA samples showed that the predominant bin corresponded to 90-100% intron length (Fig. 2C, pink bars). By contrast, the reads that mapped to other short introns (≤300 nt) annotated in GRCh38 corresponded to heterogeneously sized RNA fragments distributed across the length of the intron, as expected for rapid degradation of canonical excised intron RNAs (Fig. 2C, blue bars). Most of the FLEXI reads began and ended directly at the splice sites (87% for the 5’-splice site and 58% for the 3’-splice site), with <10% extending a short distance into a flanking exon (Fig. 2D for all FLEXIs and Fig. S3B for FLEXIs in each cellular RNA samples).

Density distribution plots showed that the FLEXIs identified in cells and plasma differed from those of other short intron in being skewed toward higher GC content and more stable predicted secondary structures (*i.e*., lower MFEs for the most stable secondary structure predicted by RNA fold; p≤0.001 for both by Wilcoxon signed-rank test; Fig. 1D). These differences were most pronounced for the FLEXIs found in plasma, which were a relatively homogeneous subset with peaks at 90-nt length, 70% GC content, and -40 kcal/mol MFE (Fig. 1D), suggesting preferential export or greater resistance to plasma nucleases of shorter, more stably structured FLEXIs. Similar length distributions were found for different categories of FLEXIs in each of the 4 cellular RNA samples, except those with a binding site for DICER (see below), which showed a secondary peak at ∼90% intron length, likely corresponding to DICER cleavage products or intermediates (Fig. S3C).

Collectively, the above findings show that human cells contain large numbers of short and in most cases structured FLEXIs, only a small subset of which correspond to annotated agotrons or mirtron pre-miRNAs (16, 19). Although hydrolytic splicing, which would directly produce an excised linear intron RNA, cannot be excluded, the largely canonical branch-point sequences in FLEXIs suggest that most if not all were excised as lariat RNAs and debranched after splicing, as found previously for mirtron pre-miRNAs and agotrons (17–19).

### Abundance of FLEXI RNAs

Density plots of the relative abundance of different FLEXIs in the cellular RNA samples showed separate peaks for lower and higher abundance FLEXIs (<0.002 RPM and ≥0.002 to 0.1 reads per million (RPM) with a tail extending to 6.9 RPM; Fig. 1E). The latter overlapped the lower end of the abundance distribution for snoRNAs (dashed yellow line; Fig. 1E). Lower and higher abundance FLEXIs (dashed and solid black lines, respectively) had similar characteristics (GC contents and MFEs) that differed from those of other short introns (Fig. 1D), suggesting that low abundance FLEXIs were not simply intron RNA turnover intermediates. In HeLa S3 cells, the peak of more abundant FLEXIs was predominant and agotrons comprised a higher proportion of more abundant FLEXIs than in the other cellular RNA samples (Fig. 1E). Relatively abundant FLEXIs included 3,363 detected at ≥0.01 RPM in one or more of the cellular RNA samples.

To estimate copy number per cell values of FLEXI RNAs, we used sncRNAs of known cellular abundance in the TGIRT-seq datasets to produce a linear regression model for the relationship between sequence-bias corrected log_10_ RPM values and log_10_-transformed sncRNA copy number per cell values from the literature (21) (Fig. S6A; Table S2). This method indicated that moderately abundant FLEXIs (0.01-0.99 RPM) were present at 15 to 239 copies per cell and that the most abundant FLEXIs (≥1 RPM) were present at 363 to 1,863 copies per cell. Based on the linear regression model, FLEXIs corresponding to annotated agotrons were present at up to 1,335 copies per cell; those corresponding to mirtron pre-miRNAs at up to 225 copies per cell; and those encoding snoRNAs at up to 265 copies per cell. Copy number per cell values determined by droplet digital PCR (ddPCR) for a sampling of FLEXIs were generally of the same order of magnitude as the TGIRT-seq quantitation, but varied with the choice of sncRNA standard, with the most abundant FLEXIs estimated to be present at 3,000 to 4,000 copies per cell, similar to that of U7 RNA, when it was used as the standard (Fig. S6B).

### Abundant FLEXI RNAs have cell-type specific expression patterns

Scatter plots identified subsets of relatively abundant FLEXIs (≥0.01 RPM) that were differentially expressed between two compared human cell lines (Fig. 3A, upper panels) and reproduci-bly detected at similar abundances in biological replicates of the same cell line (Fig. S7A). The cell-type specific expression patterns of FLEXI RNAs were confirmed by principal component analysis (PCA), t-SNE (33) and ZINB-WaVE (34), which showed clustering of FLEXI expression profiles in TGIRT-seq datasets for unfragmented UHRR and whole-cell RNAs from 5 different human cell lines (HEK-293T, HeLa S3, K-562, MDA-MB-231, and MCF7) for which we had additional biological and technical replicates from this or previous studies (Fig. S8A, Table S1). Notably, the differential expression patterns of FLEXIs were significantly more discriminatory between cell types than those for all RNAs that mapped to the same hosts genes (Fig. 3A, bottom panels), even after down sampling of host genes reads to match the sequencing depth of FLEXI reads (Fig. S7B upper and lower panels; p<0.0001 by Fisher’s r-to-z transformation in both cases). The clustering of cell-type specific expression patterns appeared to improve slightly when restricted to more abundant FLEXIs detected at ≥0.01 RPM, but not further with higher abundance cutoffs, likely reflecting that clustering was already biased toward higher abundance FLEXIs (Fig. S8B). The more discriminatory expression patterns of FLEXI RNAs, a desirable characteristic for cellular RNA biomarkers, likely reflects that their abundance can be impacted post-transcriptionally by different mechanisms, including alternative splicing.

**Figure 3.**
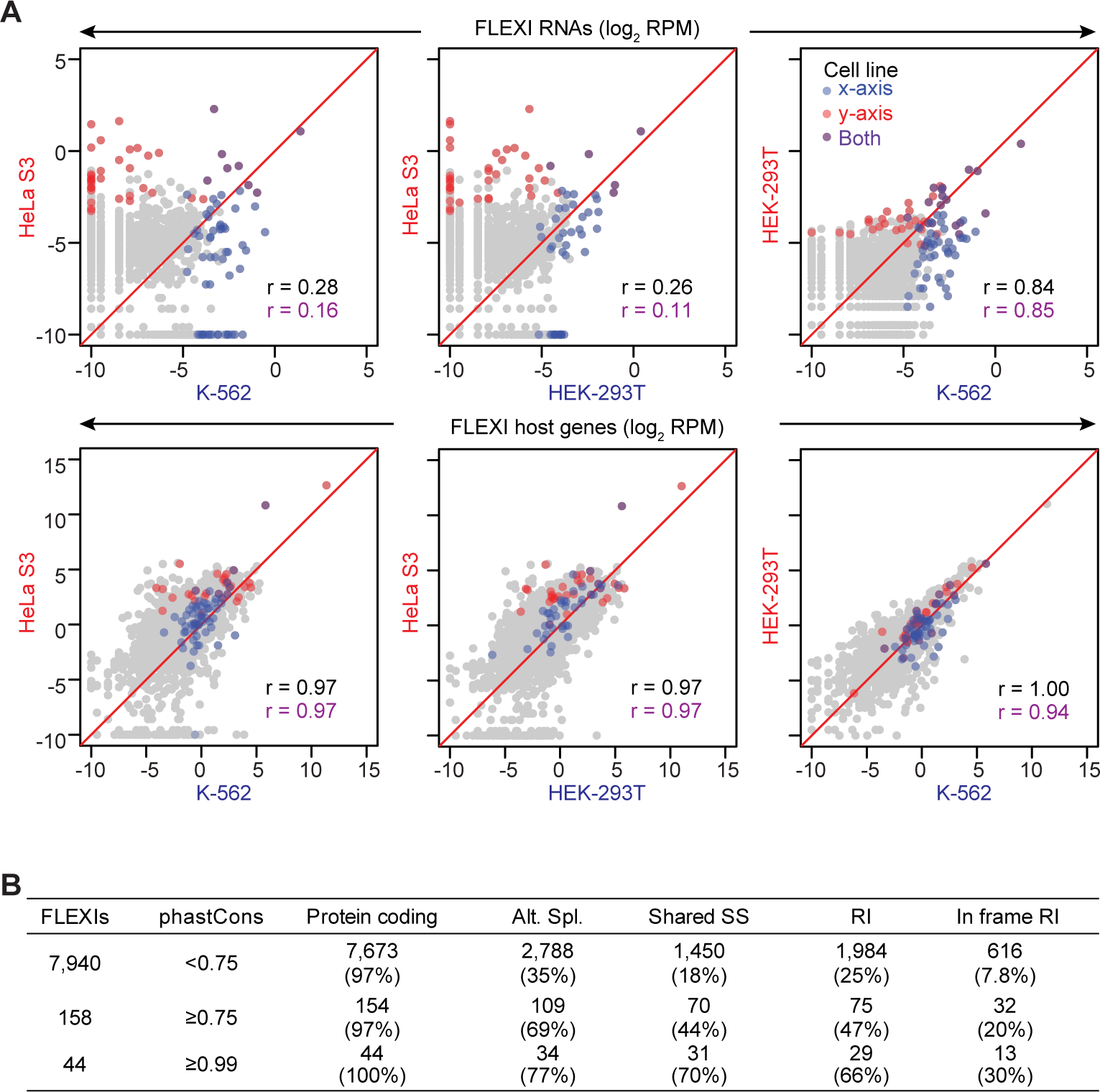
FLEXI RNA profiles are cell-type specific and alternative splicing characteristics of conserved FLEXIs. (A) Scatter plots showing pairwise comparisons of log_2_-transformed RPM of FLEXI RNA reads (top row) and reads that mapped to FLEXI host genes (bottom row) between different cellular RNA samples. Relatively abundant FLEXIs (≥0.01 RPM) and their host genes are colored red, blue, or purple depending upon whether they were reproducibly detected in all replicates of either or both compared cell types (color code in top right panel). Pearson correlation coefficients (*r*) for all detected FLEXIs and their host genes are shown in black, and those for relatively abundant color-coded FLEXIs and their host genes (red and blue) are shown in purple. (B) Characteristics of conserved FLEXIs in a merged dataset for the K-562, HEK-293T, HeLa S3, and UHRR cellular RNA samples. FLEXIs were divided into three groups based on phastCons scores for comparisons (<0.75, ≥0.75, and ≥0.99). The number of FLEXIs in each group is shown to the left, and the numbers and percentages of those FLEXIs in different categories are shown to the right. Alt. Spl. indicates FLEXIs that were alternatively spliced to generate different protein isoforms based on Ensemble GRCh38 annotations. Shared SS indicates FLEXIs that share a 5’- or 3’-splice site with a longer intron (>300 nt). RI indicates FLEXIs that are retained in some annotated alternatively spliced transcripts, and In frame RI indicates alternatively spliced retained FLEXIs that have reading frames in frame with those of flanking exons.

### Evolutionary conservation of FLEXIs

PhastCons scores provide a measure of evolutionary conservation, with higher phastCons scores indicating a higher probability that a sequence was conserved across a range of organisms (35). Density plots of phastCons scores for FLEXIs calculated across 27 primates, including humans, plus mice, dogs, and armadillos, showed that most FLEXIs, including those corresponding to annotated agotrons and mirtron pre-miRNAs, were poorly conserved (peaks at 0.06-0.09 compared to 0.02 for other annotated short introns in GRCh38), but with tails extending to higher phastCons scores. FLEXIs encoding snoRNAs had higher phastCons scores than other FLEXIs, likely reflecting conservation of the encoded snoRNA (Fig. 1F, yellow line). The relatively low phastCons scores of most FLEXIs could reflect more recent acquisition and/or more rapid sequence divergence in the human lineage.

Although most FLEXIs were poorly conserved, 158 FLEXIs that did not encode a snoRNA, had phastCons scores ≥0.75, possibly reflecting an evolutionarily conserved, sequence-dependent function (Fig. 3B). Compared to those with lower phastCons scores, these more highly conserved FLEXIs were enriched in introns that were alternatively spliced (69% compared to 35%), shared a 5’- or 3’-splice-site with a longer intron (44% compared to 18%), corresponded to an alternatively spliced retained intron (47% compared to 25%), or corresponded to an alternatively spliced retained intron with an in-frame protein-coding sequence (20% compared to 7.8%; Fig. 3B). The proportion of FLEXIs with these characteristics was even higher for the 44 most highly conserved FLEXIs (phastCons scores ≥0.99; Fig. 3B). These findings suggest that in most cases the sequence conservation of FLEXIs with high phastCons scores reflects evolutionary constraints on the sequences of different protein isoforms produced by alternative splicing.

### FLEXI RNAs have binding sites for RBPs with diverse cellular functions

To explore possible biological functions of FLEXIs, we searched for RBPs that bind FLEXIs in ENCODE eCLIP datasets for K-562 and HepG2 cells (36) and in AGO1-4 and DICER PAR-CLIP datasets for HEK-293 cells (37, 38). More than half of the detected FLEXIs (4,505; 55%) contained a CLIP-seq-identified binding site for one or more of 126 different RBPs (Fig. S9A), with 53 of these RBPs (51 in the eCLIP datasets plus DICER and AGO1-4 in the PAR-CLIP datasets) binding ≥30 different FLEXIs (Fig. 4A).

**Figure 4.**
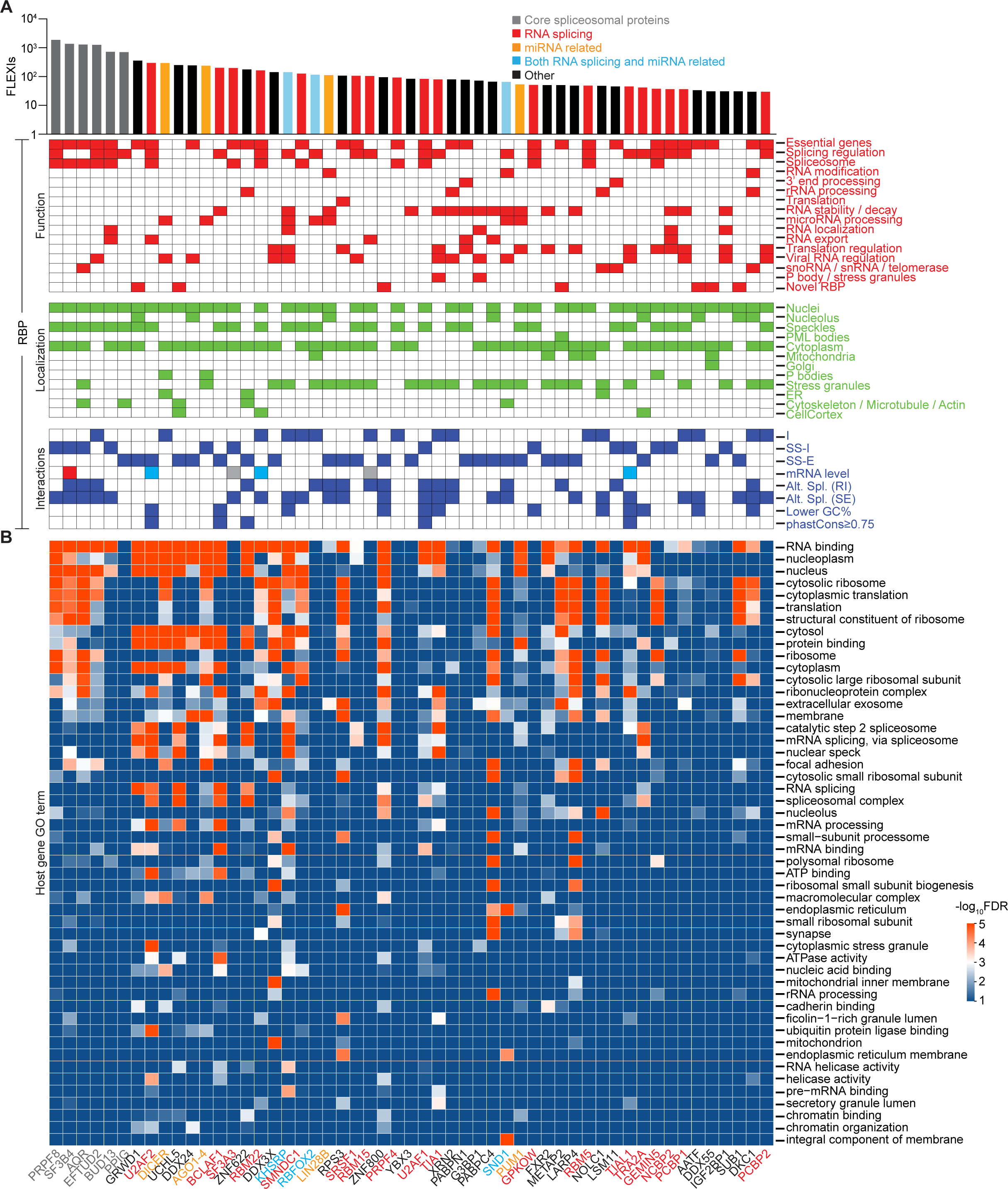
FLEXI interactions with RNA-binding proteins. (A) Bar graph showing the number of different FLEXI RNAs that have a CLIP-seq-identified binding site for each of the 53 RBPs that bind ≥30 different FLEXIs in a merged dataset for the K-562, HEK-293T, HeLa S3, and UHRR cellular RNA samples. An extended version of the bar graph for all 126 RBPs that have a CLIP-seq-identified binding site in a FLEXI RNA is shown in Fig. S9A. Bar graph RBPs are color coded by protein function as shown at the top. The grids below the bar graph show reported RBP Functions (red), Intracellular Localization (green), and FLEXI Interactions and characteristics. The Intracellular Localization Grid is based on a supplemental table in ref. (68), the mammalian RNA granule and stress granule protein databases (56, 69), and the UniProt database (https://www.uniprot.org). The Interaction Grid is based on analysis of PAR-CLIP datasets for AGO1-4 and DICER, ENCODE eCLIP and knockdown datasets for other RBPs, and the characteristics of FLEXIs identified in the cellular RNA datasets. RNA-binding sites: I, within the intron; SS-I and SS-E, across splice sites with the peak of density plots for binding sites localized within the intron or a flanking exon, respectively (Fig. S13). Changes in mRNA level based on ENCODE RBP-knockdown datasets: blue and red, significantly biased toward higher (log_2_FC>0) or lower (log_2_FC<0) mRNA levels, respectively, compared to all genes; gray, significantly changed mRNA levels, but not significantly biased in either direction (Fig. S14). Alternative Splicing (Alt. Spl.) based on ENCODE RBP-knockdown datasets compared to the control dataset (Fig. S15): RI, significant changes in retained introns; SE, significant changes in skipped exons. Lower GC%, RBPs whose binding sites were significantly enriched in FLEXIs having lower GC content than other FLEXIs (Fig. S12), phastCons scores ≥0.75, RBPs whose binding sites were significantly enriched in evolutionarily conserved FLEXIs with phastCons ≥0.75 (Fig. 6D). Changes in mRNA level and alternative splicing for AGO1-4 and DICER were not available in ENCODE. (B) Heatmap of GO terms enriched in host genes of FLEXI RNAs containing binding sites for different RBPs. GO enrichment analysis was performed with DAVID bioinformatics tools, using all FLEXI host genes as the background. The color scale shown to the right was based on -log_10_-transformed false discovery rate (FDR).

The 53 RBPs that bind ≥30 different FLEXI RNAs included 6 core spliceosomal proteins (PRPF8, SF3B4, AQR, EFTUD2, BUD13, and PPIG) and other proteins that function in RNA splicing; DICER, AGO1-4, and other proteins that have miRNA-related functions; and a surprising number of proteins that function in other cellular processes, including transcriptional regulation, chromatin assembly and disassembly, cellular growth regulation, and stress responses (Fig. 4A; protein functions described in Table S3). Examination of sequence reads in the ENCODE eCLIP datasets showed that all of the RBPs that bind ≥30 different FLEXIs were cross-linked to reads corresponding to ≥95% of the length of the identified FLEXI, with 32 showing a ≥2-fold enrichment of such reads compared to those in control antibody datasets (Fig. S10 and Supplementary file). Analysis of ENCODE and other published datasets found that many of the 53 RBPs that bind ≥30 FLEXIs were bound at pre-mRNA splice sites with subsets impacting alternative splicing of the bound FLEXI or adjacent exons (referred to below as proximate alternative splicing) and/or FLEXI host gene mRNA levels, when knocked down by shRNAs or siRNAs (Fig. 4A, RBP Interactions Grid). Notably all of the FLEXI-binding proteins in Fig. 4A were found in published datasets to be present in the cytoplasm, nucleolus, or cellular condensates (PML bodies, stress granules, or nuclear speckles) in addition to or instead of the nucleus (Fig. 4A, RBP Localization Grid based on ENCODE immunofluorescence data and published findings for AGO1-4 and Dicer (39–43)). The host genes encoding FLEXIs with binding sites for the 53 RBPs that bind ≥30 FLEXIs had a wide range of enriched Gene Ontology (GO) terms for nuclear and cytoplasmic functions (Fig. 4B).

Most of the 53 proteins that bind ≥30 different FLEXIs also had annotated binding sites in hundreds of other short introns and thousands of long introns, respectively (Fig. S9B and C). Scatter plots identified 7 RBPs whose binding sites were relatively abundant in FLEXIs (≥1% of the total) and significantly enriched in FLEXIs compared to both other short and long introns in a merged dataset for the 4 cellular samples (adjusted p≤0.05; calculated by Fisher’s exact test and adjusted by the Benjamini-Hochberg procedure; Fig. 5). These included 4 core spliceosomal proteins (PRPF8, SF3B4, EFTUD2, and BUD13), the miRNA-related proteins AGO1-4, DICER, and RBM22, a spliceosome component that functions in nuclear-cytoplasmic trafficking of ALU-7 and hSlu7 proteins under stress conditions (44) (Fig. 5A and B). Compared to long introns, FLEXIs were also significantly enriched in binding sites for DDX24, a DEAD-box protein that negatively regulates RIG-I and RIG-I-like receptors by sequestering cytosolic activator RNAs (45), and 3 proteins that interact with cytoplasmically localized back-spliced circular exon RNAs: GRWD1, a histone-binding protein that functions in ribosome assembly and regulates chromatin dynamics; UCHL5 (ubiquitin carboxy-terminal hydrolase) (46); and SF3A3 (Splicing Factor 3a subunit 3), whose interaction with a circSCAP activates TP53 (47) (Fig. 5B). The scatter plots also identified RBPs whose binding sites were over-represented in other short or long introns compared to FLEXIs, with larger numbers and greater differences for RBPs with binding sites in long introns (Fig. 5A and B).

**Figure 5.**
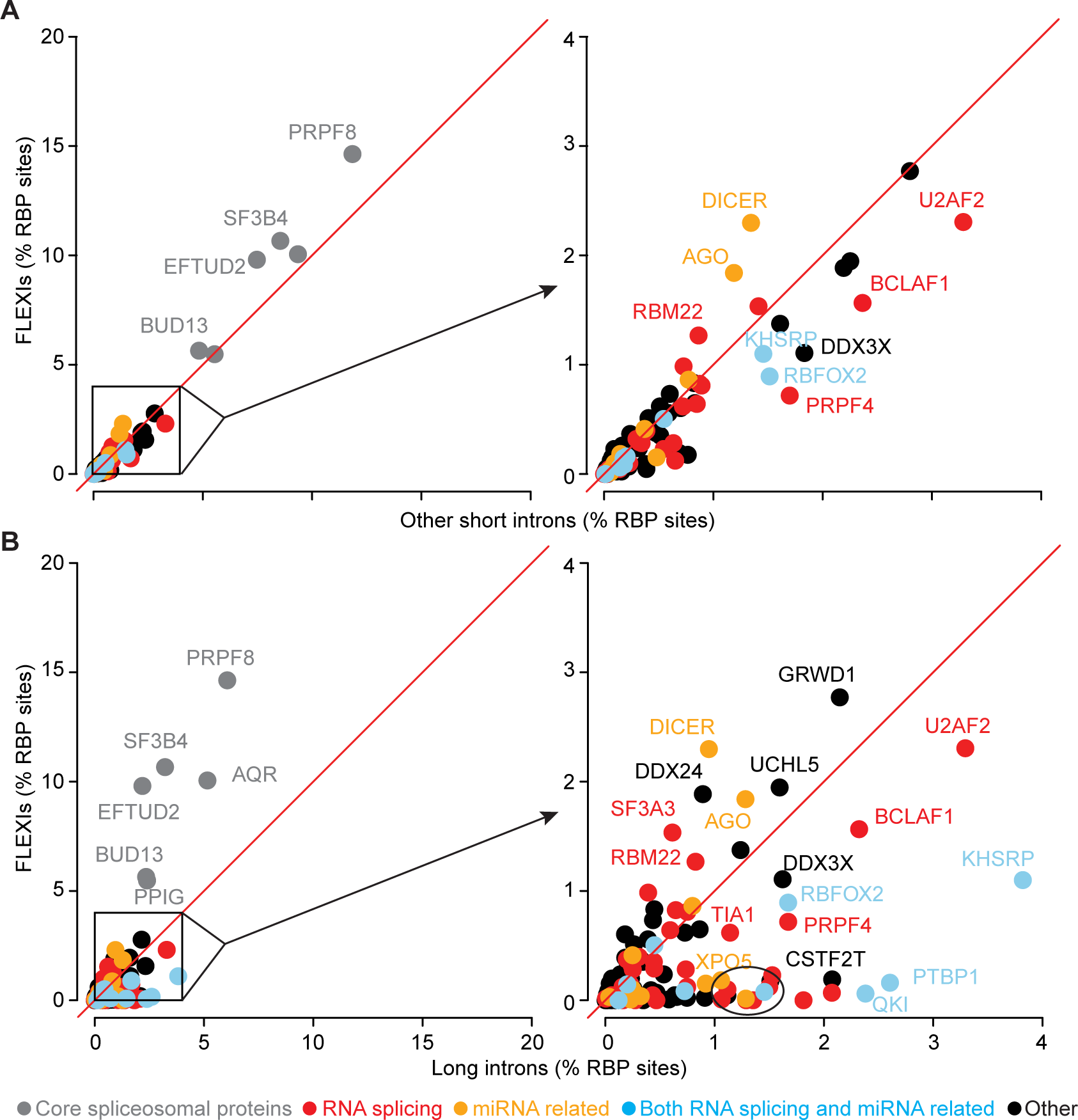
Relative abundance of annotated RBP-binding sites in FLEXIs compared to other short (≤300 nt) or long (>300 nt) introns. RBPs whose binding sites constituted ≥1% of the total RBP-binding sites and were significantly different between the compared groups (adjusted p≤0.05; calculated by Fisher’s exact test and adjusted by the Benjamini-Hochberg procedure) are indicated by the name of the protein and color coded by protein function as indicated at the bottom of the Figure. For each panel, the box within the scatter plot on the left delineates a region that is expanded in the scatter plot on the right. The circle at the bottom right of the scatterplot in panel B encompasses 12 additional RBPs (FAM120A, GTF2F1, HNRNPC, HNRNPK, HNRNPL, HNRNPM, ILF3, MATR3, NONO, PCBP2, SUGP2, and TARDBP) whose binding sites met the abundance and significance criteria for enrichment in long introns compared to FLEXIs but were too numerous to name in the Figure.

### Identification of FLEXIs enriched in binding sites for non-spliceosomal RBPs

Although FLEXIs as a whole were enriched in CLIP-seq-identified binding sites for 6 core spliceosomal proteins compared to other RBPs (Fig. 5A and B), scatter plots readily identified subsets of FLEXIs that were significantly enriched in binding sites for other subsets of RBPs (≥2% of RBP-binding sites; adjusted p≤0.05 calculated by Fisher’s exact test and adjusted by the Benjamini-Hochberg procedure; Fig. 6). These included: (i and ii) FLEXIs annotated as agotrons (n=61) or pre-miRNAs of annotated mirtrons (n=110), which were significantly enriched in binding sites for AGO1-4 and DICER (Fig. 6A and B); (iii) snoRNA-encoding FLEXIs (n=43), which were significantly enriched in binding sites for snoRNA-related proteins DKC1 (Dyskerin) and NOLC1 (Nucleolar and Coiled Body Phosphoprotein 1, alias Nopp140) (48, 49); SMNDC1 (Survival of Motor Neuron Domain Containing 1, alias SPF30), better known for binding snRNAs; and AATF (Apoptosis Antagonizing Transcription Factor, as well as several other proteins (Fig. 6C); and (iv) highly conserved, FLEXIs (phastCons scores ≥ 0.75, n=158), which were significantly enriched in binding sites for splicing factors U2AF1 and U2AF2, the splicing regulator TIAL1, ZNF622, which functions in cytoplasmic maturation of 60S ribosomal subunits (50), and the transcription factor BCLAF1 (Fig. 6D). The FLEXIs in these 4 subsets differed from FLEXIs as a whole in length, GC content, and/or MFE for the most stable secondary structure predicted by RNAfold (Fig. 6A-D, right hand panels). AATF, which we found enriched in subsets of FLEXIs encoding snoRNAs, was found previously to bind to both DKC1 mRNA and snoRNAs, potentially linking the ribosome biosynthesis function of snoRNAs to the regulation of cell proliferation and apoptosis by suppressing TP53 activity (51), and BCLAF1, which we found significantly enriched in conserved FLEXIs, was previously suggested to link RNA splicing to transcriptional and other cell regulatory pathways (52, 53).

**Figure 6.**
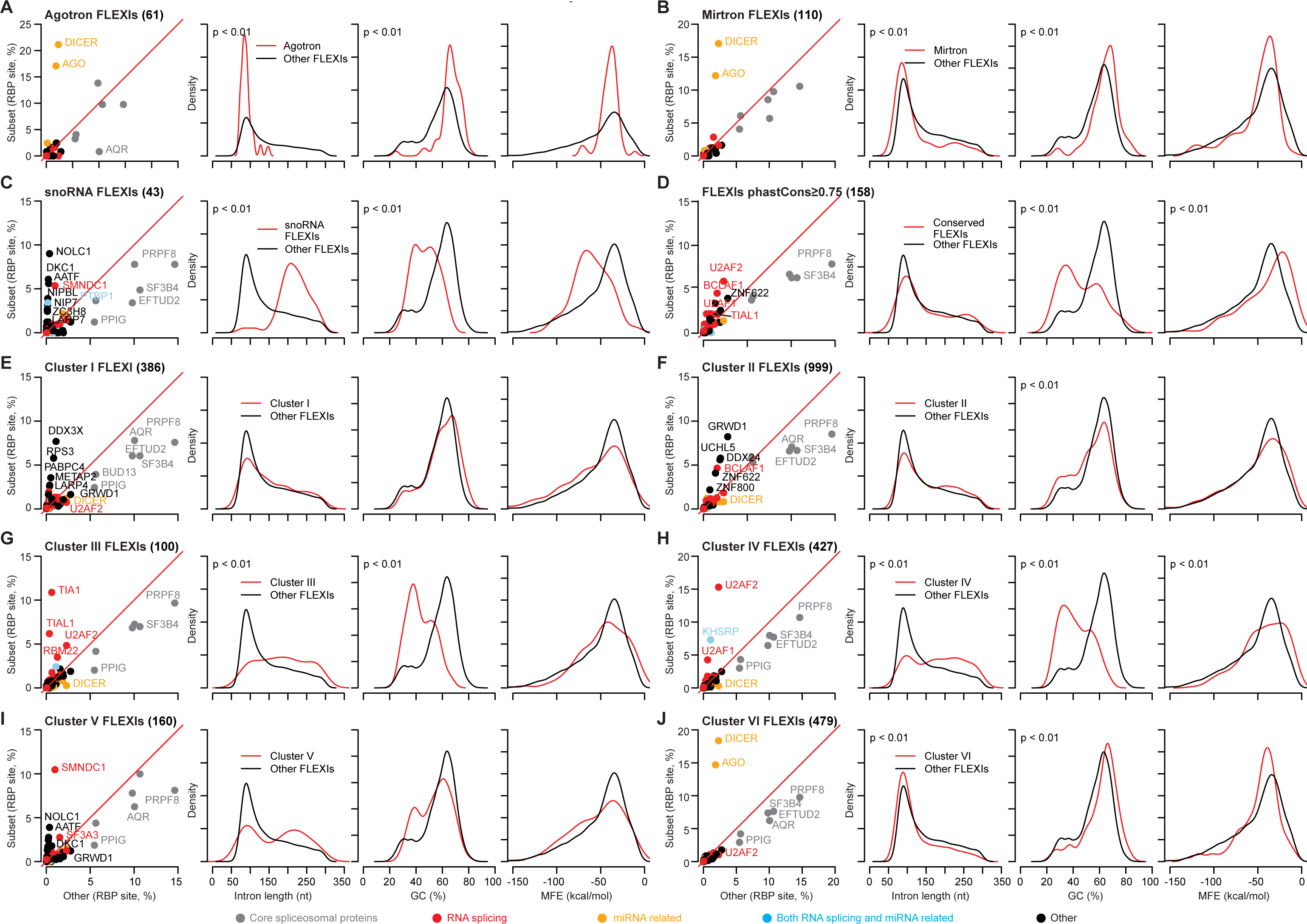
Identification and characteristics of subsets of FLEXIs that were enriched in binding sites for different subsets of RBPs. Different subsets of FLEXIs are named at the top left with the number of FLEXIs in that subset indicated in parentheses. The leftmost panel shows a scatter plot for CLIP-seq-identified binding sites in the Subset of FLEXIs (y-axis) compared to all Other FLEXIs (x-axis) in a merged dataset for the K-562, HEK-293T, HeLa S3, and UHRR cellular RNA samples, and the right three panels show density plots comparing different characteristics of the subset FLEXIs to other FLEXIs in the same merged datasets. (A) FLEXIs annotated as agotrons; (B) FLEXIs that are pre-miRNAs of annotated mirtrons; (C) FLEXIs encoding an annotated snoRNA. Binding sites for AATF were found only in FLEXIs encoding C/D-box snoRNAs, while those for DKC1, a core H/ACA-box snoRNA-binding protein (70), were found in FLEXIs encoding both H/ACA- and C/D-box snoRNAs. (D) Conserved FLEXIs with phastCons score ≥0.75; (E-J) FLEXIs identified by hierarchical clustering as being enriched in binding sites for different subsets of RBPs (see Fig. 7). RBP-binding site annotations in the scatter plots were based on the ENCODE eCLIP datasets for 150 RBPs (68) and AGO1-4 and DICER PAR-CLIP datasets (37, 38). In the scatter plots, RBP-binding sites whose abundance was ≥2% of all RBP-binding sites and significantly different between the compared subset and all other FLEXIs (adjusted p≤0.05; calculated by Fisher’s exact test and adjusted by the Benjamini-Hochberg procedure) are labeled with the name of the RBP color coded by protein function as indicated at the bottom of the Figure. The density distribution plots compare the length, GC content, and MFE for the most stable secondary structure predicted by RNAfold for FLEXI RNAs comprising the subset (red) to those of all other detected FLEXIs (black). p-values are shown at the top left of those density plots in which the distribution for the subset of FLEXIs differed significantly from other FLEXIs (p<0.01 by Kolmogorov–Smirnov test and reproducible in ≥95% of 1,000 Monte-Carlo simulations; see Methods). Scatter plots and density plots for FLEXIs with binding sites for each RBP associated with these clusters are shown in Fig. S12.

### Computational identification of FLEXIs co-enriched in binding sites for different subsets of RBPs

Encouraged by the above findings, we used hierarchical clustering as described in Materials and Methods to computationally identify subsets of RBPs whose binding sites were co-enriched in different subsets of FLEXIs without *a priori* assumptions about the function of the protein or bound FLEXIs. This approach identified 6 clusters of RBPs whose binding sites were significantly co-enriched (relative abundance ≥2%, adjusted p≤0.05) in the same subsets of FLEXI in all or some of the 4 human cell lines (Fig. 7A and Fig. S11). Scatter plots confirmed the enrichment of binding sites for each subset of RBPs relative to those for core spliceosomal proteins in each cluster of FLEXIs (Fig. 6E-J), and scatter plots for each individual RBP in each cluster confirmed co-enrichment of binding sites for the other RBPs in the same cluster (Fig. S12).

**Figure 7.**
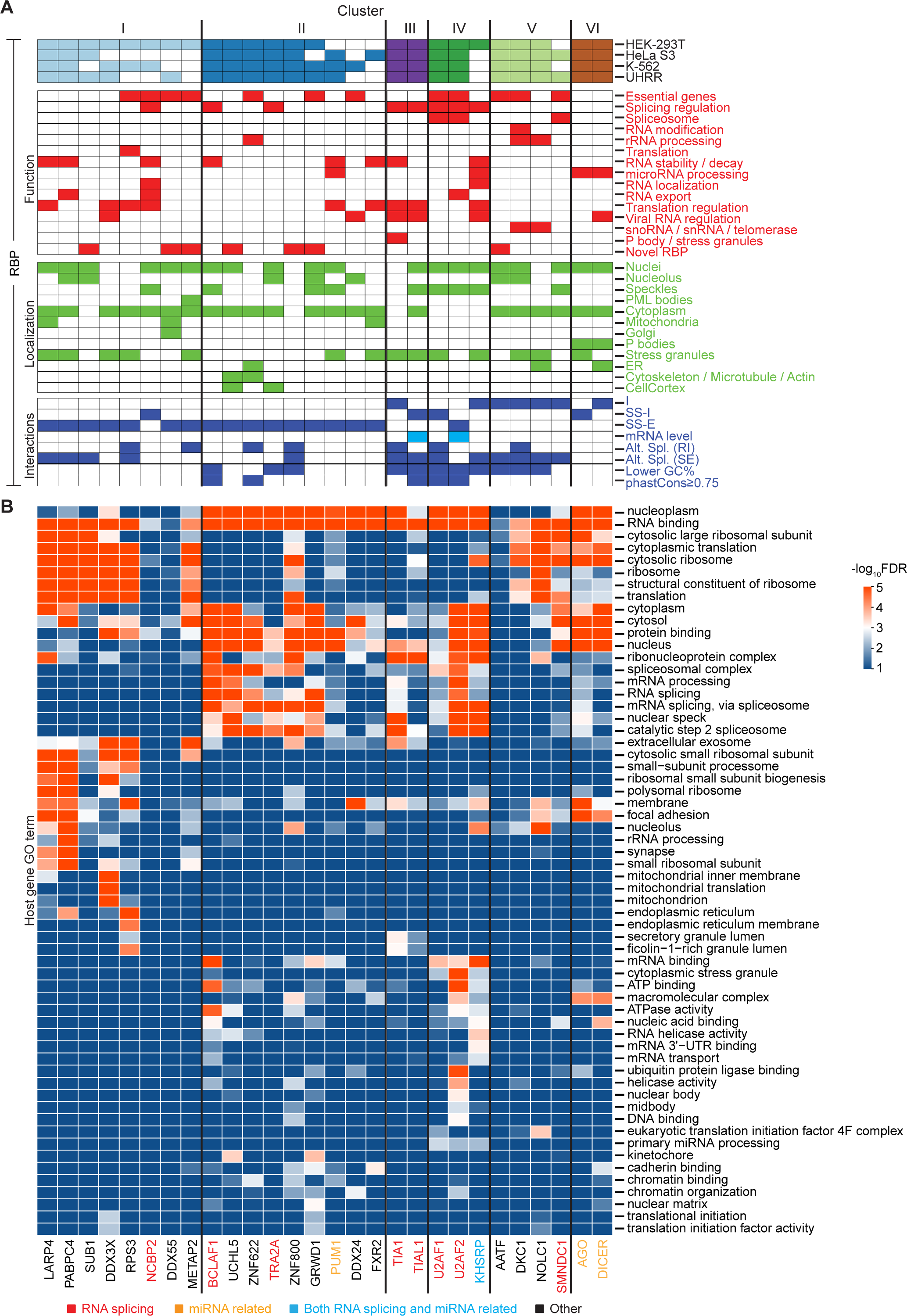
Identification of RBPs whose binding sites were co-enriched in clusters of FLEXI RNAs. (A) Hierarchical clustering of FLEXI RNAs based on patterns of overand underrepresented RBP-binding sites (see Methods and Fig. S11). Clusters I to VI are delineated at the top of panel, and the names of the RBPs associated with each cluster are shown at the bottom of panel B and were color coded by protein function as indicated at the bottom of the Figure. The RBPs that met the abundance and significance criteria for association with the clusters in HEK-293T, HeLa S3, K-562, and UHRR are indicated by filled boxes at the top. The grids below summarize RBP Functions, Localization, and Interactions with FLEXs based on data sources indicated in the legend of Fig. 4. (B) Heatmap of GO terms enriched in host genes of FLEXI RNAs containing binding sites for cluster-associated RBPs. GO enrichment analysis was performed with DAVID bioinformatics tools, using all FLEXI host genes as the background. The color scale to the right was based on -log_10_-transformed false discovery rate (FDR).

Clusters I and II had the largest numbers of FLEXIs and were the most revealing of novel RBP-FLEXI associations. Cluster I was comprised of 386 FLEXIs with two RBPs (LARP4 and PABPC4) that satisfied the relative abundance and significance criteria for association with the cluster in all 4 cell lines and 6 other RBPs (Transcription Regulator SUB1, DDX3X, RPS3, NCBP2, DDX55, and Methionine Aminopeptidase 2 (METAP2)) that satisfied these criteria in HEK-293T cells but not in some of the other cell lines (Fig. 7A). LARP4 and PABPC4 function in regulating the stability and translation of mRNAs (54, 55), and the other 6 proteins in this cluster either have mRNA synthesis or translation-related functions or could link these functions to other cellular regulatory processes (Table S3). Analysis of ENCODE eCLIP and RBP-knockdown datasets showed that all of the proteins in Cluster I bind at splice sites with knockdown of 5 of the 8 proteins impacting proximate alternative splicing of the bound FLEXI (retained intron (RI) or skipped adjacent exons (SE; Fig. 7A, RBP Interactions Grid; based on data in Figs. S13-15). All but one of these proteins (SUB1) were annotated as being present in the cytoplasm in addition to or instead of the nucleus, with 5 of the 8 proteins also annotated as being present in stress granules (Fig. 7A, RBP Localization Grid).

Cluster II was comprised of 999 FLEXIs with 5 proteins (BCLAF1, UCHL5, ZNF622, TRA2A, ZNF800) satisfying the abundance and significance criteria for association with this cluster in all 4 cell lines, two (GRWD1 and PUM1) satisfying these criteria in 3 of the 4 cell lines, and two others (DDX24 and FXR2) satisfying these criteria in only one of the cell lines (Fig. 7A). All of the proteins associated with this cluster function in regulation of transcription or RNA splicing (Table S3) and bind at splice sites, with knockdown of two of the proteins impacting proximate alternative splicing of the bound FLEXI (Fig. 7A and Fig. S13-15). All but one of these proteins were annotated as being localized in the cytoplasm in addition to or instead of the nucleus, the sole exception being ZNF800, whose cellular localization was not available in ENCODE.

Clusters III and IV were comprised of 100 and 427 FLEXIs, respectively, that were enriched in binding sites for the splicing regulators TIA1 and TIAL1 (Cluster III) or splicing factors U2AF1 and U2AF2 (Cluster IV), with splicing regulator KHSRP satisfying the abundance and significance criteria for association with Cluster IV in HEK-293T cells (Fig. 7A). Three of these proteins (TIAL1, U2AF1, and U2AF2) bind at splice sites and the other two (TIA1 and KHSRP) bind within the introns, with knockdown of all 5 proteins impacting proximate alternative splicing of the bound FLEXI and/or mRNA levels of FLEXI host genes (Fig. 7A). Notably, in addition to or in one case instead of the nucleus, all 5 proteins in these clusters were annotated as being present in nuclear speckles and/or stress granules condensates (56, 57), possibly reflecting post-splicing destinations.

Cluster V was comprised of 160 FLEXIs that were enriched in binding sites for the 4 proteins that were identified in Figure 6C as binding snoRNA-encoding FLEXIs (DKC1, NOLC1, AATF, and SMNDC1), with the first 3 proteins satisfying the relative abundance and significance criteria for association with this cluster in all 4 of the cell lines and SMNDC1 satisfying these criteria in 2 of the cell lines (Fig. 7A). Only 43 of the 160 FLEXIs in this cluster encoded an annotated snoRNA. All 5 of the RBPs in this cluster bind within the intron with their knockdown affecting proximate alternative splicing of the bound FLEXI (Fig. 7A).

Finally, Cluster VI was comprised of 479 relatively GC-rich FLEXIs that were enriched in binding sites for AGO1-4 and DICER (Fig. 7A). Only 23 of the 250 FLEXIs in this cluster with a binding site for AGO1-4 corresponded to an annotated agotron (19), and only 44 of the 308 FLEXIs in this cluster with a binding site for DICER corresponded to a pre-miRNA of an annotated mirtron (58). The remaining FLEXIs bound by these proteins could be unannotated agotrons or mirtrons; could be cleaved by DICER to yield unannotated short regulatory RNAs; or could reflect binding of AGO1-4 and DICER to structured introns that were derived from or could evolve into agotrons or mirtrons, an evolutionary scenario suggested previously for mirtrons (18). Analysis of PAR-CLIP datasets showed that DICER binding sites were localized within FLEXIs, while those for AGO1-4 were at splice sites (Fig. 7), potentially enabling regulation of host gene RNA splicing (59, 60). Although we continue to focus below on the subsets of RBPs that met the criteria for hierarchical clustering in Clusters I-VI, heat maps showed that numerous other RBPs had overlapping binding sites in FLEXIs belonging to these clusters, suggesting more complex regulatory networks to explore in subsequent studies (Fig. S16).

### Clusters of FLEXI RNAs identified by co-enrichment of RBP-binding sites originate from host genes with related biological functions

To assess if the host genes encoding FLEXIs in each of the 6 identified clusters might be functionally related, we analyzed GO terms for host genes encoding FLEXIs that bind each RBP in each cluster using DAVID bioinformatic tools and displayed the results as a heat map (Fig. 7B). The heat map showed that enriched GO terms for FLEXI host genes within each of the 6 clusters were more congruent with each other than those for the full set of host genes for FLEXIs with binding sites for 53 RBPs that bind ≥30 FLEXIs in Fig. 5B. The host genes for FLEXIs bound by AGO1-4 and DICER (Cluster VI) were enriched in GO terms for cytoplasmic translation and focal adhesion, while those for FLEXIs in Clusters II, III, and IV were enriched in GO terms for RNA splicing, mRNA processing and nuclear speckles, with some host genes for FLEXIs in Clusters I, II, IV, and VI enriched in a GO term for protein binding. The host genes for FLEXIs bound by the snoRNA-related proteins in Cluster V have functions related to cytosolic ribosomes and cytoplasmic translation, with the host genes of FLEXIs bound by NOLC1 also involved in focal adhesion.

Notably, enriched GO terms for the host genes of FLEXIs in Cluster I included cytosolic ribosome and cytoplasmic translation, which were also enriched in host genes of the snoRNA-related FLEXIs in Cluster V (Fig. 7B). Consistent with these GO terms, we found that the Cluster I and V FLEXIs were enriched in host genes encoding cytoplasmic ribosomal proteins (9.6% and 11.2% of FLEXI host genes, respectively, compared to 1.6% of all FLEXI host genes; p<1×10^-11^ by Fisher’s exact test; incorporated into more detailed analysis below). These findings raised the possibility that Clusters I and V RBPs function in parallel pathways for regulating host genes involved in ribosome synthesis and translation. Collectively, the GO term analysis indicated that the host genes for FLEXIs that bind the same subset of RBPs are functionally related and might be coordinately regulated by these RBPs.

### Host genes for FLEXIs are enriched in other short and long introns with binding sites for same subsets of RBPs

All of the RBPs identified as having annotated binding sites in FLEXIs also had annotated binding sites in other short and long introns (Fig. S9). UpSet plots showed that most FLEXIs containing binding sites for the RBPs in Clusters I to VI were enriched in host genes that also contained other short or long introns with binding sites for the same RBPs, while the majority of other short introns and the vast majority of long introns were present in host genes that lacked FLEXIs with binding sites for Cluster I to VI RBPs (Fig. 8A). Consistent with these findings, hierarchical clustering, done similarly to that for FLEXIs using 10 subsets of 2,000 randomly selected other short or long introns, largely recapitulated Clusters I to VI for FLEXIs with some simulations finding additional RBPs associated with these clusters, as well as additional clusters of other short and long introns co-enriched in binding sites for other RBPs (Clusters VII to X; Fig. 8A). The host genes for Clusters I and V identified by hierarchical clustering of other short or long introns included those with binding sites for the same RBPs identified by hierarchical clustering for FLEXIs and were likewise enriched in GO terms for cytoplasmic translation and cytosolic ribosome (Fig. 9, left and right heat maps, respectively, compare with Fig. 7B). Analysis of CLIP-seq and knockdown datasets showed that in most cases the Cluster I to VI RBPs bound other short and long introns at similar locations at splice sites or within the introns as they did for FLEXIs, with knockdown of some RBPs impacting proximate alternative splicing of the bound intron or host gene mRNA levels (Fig. S13-15). Collectively, these findings indicate that different subsets of host genes are enriched not only in FLEXIs but also other short introns and long introns that bind similar subsets of RBPs, potentially reinforcing coordinate regulation by these RBPs.

**Figure 8.**
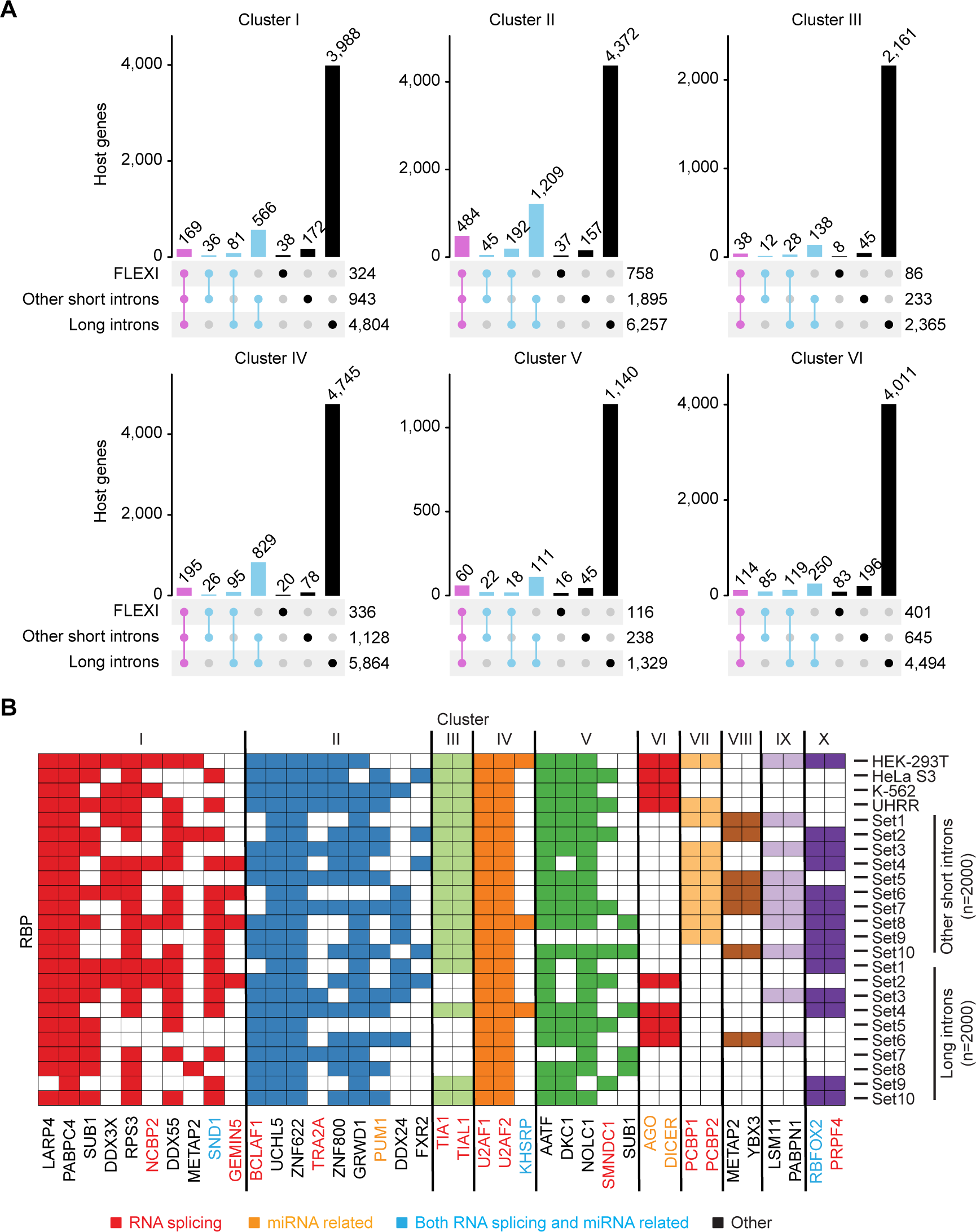
Host genes for FLEXIs in Clusters I to VI are enriched in other short and long introns that have binding sites for the same subsets of RBPs. (A) UpSet plots showing the host genes of FLEXIs, other short introns (≤300 nt) and long (>300 nt) introns containing CLIP-seq-identified binding sites for the subsets of RBPs associated with FLEXIs in Clusters I to VI. (B) Hierarchical clustering for co-enrichment of RBP-binding sites in other short and long introns was done as in Fig. 7 for 10 subsets of 2,000 randomly selected other short introns or long introns (Set1-10). Hierarchical clustering recapitulated Clusters I to VI found for FLEXIs with additional RBPs associated with these clusters as well as additional clusters (VII to X) found for some subsets of other short or long introns. The subsets of FLEXIs in HEK-293T, HeLa S3, K-562, and UHRR are shown at the top for comparison. Names of RBPs are color coded by protein function as indicated at the bottom of the Figure.

**Figure 9.**
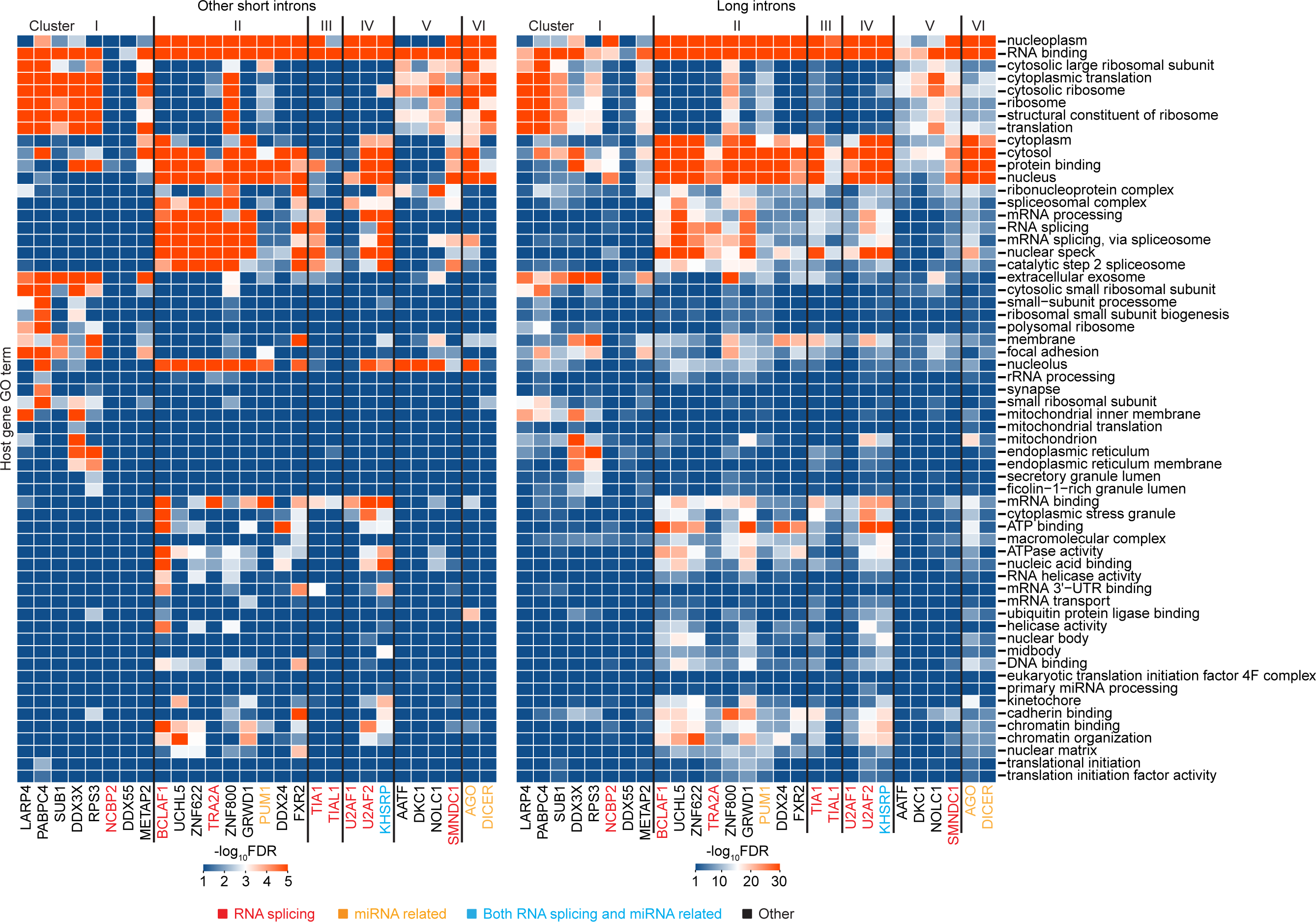
GO terms for host genes of other short and long introns that contain binding sites for RBPs in Clusters I to VI are similar to those for FLEXIs in Clusters I to VI. GO enrichment analysis was done using DAVID bioinformatics tools for host genes containing other short (left) or long (right) introns. The color scales at the bottom of each heat map were based on - log_10_-transformed FDR. RBP names are color coded by protein function as indicated at the bottom of the Figure.

### The intracellular localization of FLEXIs correlates with that of their CLIP-seq-identified RNA-binding proteins

Most of the RBPs associated with Clusters I and II were annotated as being present in both the cytoplasm and nucleus, with a number of these proteins having known cytoplasmic functions (Fig. 7). To investigate the intracellular location of FLEXIs with binding sites for different RBPs, we used TGIRT-seq to analyze RNAs in nuclear and cytoplasmic fractions prepared from 4 different cell lines (K-562, HeLa S3, MDA-MB-231, and MCF7; Fig. 10). The breast cancer cell lines MDA-MB-231 and MCF7 were included in this analysis to assess the generality of conclusions based on the initially analyzed cellular RNA samples, as well as the potential utility of FLEXIs as breast cancer biomarkers. TGIRT-seq of MDA-MB-231 and MCF7 whole-cell RNAs identified 1,288 additional FLEXIs that were not detected in any of the previously analyzed cellular RNA samples (501 in MCF7, 599 in MDA-MB-231, and 188 in both MCF7 and MDA-MB-231), including cell-type specific FLEXIs from oncogenes and tumor suppressor genes (Fig. S17 and Supplementary file). The clean separation of nuclear and cytoplasmic fractions from the 4 cell lines was confirmed by scatter plots comparing different RNA biotypes in each fraction from each cell type (Fig. S18). PCA analysis and a sample distance heat map showed that the TGIRT-seq datasets clustered mainly by cell type, with nuclear, cytoplasmic, and whole-cell RNAs from each cell type forming separate but closely spaced subclusters (Fig. S19).

**Figure 10.**
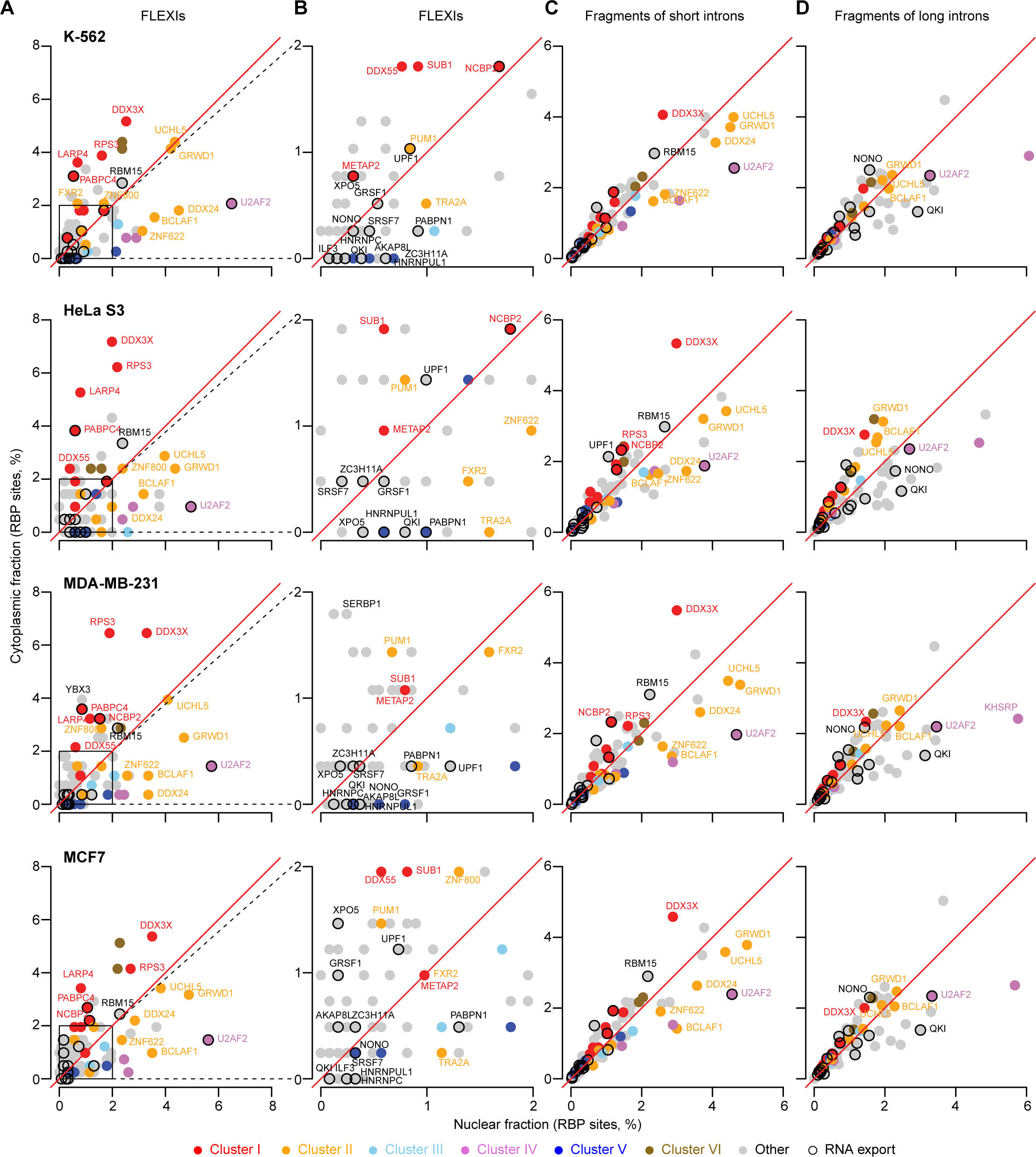
The intracellular localization of FLEXI RNAs correlates with that of their CLIP-seq-identified RNA-binding proteins. Scatter plots comparing the relative abundance of annotated RBP-binding sites in FLEXIs and fragments of other short introns and long intron RNAs in nuclear or cytoplasmic fraction from K-562, HeLa S3, MDA-MB-231, and MCF7 cells. (A and B) Scatter plots for FLEXIs. The box within the scatter plot in panels A delineates a region that is expanded in panels B. (C and D) Scatter plots for other short and long introns, respectively. Each point in the scatter plots represents the relative abundance of RBP-binding sites in intron RNAs in the nuclear and cytoplasmic with points for different categories of RBPs color coded as shown below the plots. In panels A and B, only FLEXIs that were differentially enriched in the nuclear or cytoplasmic fractions (FC>1.5 by DESeq2 analysis; denoted nuclear and cytoplasmic FLEXIs) were used for the analysis. RBPs that belong to Clusters I and II or function in nuclear RNA export are indicated by named points in the scatter plots. Also named are other RBPs whose abundance was ≥2% of all RBP-binding sites and were significantly enriched in intron RNAs in the nuclear or cytoplasmic fractions (adjusted p≤0.05; calculated by Fisher’s exact test and adjusted by the Benjamini-Hochberg procedure).

Fig. 10A and B show scatter plots for each of the 4 cell lines comparing the relative abundance of CLIP-seq identified RBP-binding sites in FLEXIs enriched in the nuclear or cytoplasmic fractions (fold change (FC)>1.5 via DESeq2 analysis, denoted nuclear or cytoplasmic FLEXIs, respectively). The percentages and numbers of different FLEXIs with a binding site for each RBP associated with Clusters I to VI in the nuclear, cytoplasmic or both fractions are shown in Fig. 11. Both Figures show that cytoplasmic FLEXIs in all 4 cell lines were enriched in binding sites for each of the 8 proteins associated with Cluster I (name labeled red points in scat-terplots). These included not only proteins with known cytoplasmic functions (LARP4, PABPC4, RPS3, and METAP2), but also DDX3X, which functions in nuclear RNA export (59–61), as well as NCBP2 (nuclear cap binding protein 2) and SUB1 (a transcriptional regulator). Cytoplasmic FLEXIs in all 4 cell lines also appeared enriched in those with binding sites for AGO1-4 and DICER, consistent with the functions of agotrons and mirtrons (Figs. 10A and B, unlabeled brown points, and Fig. 11).

**Figure 11.**
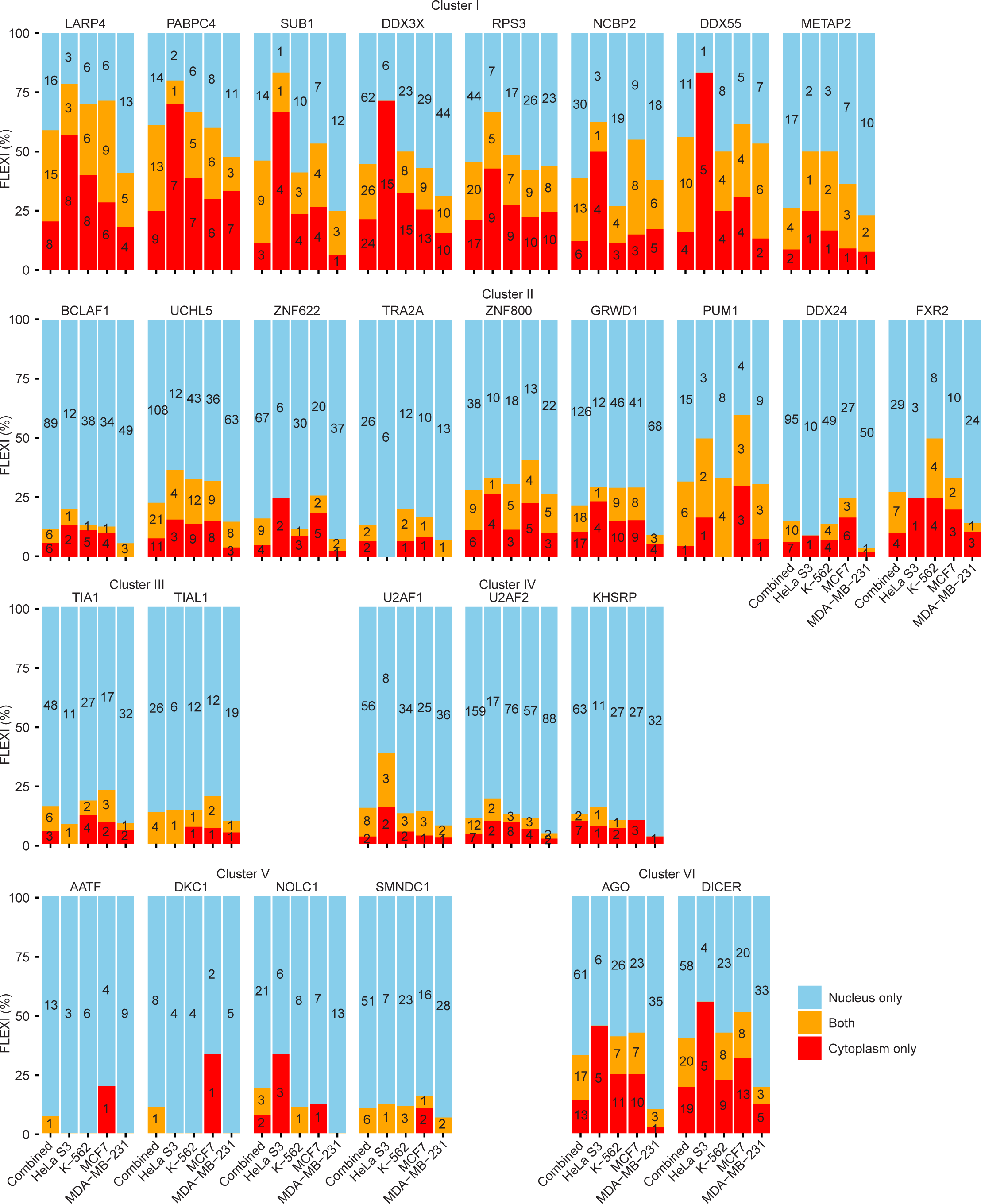
Distribution of FLEXI RNAs detected in nuclear, cytoplasmic or both fractions. Stacked bar graphs show the percentages of FLEXI RNAs containing binding sites for RBPs associated with Clusters I to VI in nuclear, cytoplasmic, or both fractions (color coded as shown bottom right) in combined (left bar graph) or separate (right bar graphs) in TGIRT-seq datasets for 4 different cell lines. The number of FLEXIs identified in either or both fractions is indicated within each section of the stacked bar graph.

The FLEXIs with binding sites for Cluster II-associated proteins (name labeled orange points in scatter plots) had more variable distributions. Those with binding sites for PUM1, FXR2, and ZNF800 had the highest degree of enrichment in cytoplasmic FLEXIs; those with binding sites for UCHL5 and GRWD1 were more equally distributed between cytoplasmic and nuclear FLEXIs; and those with binding sites for BCLAF1, ZNF622, DDX24, and TRA2A had the highest degree of enrichment in nuclear FLEXIs (Figs. 10 and 11). FLEXIs with binding sites for RBPs associated with Clusters III, IV, and V (light blue, purple and dark blue points in scatter plots) were enriched nuclear FLEXIs (Figs. 10 and 11).

The scatter plots also showed that cytoplasmic FLEXIs were enriched in binding sites for 9 other proteins with functions consistent with their presence in the cytoplasm (36, 62) in one or more of the cell lines (Fig. 10A and B; Table S3). These include 7 proteins that function in nuclear RNA export (AKAP8L, GRSF1, RBM15, SRSF7, UPF1, XPO5, and ZC3H11A) plus SERBP1, which stabilizes inactive ribosomes and is involved in PML-body formation, and YBX3, a non-specific RNA-binding protein and homolog of YBX1, which functions in exosomal packaging of sncRNAs (26).

In contrast to FLEXIs, other short or long introns were comprised of RNA fragments. Scatter plots comparing the distribution of binding sites in RNA fragments of other short and long introns in the nuclear and cytoplasmic showed at most moderate enrichment in one or the other fraction in most cases (Fig. 10C and D). A notable exception was DDX3X, whose binding site were significantly enriched in RNA fragments of other short introns in the cytoplasmic fraction from all 4 cells lines (Fig. 10C). RNA fragments of long introns in the cytoplasmic fraction were significantly enriched in binding sites for the RNA export protein QKI in all 4 cell lines, while RNA fragments with binding sites for the RNA export protein NONO were significantly enriched in the nuclear fraction from 3 of the 4 cell lines (Fig. 10D).

Stacked bar graphs based on CLIP-seq datasets showed that the majority of individual FLEXIs in Clusters I to VI have a binding site for a single RBP associated with that cluster (gray bars), with smaller percentages having overlapping binding sites for two or more RBPs associated with the same cluster (red and blue bars, respectively) and low percentages having nonoverlapping binding sites for 2 or more RBPs associated with the same cluster (orange and purple, respectively; Fig. 12A). A precedent for this situation is the alternative splicing factor RBFOX2, which binds at some splice sites via a primary conserved sequence motif and at others by binding at secondary sites or by interacting with partner proteins bound at those sites (63). Consistent with the latter mechanism, the STRING protein-network database confirmed interactions between nearly all of the proteins associated with each of the hierarchical clusters (Fig. 12B).

**Figure 12.**
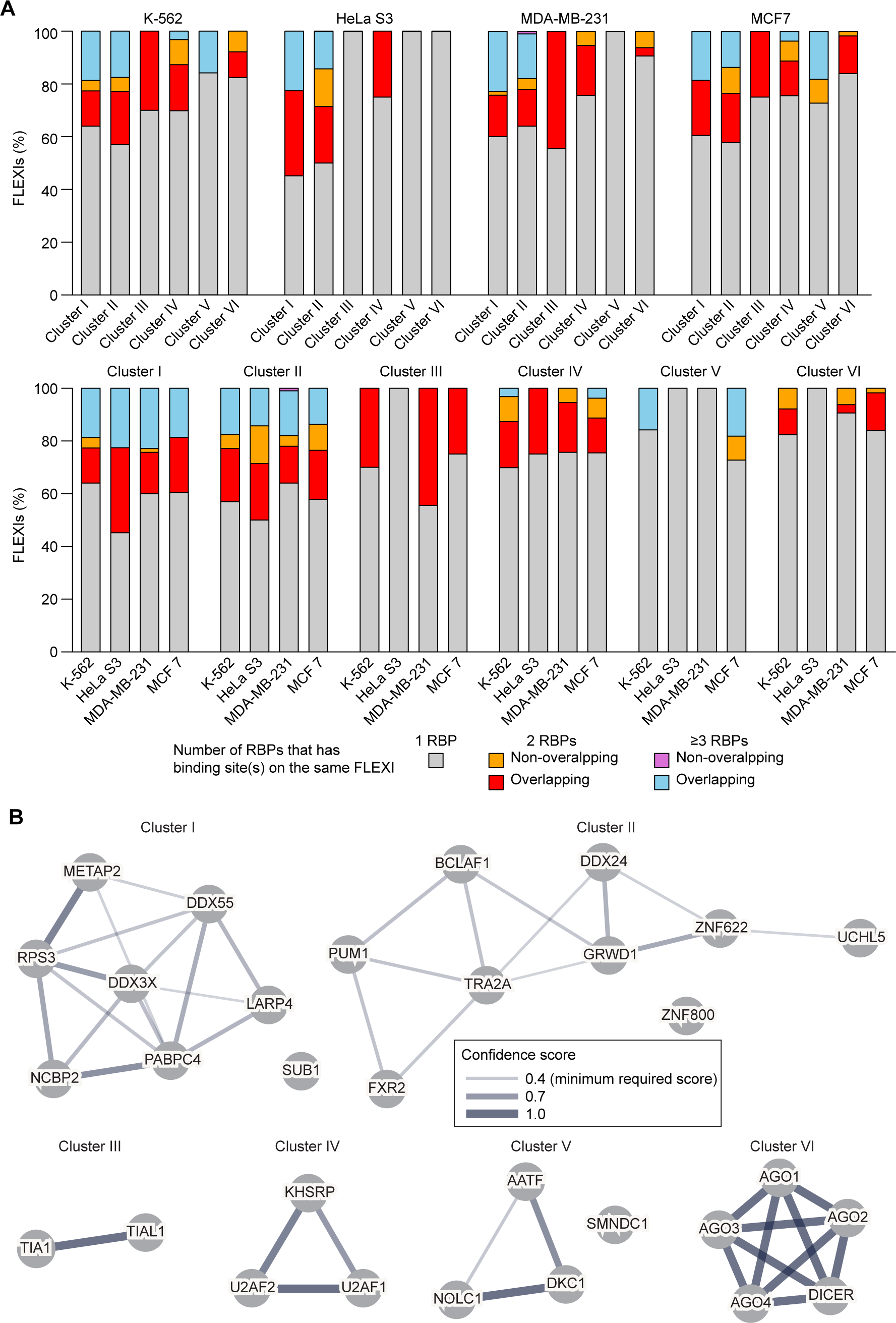
Numbers of CLIP-seq-identified binding sites for different RBPs in individual FLEXI RNAs belonging to Clusters I-VI. (A) Stacked bar graphs organized by cell type (top row) or cluster (bottom row) showing the percentage of FLEXI RNAs that have an annotated CLIP-seq-identified binding site for a single RBP, 2 RBPs, or ≥3 RBPs associated with the same cluster, with FLEXI RNAs having binding sites for ≥2 RBPs subdivided by whether or not the CLIP-seq-identified binding sites were overlapping or non-overlapping. (B) Protein-protein interactions of RBPs associated with Clusters I to VI based on protein-interaction network analysis using the STRING database (https://string-db.org/). Interacting RBPs are connected by lines whose thickness indicates the confidence level of the interaction score.

## Discussion

In total for all datasets in this study, we identified 9,429 different FLEXIs expressed from 4,224 host genes, representing ∼18% of the 51,645 short introns (≤300 nt) annotated in Ensembl GRCh38 Release 93. FLEXIs were skewed toward higher GC content and more stable predicted secondary structures than those of other short introns (Fig. 1D), likely contributing to the higher stability of FLEXIs in cells and plasma. Subsets of relatively abundant FLEXIs had more discriminatory cell-type specific expression patterns than other RNAs mapping to the same host gene in the same datasets (Fig. 3 and Fig. S7), a desirable feature for RNA biomarkers as discussed further below. The conserved branch-point A residues of FLEXIs (Fig. 1C) suggests that they splice via a lariat intermediate, but does not preclude that they can also splice via a hydrolytic mechanism, as found *in vivo* for certain group II intron RNAs, which also have a conserved branch-point A residue (64).

By searching CLIP-seq datasets, we identified 126 different RBPs that bind FLEXIs, including 53 RBPs that bind ≥30 different FLEXIs (Fig. 4A). These 53 RBPs included not only proteins that function in RNA splicing, but also AGO1-4 and DICER and other proteins that have miRNA-related functions, and a surprising number of proteins that function in other cellular processes, including transcription (AATF, BCLAF1, and SUB1), chromatin remodeling (GRWD1), translation (LARP4, METAP2, PABPC4, RPS3), ribosome maturation (ZNF622), and cellular growth regulation (DDX3X and IGF2BP1; Fig. 4A and Table S3). In addition to being present in the nucleus, all 53 RBPs that bind ≥30 different FLEXIs were also annotated in published datasets as being present in the cytoplasm and/or nuclear or cytoplasmic condensates (Fig. 4). Additionally, a number of RBPs that bind FLEXIs function in nuclear RNA export (BUD13, NCBP2, PABPC4, PABPN1, RBM15, and U2AF2 among the 53 RBPs that have annotated binding sites in ≥30 FLEXI RNAs in Fig. 4A, and AKAP8L, GRSF1, HNRNPA1, HNRNPC, HNRNPUL1, ILF3, KHDRBS1, NONO, QKI, SRSF7, UPF1, XPO5, ZC3H11A among the remaining RBPs that bind smaller numbers of FLEXIs in Fig. S9). Bound RBPs may contribute to the stability of FLEXIs in cells and plasma as well as shielding against cellular inflammatory responses to structured cytoplasmic RNAs.

By using hierarchical clustering, we identified 6 clusters of FLEXIs (Clusters I to VI) that were significantly co-enriched in binding sites for 6 clusters of RBPs, including regulatory proteins and RBPs whose knockdown affected alternative splicing and/or host gene mRNA levels (Fig. 7). We then found that other short and long introns formed hierarchical clusters similar to Clusters I to VI for FLEXIs and that the host genes for FLEXIs in Clusters I to VI were enriched in other short and long introns with binding sites for the same subsets of RBPs (Fig. 8). Collectively, these findings suggest that different subsets of functionally related host genes are coordinately regulated by different subsets of RBPs whose binding sites are enriched in different classes of introns within that gene set. Particularly suggestive was Cluster I, which included two proteins, LARP4 and PABPC4, that function in regulating the stability and translation of mRNAs (54, 55), along with additional proteins that have translation-related functions (NCBP2, METAP2, and RPS3), or that could link these functions to other cell regulatory pathways (SUB1, RPS3, DDX3X, and DDX55).

Notably, the host genes encoding the FLEXIs in Cluster I were enriched in GO terms similar to those for the snoRNA-related FLEXIs in Cluster V (Fig. 7B), suggesting that the FLEXIs and RBPs of Cluster I may function in parallel to snoRNAs in regulating ribosome synthesis (Fig. 7B). Along the same lines, Cluster II included 3 proteins (UCHL5, ZNF622, and TRA2A) that were distinguished from other proteins associated with the FLEXI-RBP clusters as being localized to the endoplasmic reticulum, cytoskeleton, and cell cortex, possibly reflecting a cytoskeleton or membrane-associated function. The RBPs associated with Clusters III and IV, which function in alternative splicing, were found localized not only in the nucleus but also in stress granules and nuclear speckles, the former possibly a mechanism for muting inflammatory responses to naked cytoplasmic RNAs, and the latter possibly a mechanism for storing alternative splicing factors for subsequent use (65). Although the physiological significance of CLIP-seq identified RNA-protein interactions requires further investigation, the specificity of the CLIP-seq-identified binding sites that formed the basis for our analysis is supported by the findings that host genes for FLEXIs with binding sites for RBPs in Clusters I to VI were co-enriched with other short and long introns with binding sites for the same RBPs (Fig. 8); that knockdown of subsets of RBPs in each cluster impacted alternative splicing and/or host gene mRNA levels; and that interactions between the RBPs in each cluster, suggested by the hierarchical clustering based on RNA-protein interactions, were confirmed independently by protein-interaction network analysis (Fig. 12B).

Lewis *et al.* (66) recently reported that 5 of the 8 RBPs that we identified as belonging to Cluster I (LARP4, PABPC4, DDX3X, RPS3, and SUB1) have eCLIP-identified binding sites enriched in mRNAs of nuclear-gene encoding mitochondrially localized proteins. They also found that LARP4-binding sites were enriched near the 3’ end of mRNAs encoding components of respiratory chain complexes and mitochondrial ribosomal proteins (66). Prompted by these findings, we found that FLEXIs with binding sites for LARP4 and other Cluster I RBPs were also enriched in host genes encoding mitochondrially localized proteins annotated in MitoCarta3.0 (8.3 to 25.5% of FLEXIs host genes compared to 7.5% of all FLEXI host genes and 5.7% of all GRCh38 annotated protein-coding genes; Fig. S20A and B). Extending these findings, we found that FLEXIs with binding sites for LARP4 and other Cluster I RBPs were also enriched in host genes encoding cytoplasmic ribosomal proteins (8.3 to 42.6%), consistent with enrichment for the GO terms cytoplasmic ribosome and cytoplasmic translation for FLEXI host genes in this cluster (Fig. 7A). Some but not all FLEXIs with binding sites for Cluster I RBPs were enriched in genes encoding mitochondrial ribosomal proteins (DDX3X, 6.8%; LARP4, PABPC4, RPS3, 1.9-2.2% compared to 0% for the other 4 RBPs), with similar trends holding for other short and long introns with binding sites for the same RBPs (Fig. S20A). Consistent with the findings of Lewis *et al.* (66), we found that binding sites for LARP4 and Cluster I proteins PABPC4, DDX55 and SUB1 were localized near the 3’ end of mRNAs (defined as UTRs + coding sequences (CDS)). Extending these findings, we found that binding sites for these 4 proteins were also localized to 3’-proximal introns, suggesting that they bind initially to unspliced pre-mRNAs and that their localization, possibly as a complex (Fig. 12B), might be dictated by proximity to 3’-poly(A) tails (Fig. S20C). METAP2- and RPS3-binding sites were more uniformly distributed across the length of the mRNAs, but remained concentrated in 3’-proximal introns, while DDX3X- and NCBP2-binding sites were localized near the 5’ end of mRNAs and in 5’-proximal introns, with DDX3X still showing a smaller peak for 3’-proximal introns (Fig. S20C). Considered together, these findings suggest that the RBPs that bind FLEXIs in Cluster I function in at least two overlapping regulatory pathways, one for cytoplasmic translation and cytosolic ribosomes parallel to that for snoRNAs, and the other for mitochondrial function, both potentially related to regulation of cell growth and proliferation.

Collectively, our findings suggest a model in which newly synthesized or excess RBPs that are not associated with their primary physiological substrates enter the nucleus where they bind FLEXIs and other short or long introns in pre-mRNAs from subsets of genes with related functions, contributing to the regulation of different gene sets. After splicing, the RBPs remain stably bound to FLEXIs and co-localize with them to the cytoplasm or other intracellular locations, where they may dissociate from FLEXIs to perform different cellular functions. As these RBPs include transcription factors and nuclear export proteins as well as RNA splicing factors, coregulation of gene sets by these RBPs could occur at multiple levels. After splicing, FLEXIs may serve simply as vehicles that stabilize RNA-binding proteins until they reach an appropriate location to perform their cellular functions or could be actively engaged in those functions. Although only 6 subsets of RBPs met the arbitrary significance criteria for co-enrichment in the same subsets of FLEXIs, a heat map of overlapping RBP-binding sites identified multiple additional RBPs whose binding sites overlapped those of RBPs within each cluster of FLEXIs (Fig. S16), possibly enabling cross talk between different gene sets. Collectively, these findings suggest overarching mechanisms for communicating and coordinating regulation of different cellular processes.

Finally, relatively abundant FLEXIs that have cell-type specific expression patterns may have immediate utility as cellular RNA biomarkers. Besides being linked to the expression of thousands of protein-coding and lncRNA genes, FLEXI RNA levels reflect differences in alternative splicing and intron RNA stability, potentially affording higher resolution of cellular differences than mRNAs transcribed from the same gene. Because most FLEXIs are predicted to fold into stable secondary structures, their initial identification may be best done by TGIRT-seq or other methods using related enzymes that yield full-length, end-to-end sequence reads of structured RNAs. Once identified, the validation of candidate FLEXI biomarkers and their routine monitoring would best be done by methods such as RT-qPCR, microarrays or other hybridization-based assays, or targeted RNA-seq that enable high-throughput quantitative read outs of panels of RNA biomarkers for different diseases.

### Data access

TGIRT-seq datasets have been deposited in the Sequence Read Archive (SRA) under accession numbers PRJNA648481 and PRJNA640428. A gene counts table, dataset metadata file, FLEXI metadata file, RBP annotation file, PCR-product sequencing files, and scripts used for data processing and plotting have been deposited in GitHub: https://github.com/reykeryao/FLEXI.

## Acknowledgements

We thank Marta Mastroianni for cell culture and RNA preparations, Shelby Winans Bretton for early research contributions, Camilo Molina Romero and Kavyaa Choudhary for help with validating amplicons for exonuclease and droplet digital PCR analysis, Blerta Xhemalce (University of Texas at Austin) for fresh stocks of MDA-MB-231 and MCF7 cells, Carla Van Den Berg (University of Texas at Austin) for comments on the manuscript and use of droplet digital PCR equipment, and Jim Manley (Columbia), Sandra Wolin (NIH/NCI), and Richard Maraia (NIH/NICHD) and for comments on the current and/or an earlier version of the manuscript. The authors acknowledge the University of Texas at Austin Genomic Sequencing and Analysis Facility (GSAF, Center for Biomedical Research Support, RRID#: SCR_021713, https://wikis.utexas.edu/display/GSAF/Home+Page) for providing RNA-seq services and the Texas Advanced Computing Center (TACC; http://www.tacc.utexas.edu) and Biomedical Research Computing Facility (BRCF; https://site.research.utexas.edu/cbrs/cores/cbb/computing-resource) for providing high performance computing resources. This work was supported by NIH grant R35 GM136216 to AML. MA was supported by NIH grant R01 GM040478.

---

**Figure S1.**
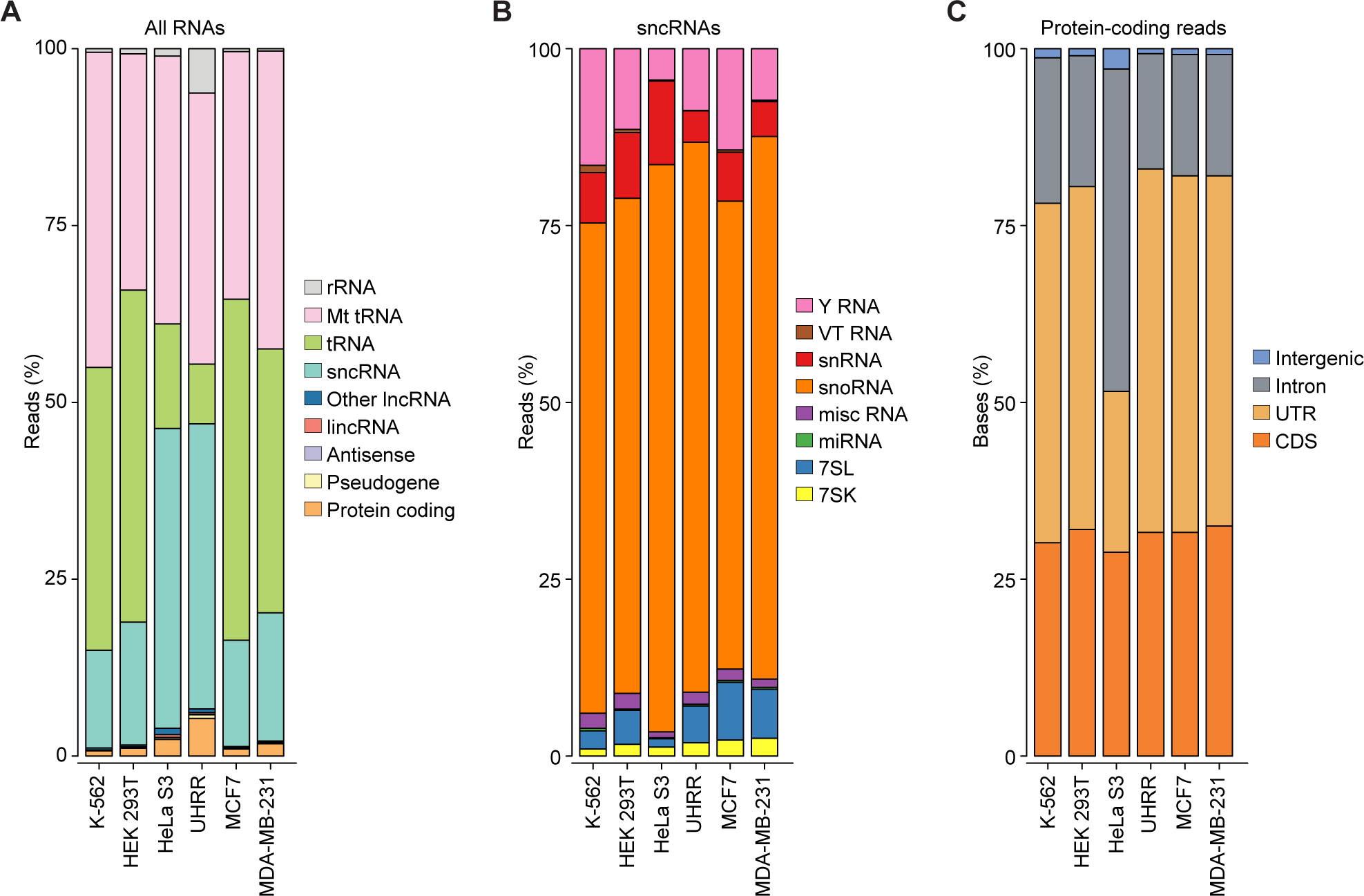
Classes of RNAs identified by TGIRT-seq of rRNA-depleted unfragmented cellular RNAs. Stacked bar graphs showing the percentages of sequencing reads or bases that mapped to different categories of Ensembl GRCh38 Release 93 annotated genomic features in combined technical replicates for the indicated cellular RNA samples. (A) Percentages of sequencing reads that mapped to different classes of cellular RNAs. rRNA includes cellular and mitochondrial (Mt) rRNAs, and protein-coding gene RNAs includes all transcripts of proteincoding genes from both the nuclear and mitochondrial genomes. (B) Percentages of sequencing reads that mapped to different sncRNAs. misc RNA: miscellaneous RNAs, such as ribozymes, small NF90-associated RNAs (snaRs), promoter-associated RNAs (pRNAs), and other ncRNAs that do not fall into other categories. (C) Percentage of bases that mapped to different regions of the sense strand of protein-coding genes in the nuclear genome. Because the cellular RNAs had not been chemically fragmented and the variety of TGIRT enzyme used in this study does not efficiently read through poly(A) at the 3’ end of mature mRNAs, protein-coding gene RNAs comprised only a low percentage of total reads (0.7-5.3%). Abbreviations: CDS, coding sequences; intergenic, regions upstream or downstream of transcription start and stop sites of protein-coding genes annotated in Ensembl; Intron, intronic regions; UTR, 5’- or 3’-untranslated regions.

**Figure S2.**
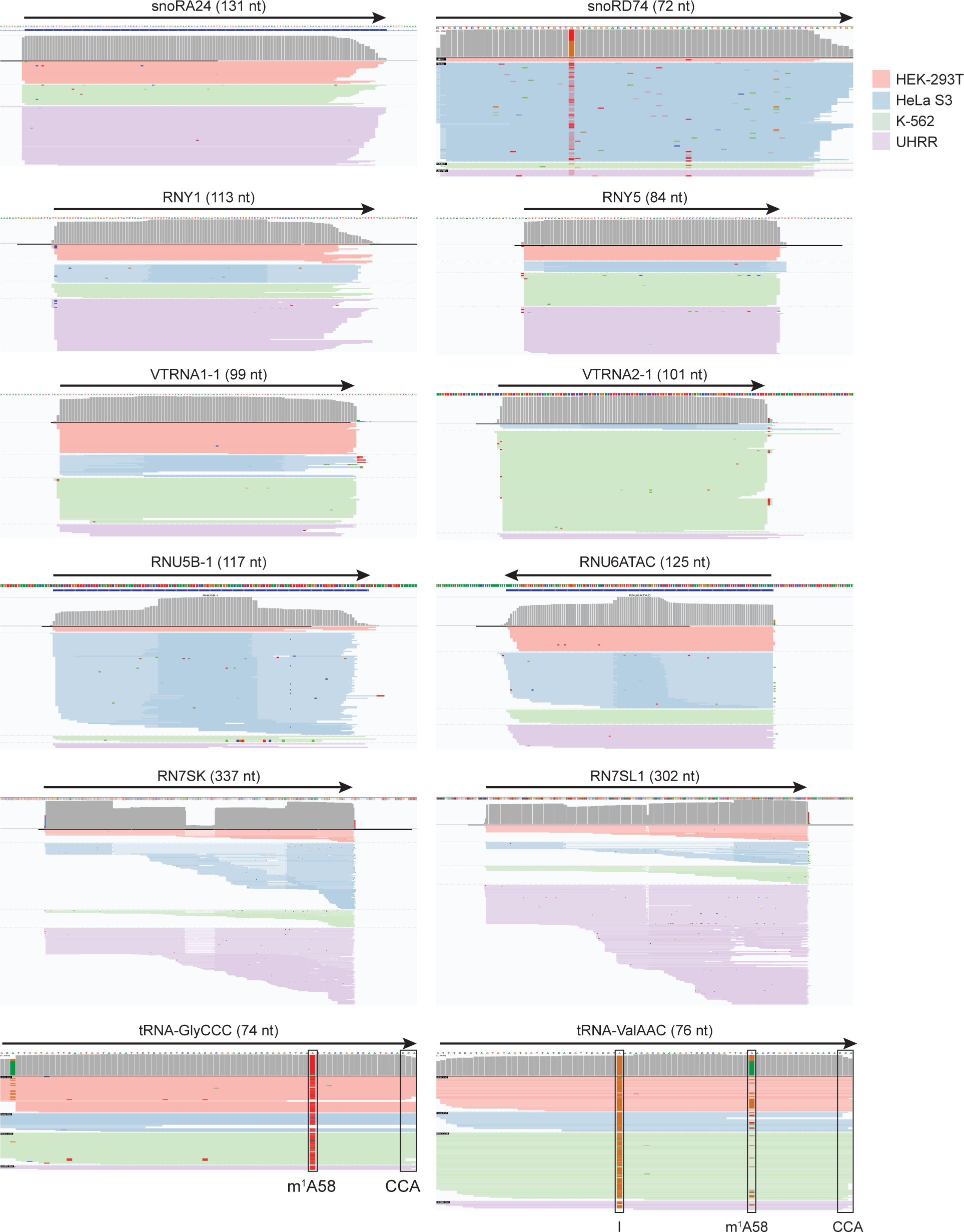
Full-length, end-to-end sequencing reads of sncRNAs in TGIRT-seq cellular datasets. Integrative Genomics Viewer (IGV) screenshots showing coverage tracks and read alignments for sncRNAs detected in TGIRT-seq datasets of unfragmented cellular RNAs. The name of the sncRNA is shown at the top with its length indicated in parentheses and the arrow indicating the 5’ to 3’ orientation of the RNA. Coverage tracks (gray) and read alignments for combined technical replicates of each cellular RNA sample type color coded as indicated in the Figure are shown below. Reads were down sampled to a maximum of 100 for display in IGV. Gray in the coverage tracts indicates bases in the read that matched the reference base, and other colors indicate bases in the read that did not match the reference base (red, thymidine; green, adenosine; blue, cytidine; and brown, guanosine). Misincorporation at known sites of posttranscriptionally modified bases in tRNAs are highlighted in the alignments: m^1^A58: 1-methyladenosine at position 58; I: inosine. CCA indicates the post-transcriptionally added 3’ CCA sequences of tRNAs mapped against a reference set of mature tRNA sequences.

**Figure S3.**
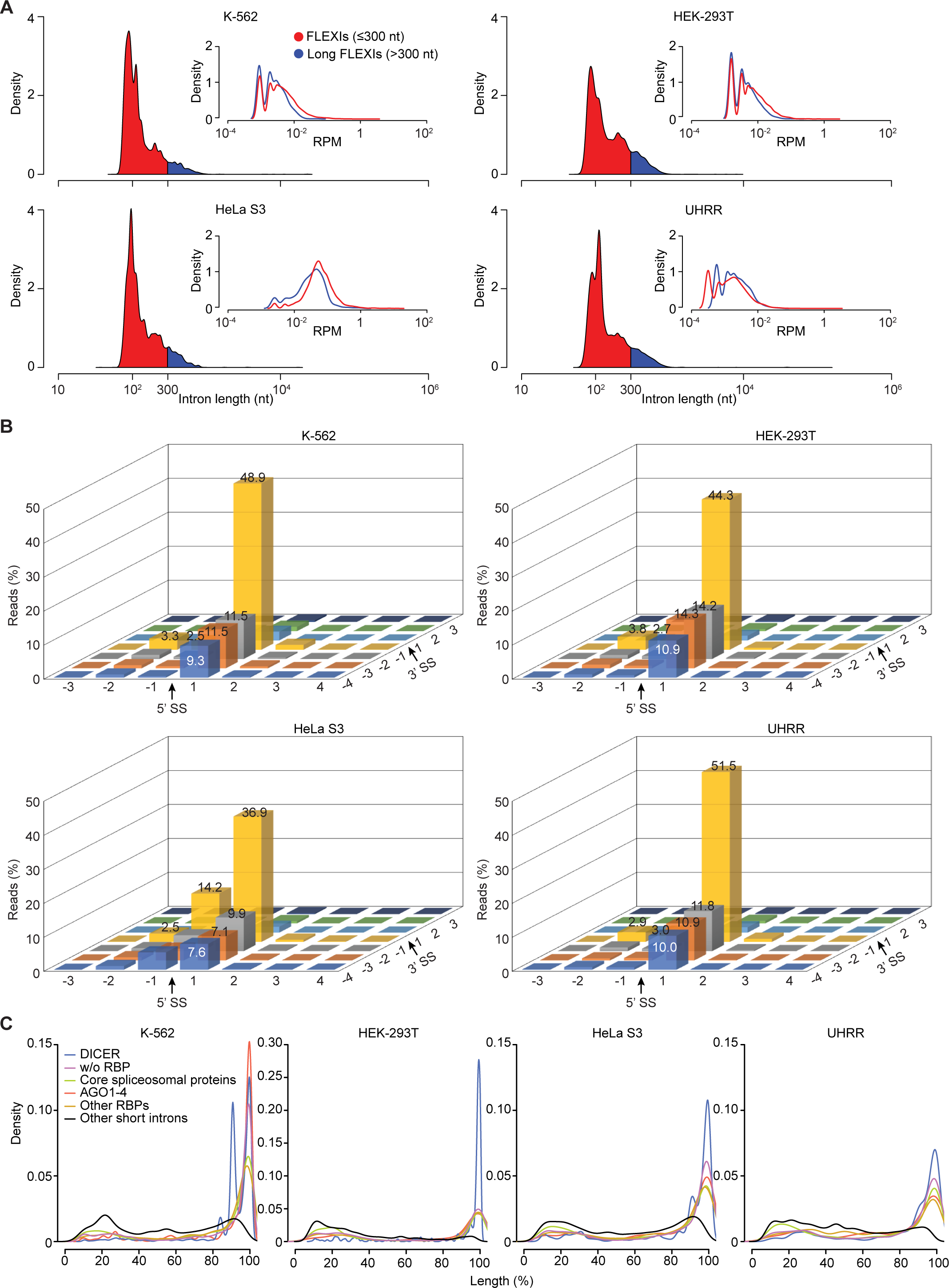
Characteristics of FLEXI RNAs. (A) Density plots showing the length distribution of FLEXI RNAs (≤300 nt, red) and small numbers of longer FLEXI RNAs (>300 nt, blue) in combined datasets for each of the 4 cellular RNA samples. The inset density plots compare the abundance (RPM) of FLEXIs ≤300 nt and >300 nt in the same datasets. (B) Three-dimensional bar chart showing the percentage of FLEXI RNA reads ending at different positions around intron-exon junctions in the combined datasets for each of the four cellular RNA samples. (C) Density plots showing the length distribution of all reads of any length mapping within different categories of FLEXIs compared to those mapping to other Ensemble GRCh38-annotated short introns (≤300 nt) in the same combined datasets. Separate plots color coded by category as shown in the Figure are overlaid for FLEXI RNAs containing annotated binding sites for AGO1-4, DICER, 6 core spliceosome proteins (AQR, BUD13, EFTUD2, PPIG, PRPF8 and SF3B4), other annotated RBPs, FLEXIs without (w/o) annotated RBP-binding sites, and other GRCh38-annotated short introns (≤300 nt). Percent length was calculated from the read spans of TGIRT-seq reads normalized to the length of the corresponding intron based on the Ensembl GRCh38 Release 93 human genome reference. FLEXIs encoding embedded snoRNAs/scaRNAs were excluded from the comparisons to avoid contributions from full or partially processed sncRNAs.

**Figure S4.**
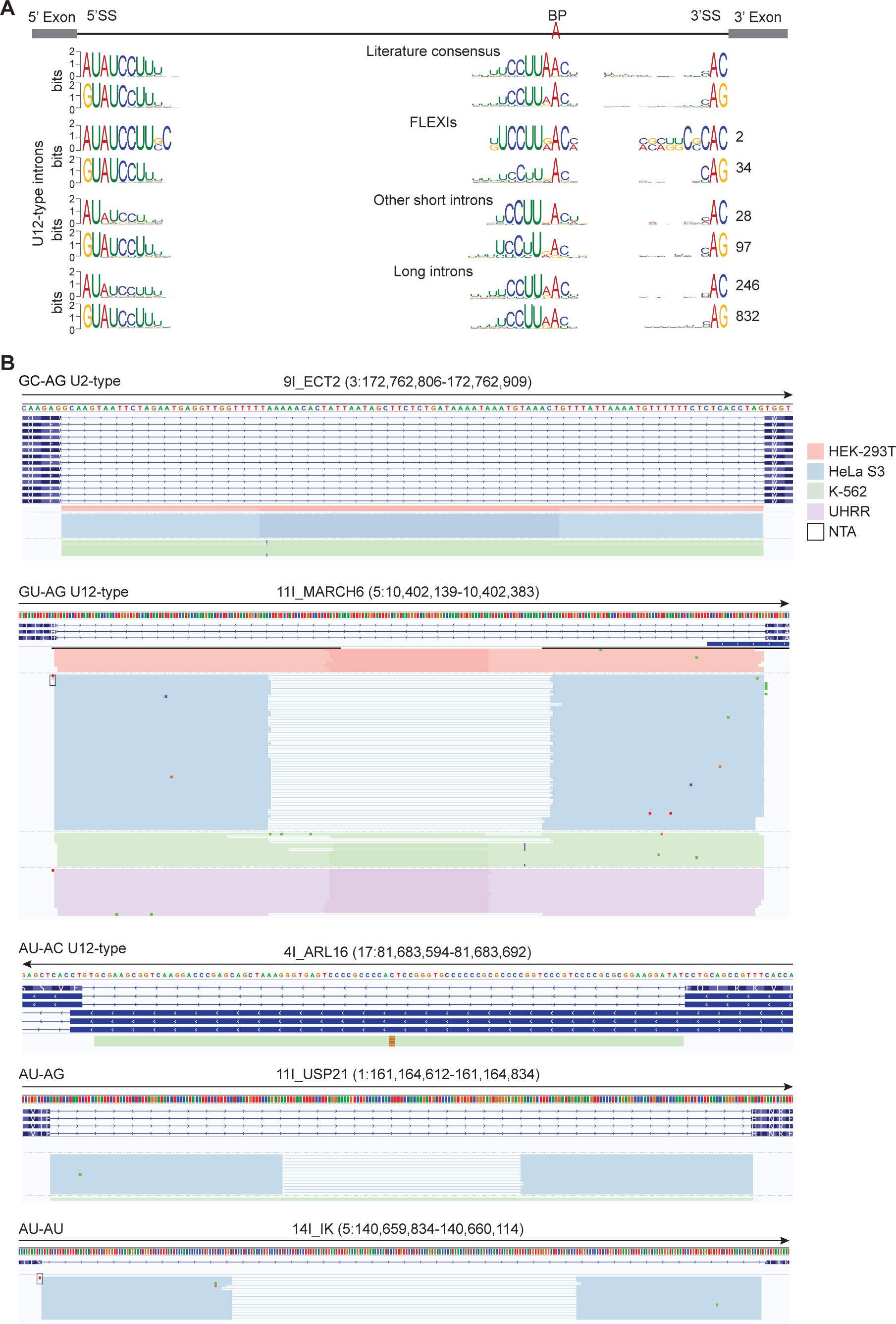
FLEXI RNAs corresponding to U12-type introns or introns with non-GU-AG 5’- and 3’-splice sites. (A) 5’- and 3’-splice sites (5’SS and 3’SS, respectively) and branch-point (BP) consensus sequences of GU-AG and AU-AC U12-type FLEXIs in a combined datasets for the 4 cellular RNA samples (HEK-293T, HeLa S3, K-562, and UHRR) compared to literature consensus sequences for human U12-type introns (top) (28, 30) and consensus sequences for other U12-type short (≤300 nt) and long (>300 nt) introns in the same combined datasets. The number of introns for each consensus sequence is indicated to the right of the sequence. (B) IGV screen-shots showing examples of coverage tracks and read alignments for FLEXI RNAs having U12-type or non-canonical 5’- and 3’-splice sites. Dinucleotides at the 5’- and 3’-ends of the intron are indicated at the upper left. The host gene name and genomic coordinates of the FLEXI are shown at the top with the arrow below indicating the 5’ to 3’ orientation of the encoded RNA followed by tracks showing the genomic sequence and gene annotations for different transcript isoforms (exons, thick bars; introns, thin lines). Read alignments for FLEXI RNAs are shown below the coverage tracks and are color coded by sample type as indicated in the key at the upper right of the panel. Boxes indicate non-templated addition (NTA) by TGIRT-III. Reads were down sampled to a maximum of 100 for display in IGV.

**Figure S5.**
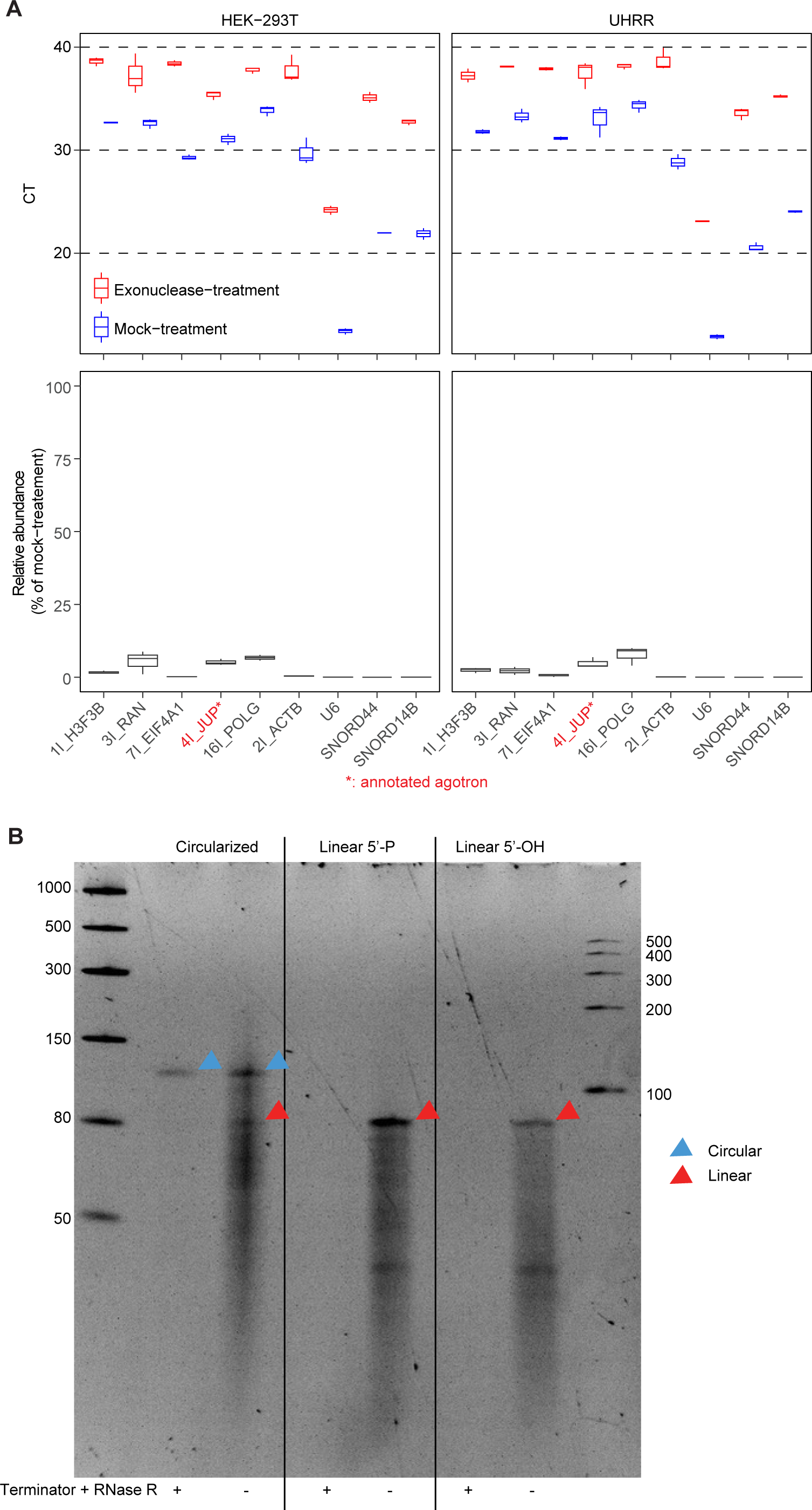
FLEXI RNAs are degraded by 5’- and 3’-exonucleases. DNase-treated, size-selected (≤200 nt) HEK-293T and UHRR RNAs were incubated with (+) or without (mock treatment; -) Terminator and RNase R exonucleases, and the products for 6 different FLEXI RNAs and 3 sncRNAs products were analyzed by RT-qPCR using SYBR Green with genespecific DNA oligonucleotide primers near the 3’ end of the RNA (Table S5; see Methods). (A) Cycle threshold (CT) values for qPCRs of exonuclease- and mock-treated FLEXIs and sncRNAs (top panel) and calculated percentage of RNAs remaining after exonuclease treatment relative to the parallel mock treatment (bottom panel). Three experimental replicates of exonuclease digestion and matched mock-treatments were performed for each RNA type. Each experimental replicate was quantified as the CT mean of 3 qPCR technical replicates and displayed as a box plot. The specificity of PCR products was confirmed by extracting gel bands corresponding to representative qPCR amplicons followed by Sanger sequencing directly or after TOPO-TA cloning (Supplementary file). (B) Denaturing PAGE of synthetic 3I_RAN FLEXI RNA that was circularized *in vitro* by T4 RNA Ligase 1, linear with a 5’ phosphate required for circularization, or linear with a 5’ OH after mock (-) or exonuclease treatment (+) with Terminator and RNase R exonucleases. Size markers in the left-most lane are the Low range ssRNA ladder (New England Biolabs), and those in the right-most lane are the RNA Century Ladder (Ambion). Circular and linear forms are indicated by arrowheads color coded as shown to the right of the gel.

**Figure S6.**
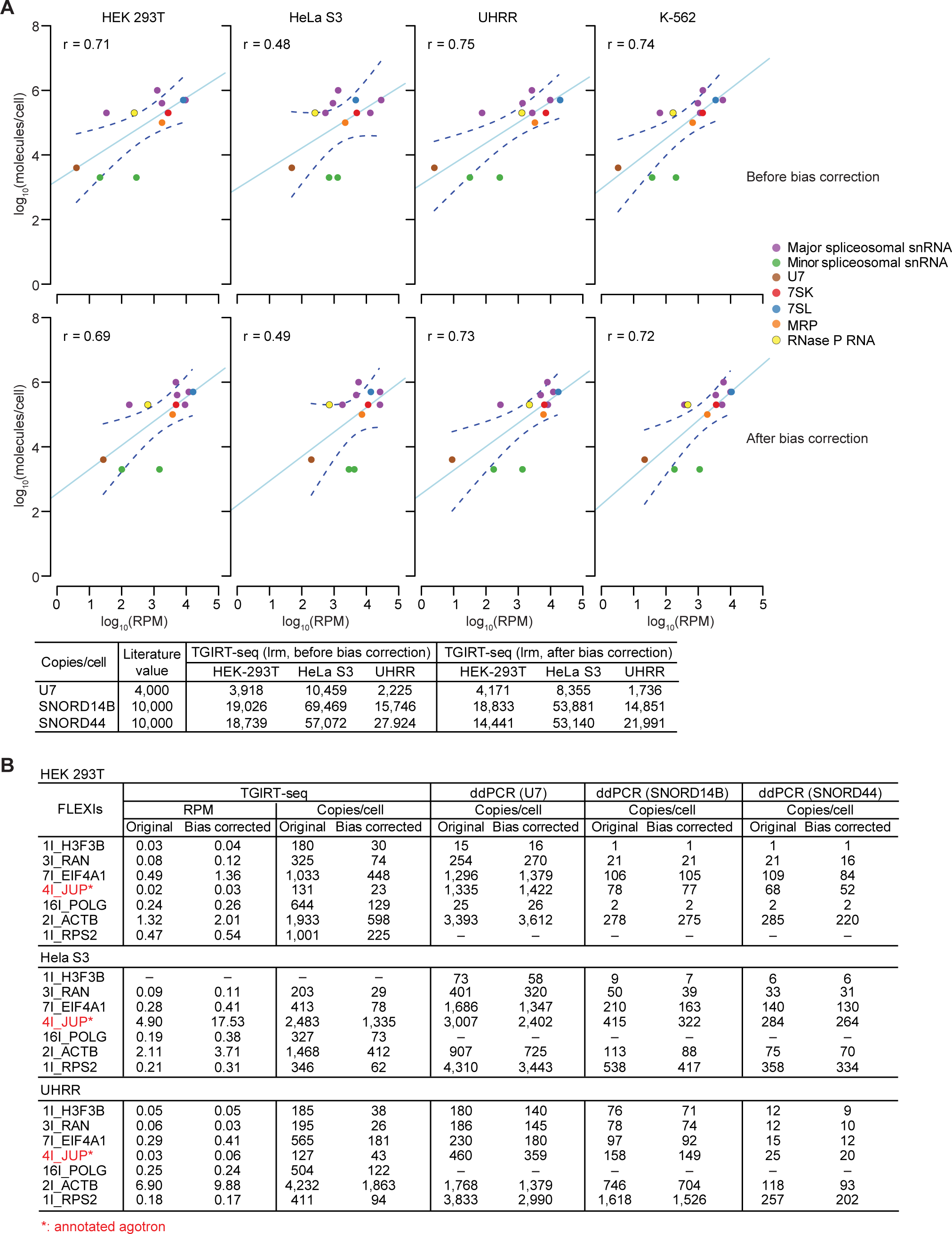
Estimates of FLEXI RNA abundance using a linear regression model based on sncRNA standards in TGIRT-seq datasets and droplet digital PCR. (A) Linear regression of molecules per cell values based on RPM values of sncRNAs of known cellular abundance in TGIRT-seq datasets. The scatter plots show the relationship between log_10_-transformed copy number per cell values for sncRNAs reported in the literature (71) and log_10_ transformed RPM of sncRNAs in combined datasets for each of the 4 cellular RNA samples before (top panels) or after (bottom) bias correction of TGIRT-seq reads using a random forest regression model (21). The linear regression was plotted as a light blue line with 95% confidence intervals shown as dashed blue lines. sncRNA used in the linear regression were spliceosomal snRNAs U1, U2, U4, U5, and U6; minor spliceosomal snRNAs U4ATAC and U6ATAC; U7, 7SL and 7SK RNA; MRP and RNase P RNA (Table S2) (71). Pearson (*r*) correlation coefficients are shown in the upper left of each panel. A test of the linear regression model (*lrm*) for 3 sncRNAs (U7, SNORD14B, and SNORD44), used as digital droplet PCR (ddPCR) standards described in panel B, gave copy number per cell estimates within the same order of magnitude as literature values (71), but with copy number per cell values varying between different cell types (Table at the bottom of panel A). (B) Abundance estimates for FLEXI RNAs based on the TGIRT-seq linear regression model and droplet digital PCR (ddPCR). The left-hand columns show RPM and copy number per cell values of FLEXIs, estimated using the linear regression model for cellular RNA samples. The right-hand columns show copy number per cell values based on ddPCR abundance relative to 3 different sncRNA standards run in parallel (U7, SNORD14B, and SNORD44). For ddPCR, reverse transcription was performed on DNase-treated, size-selected (≤200 nt) RNA preparations using Maxima H reverse transcriptase with a gene-specific primer at the 3’ end of the putative FLEXI or sncRNA. ddPCR was then performed using primer sets within the FLEXI sequence or spanning the 5’ exon-FLEXI junction, with the latter subtracted from the FLEXI signal when applicable. Copy number per cell values for FLEXIs were calculated based on copy number per cell values for U7, SNORD14B, and SNORD44 in each cellular RNA sample, calculated using the linear regression model without or with TGIRT-seq bias correction (see Table at the bottom of panel A). Primer amplification specificity was verified by gel extraction of a representative set of matched qPCR amplicons, which were then Sanger sequenced directly or after TOPO-TA cloning. The ddPCR values relative to U7 RNA, which has the closest cellular abundance to FLEXIs by TGIRT-seq reads, showed general agreement with the copy number per cell values estimated from the linear regression model without TGIRT-seq bias correction, while the other sncRNA controls, predicted to be in much higher abundance than FLEXIs, gave FLEXI quantifications closer to the bias-corrected linear regression model.

**Figure S7.**
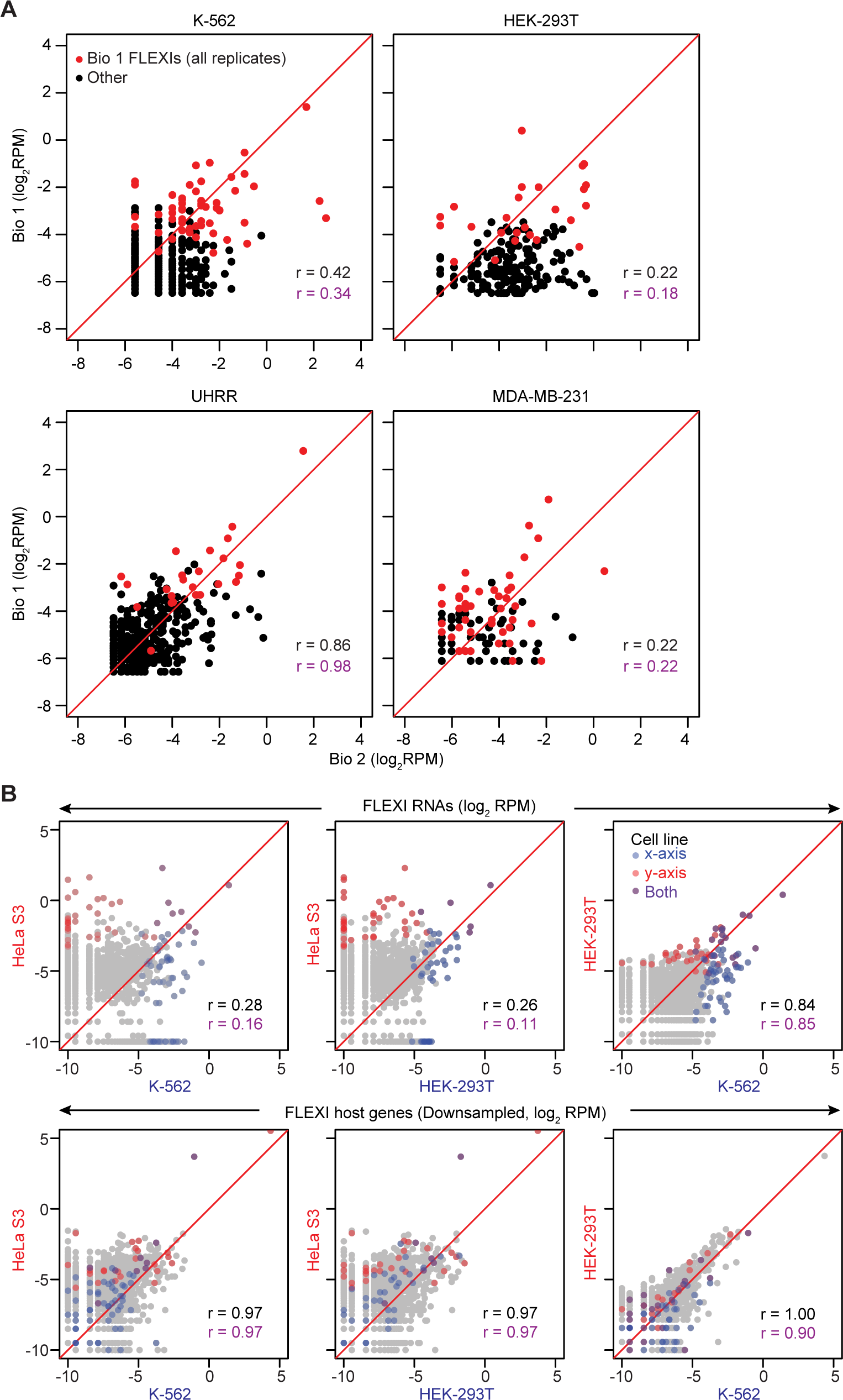
Cell specificity and reproducibility of detected FLEXI RNAs. (A) Scatter plots comparing log_2_-transformed RPM of relatively abundant (≥0.01 RPM) FLEXI RNAs in biological replicates (Bio 1 and Bio 2) for 4 different cellular RNA samples. Bio 1 is a merged dataset of all technical replicates for the indicated cellular RNA samples obtained for this study, and Bio 2 is a merged dataset for all technical replicates of different RNA preparations from the same cell types obtained for other studies (including a different lot number for UHRR; see Table S1). FLEXIs that were detected in all Bio 1 technical replicates are shown as red dots. (B) Scatter plots comparing log_2_-transformed RPM of FLEXI RNA reads in different cellular RNA samples (top row) to those mapping to FLEXI host genes in the same datasets down sampled to match the sequencing depth of FLEXI reads (bottom row). Relatively abundant FLEXIs (≥0.01 RPM) detected in all replicates of each cell type and their host genes are color coded as shown in the top right scatter plot. Pearson correlation coefficients (*r*) calculated for all detected FLEXIs and their host genes are shown in black, and those for relatively abundant color-coded FLEXIs and their host genes (red and blue) are shown in purple.

**Figure S8.**
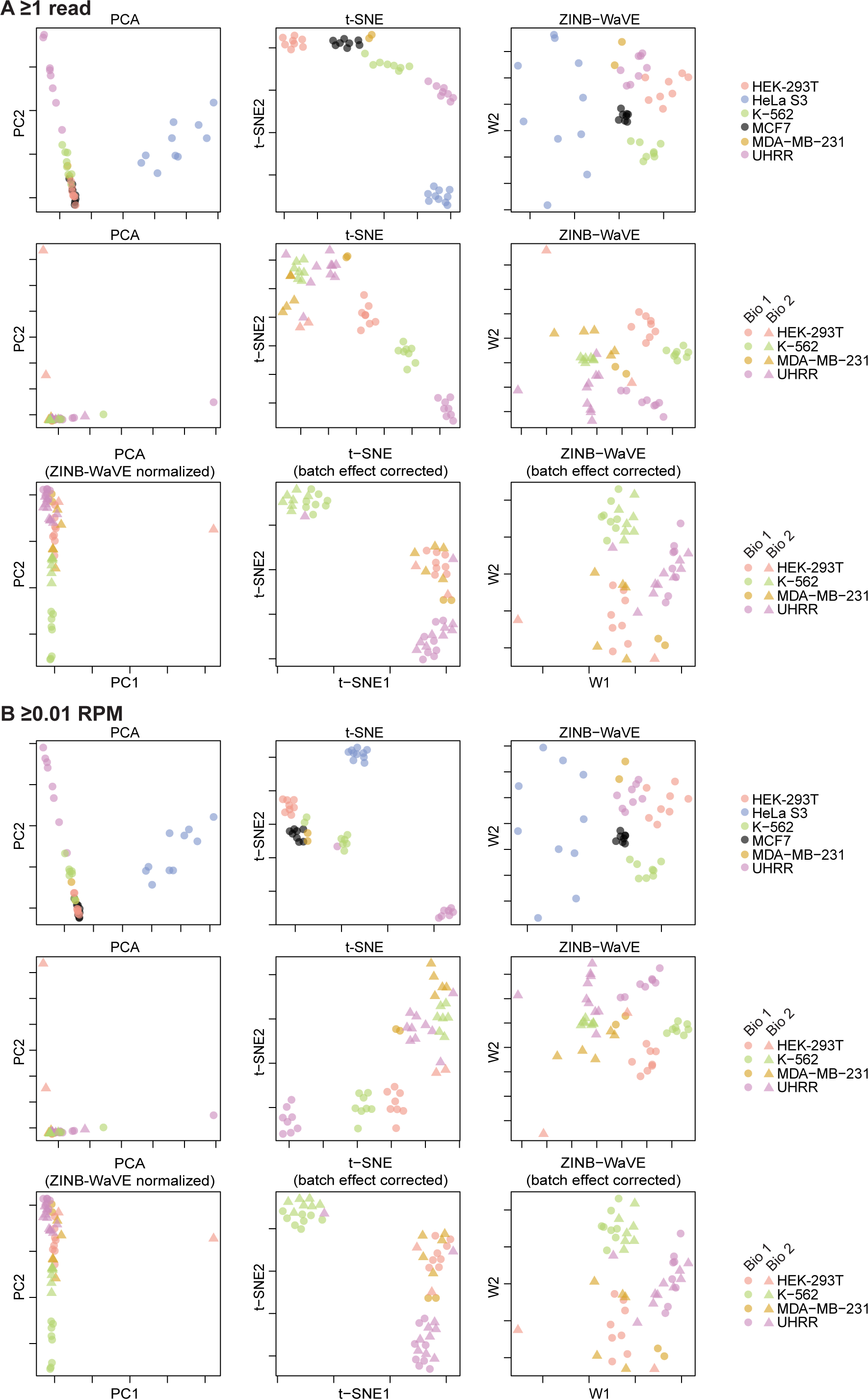
Clustering analysis of detected FLEXI RNAs. (A) and (B) PCA, PCA-initialized t-SNE (33) and ZINB-WaVE (34) analysis for FLEXI RNAs detected at ≥1 read (panel A) or ≥0.01 RPM (panel B) in biological and technical replicates of rRNA-depleted unfragmented cellular RNAs (Table S1). Both t-SNE and ZINB-WaVE are widely used for the analysis of single cell RNA-seq datasets with zero-inflated counts. The top row of each panel shows clustering of datasets for technical replicates for different cell types used in this study. The middle and bottom rows show clustering of datasets for rRNA-depleted unfragmented HEK-293T, K-562, MDA-MB-231, and UHRR RNAs from this study (biological replicate 1 (Bio 1, circles) and other studies (biological replicate 2 (Bio 2), triangles) before (middle) and after (bottom) batch effect correction by ZINB-WaVE. All of the compared datasets are listed in Table S1.

**Figure S9.**
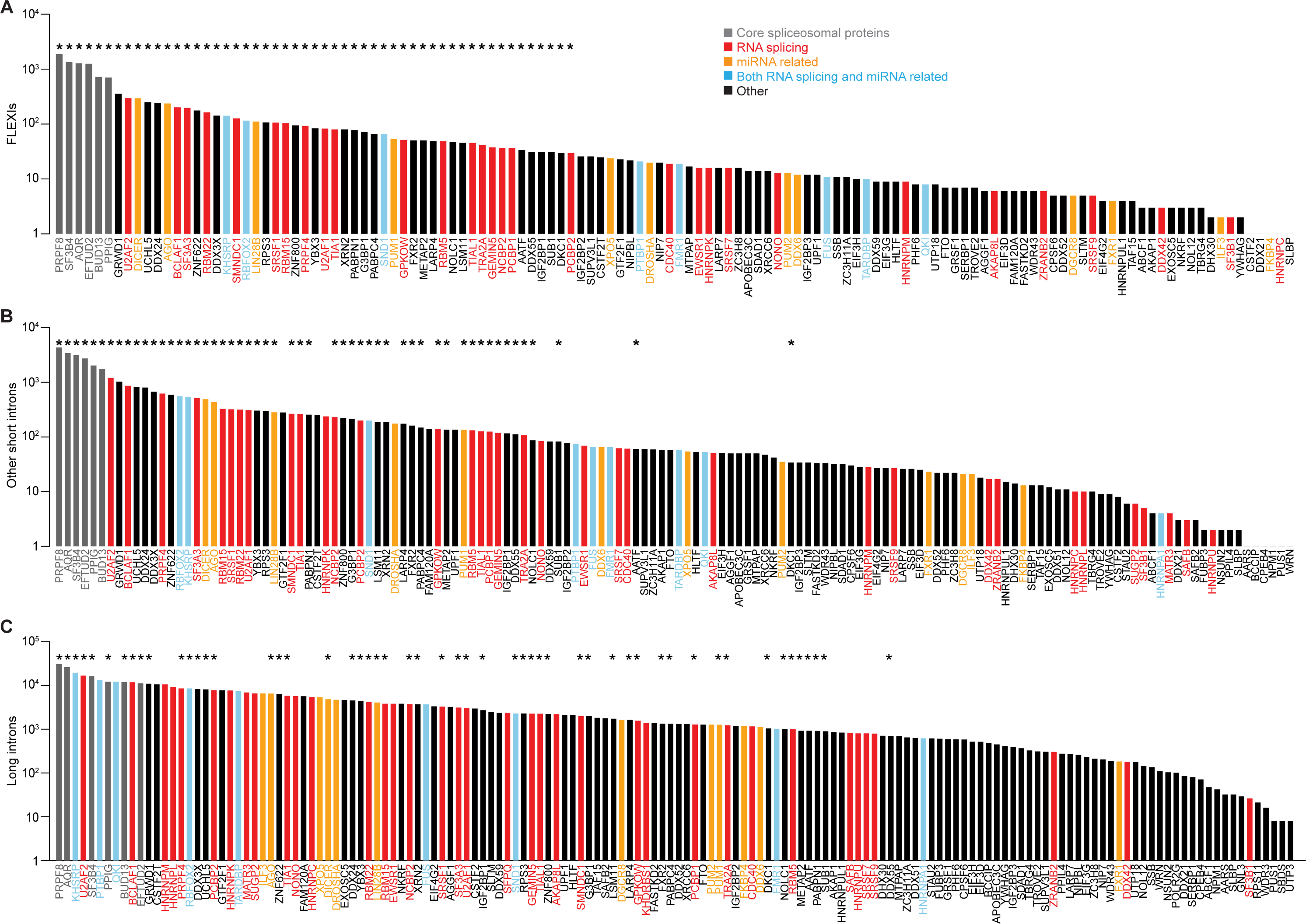
Quantitation of RBP-binding sites in different classes of intron RNAs. Bar graphs showing the number of (A) FLEXI RNAs, (B) Other Ensembl GRCh38-annotated short introns (≤300 nt), and (C) Ensembl GRCh38-annotated long introns (>300 nt) that have a CLIP-seq-identified binding site for the indicated RBP in a merged dataset for the K-562, HEK-293T, HeLa S3, and UHRR cellular RNA samples. Bars graphs are color coded by RBP function as shown at the top. Asterisks above the bars in panels B and C indicate the 53 proteins identified as binding 30 or more different FLEXIs in Fig. 4A.

**Figure S10.**
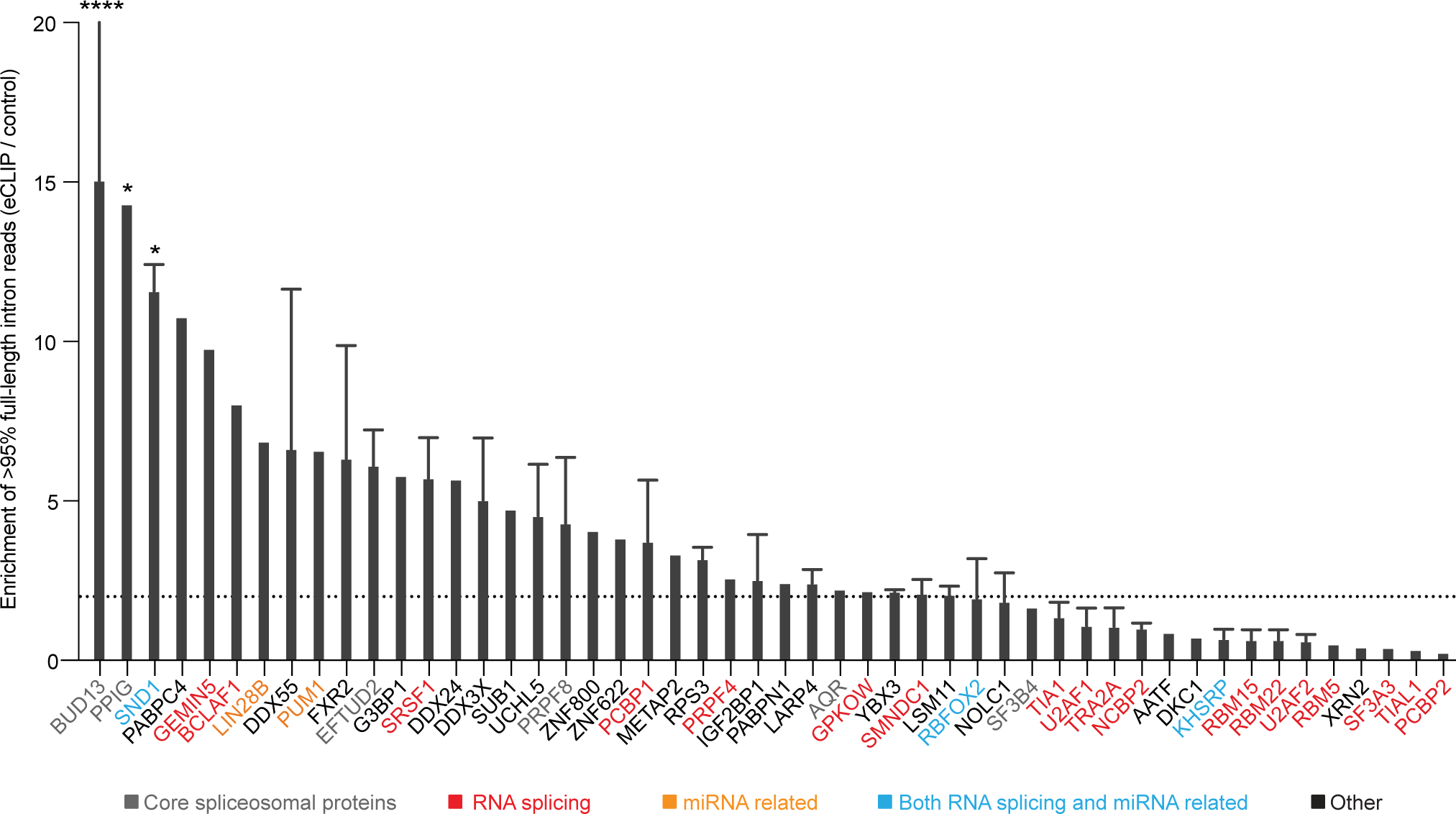
Sequencing reads corresponding to FLEXIs in ENCODE eCLIP datasets. eCLIP datasets for HepG2 and K-562 cells were obtained from ENCODE for all RBPs that bind ≥30 FLEXIs, and reads that were ≥95% of the length of FLEXIs were identified by intersecting the eCLIP reads with FLEXI coordinates. Data are shown as fold enrichment of ≥95% fulllength intron reads for each RBP, calculated as a ratio of such reads in the eCLIP dataset versus the control antibody dataset (**** p<0.0001, * p<0.05, Fisher’s least significant difference test). The dashed line indicates 2-fold enrichment. Names of proteins are color coded by protein function as indicated in the Figure. For those RBPs in which more than one dataset was available, mean values were calculated for the fold enrichment in eCLIP replicates over controls with error bars indicating the standard deviation (Supplementary file).

**Figure S11.**
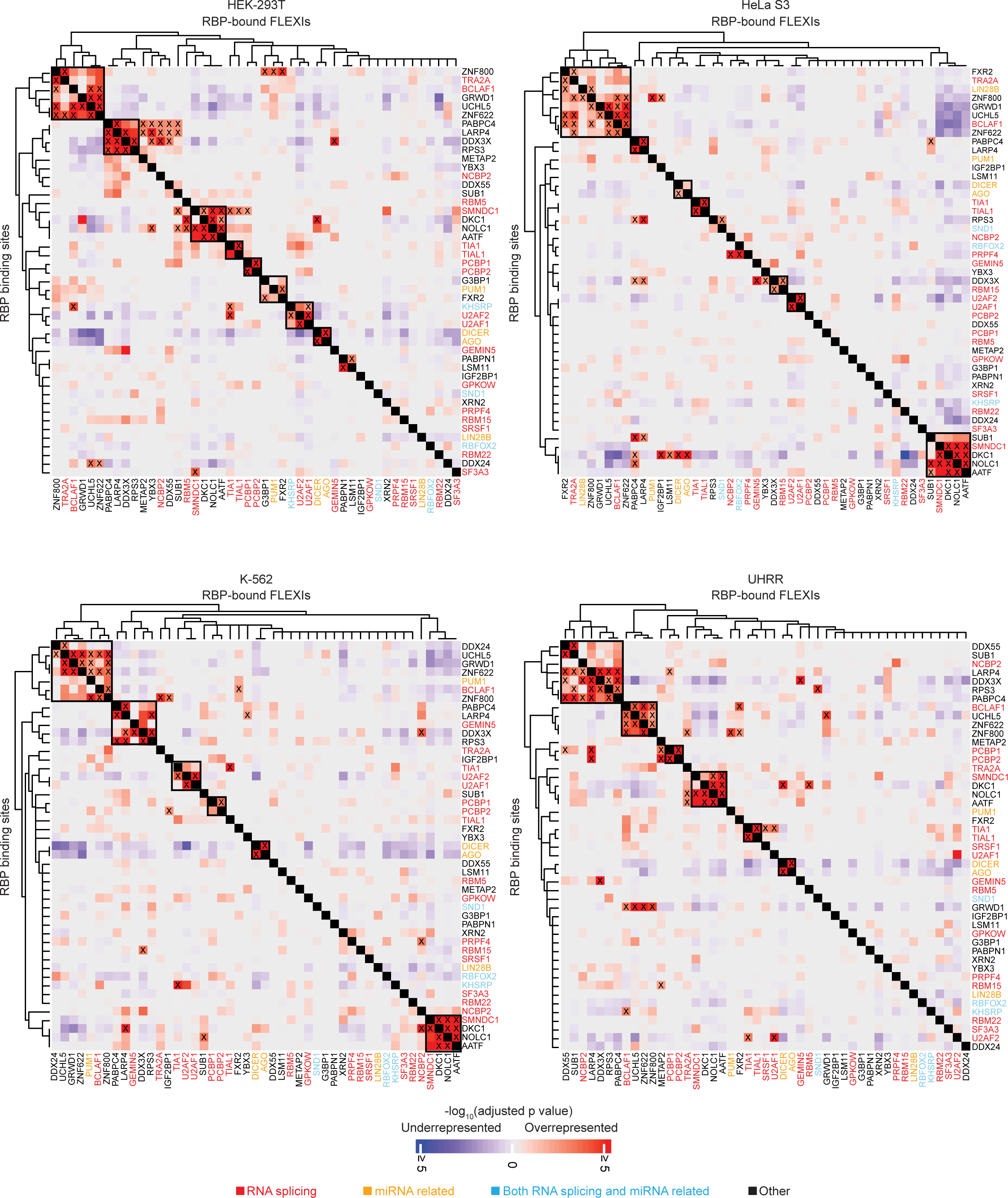
Hierarchical clustering of FLEXIs in different cell lines based on over- and under-represented RBP-binding sites. FLEXIs containing at least one annotated binding site for any RBP in any of the 4 cell lines (1,261 to 2,404 FLEXIs) were used in this analysis. For hierarchical clustering, subsets of FLEXIs (6 to 1,086) that have binding sites for each of the 47 noncore spliceosomal RBPs that bind ≥30 different FLEXI in merged TGIRT-seq datasets for the 4 cell lines were extracted for each of the cellular RNA sample types. Contingency tables were then generated by comparing the frequency of binding sites for all 126 RBPs that have an identified binding site in a FLEXI RNA in each of these subsets to those in all FLEXI RNAs. Multiple binding sites for the same RBP in the same FLEXI were counted as one binding site. p-values for over- and under-representation of the RBP-binding sites calculated by Fisher’s exact test were adjusted by the Benjamini-Hochberg procedure. RBP-binding sites were then key-coded as not significantly different or as significantly over- or under-represented (≥2% abundance, adjusted p≤0.05 calculated by Fisher’s exact test and adjusted by the Benjamini-Hochberg procedure) in the tested subset of FLEXIs compared to all FLEXIs, and the key-coded information was used to construct matrices of the Gower’s distance between the 47 subsets in each of the cellular RNA samples that were used as input for hierarchical clustering by the complete linkage clustering method (72). The results were displayed as a two-dimensional heat map to identify subsets of FLEXIs showing similar patterns of significantly over- and under-represented RBP-binding sites in each of the cellular RNA samples, with the color scale at the bottom based on -log_10_-transformed adjusted p-values. Significantly co-enriched RBPs are indicated by an X in the heatmap box, and clusters of co-enriched RBPs are delineated in larger, differently shaded boxes. RBP names are color coded by protein function as indicated in the Figure.

**Figure S12.**
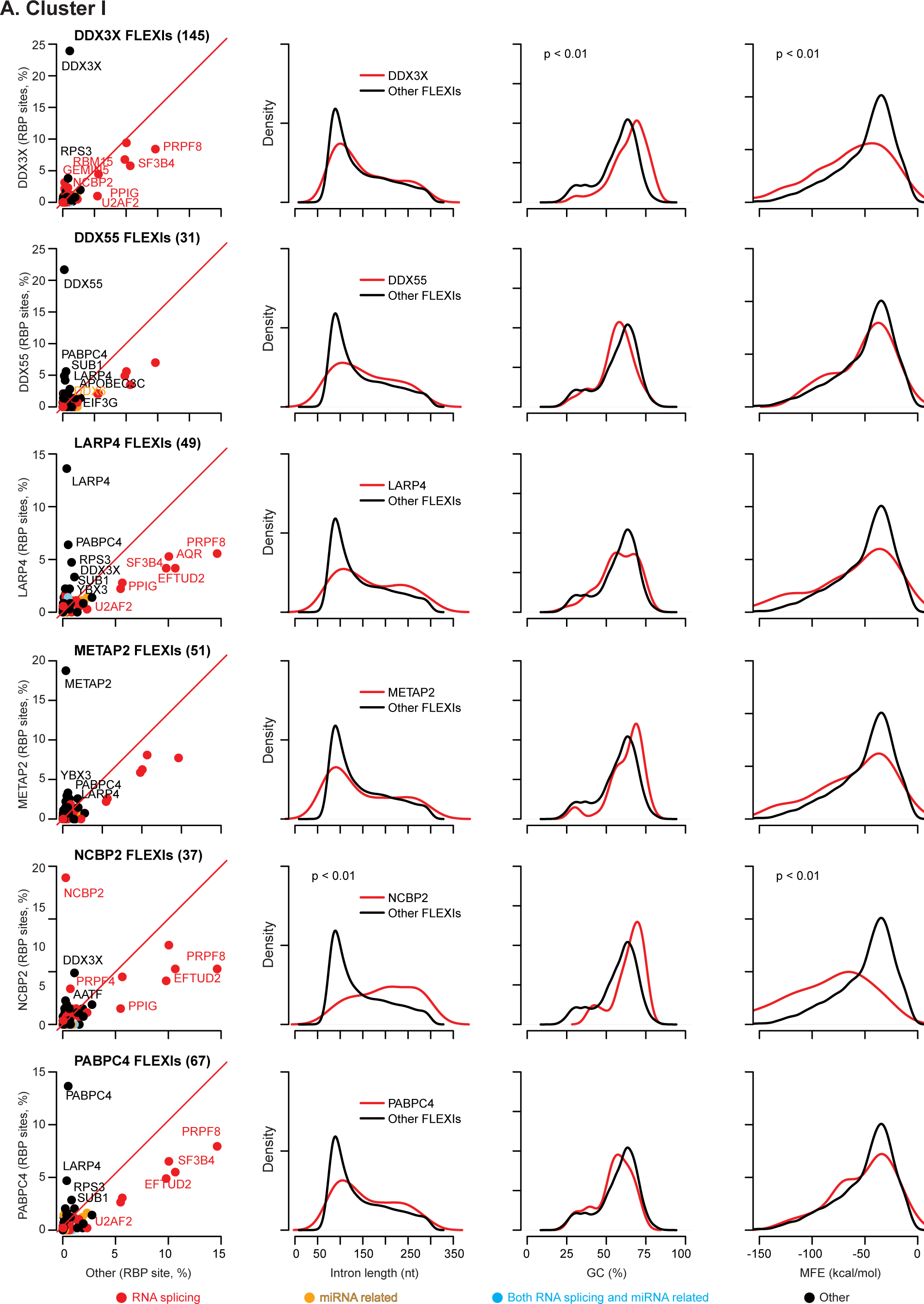
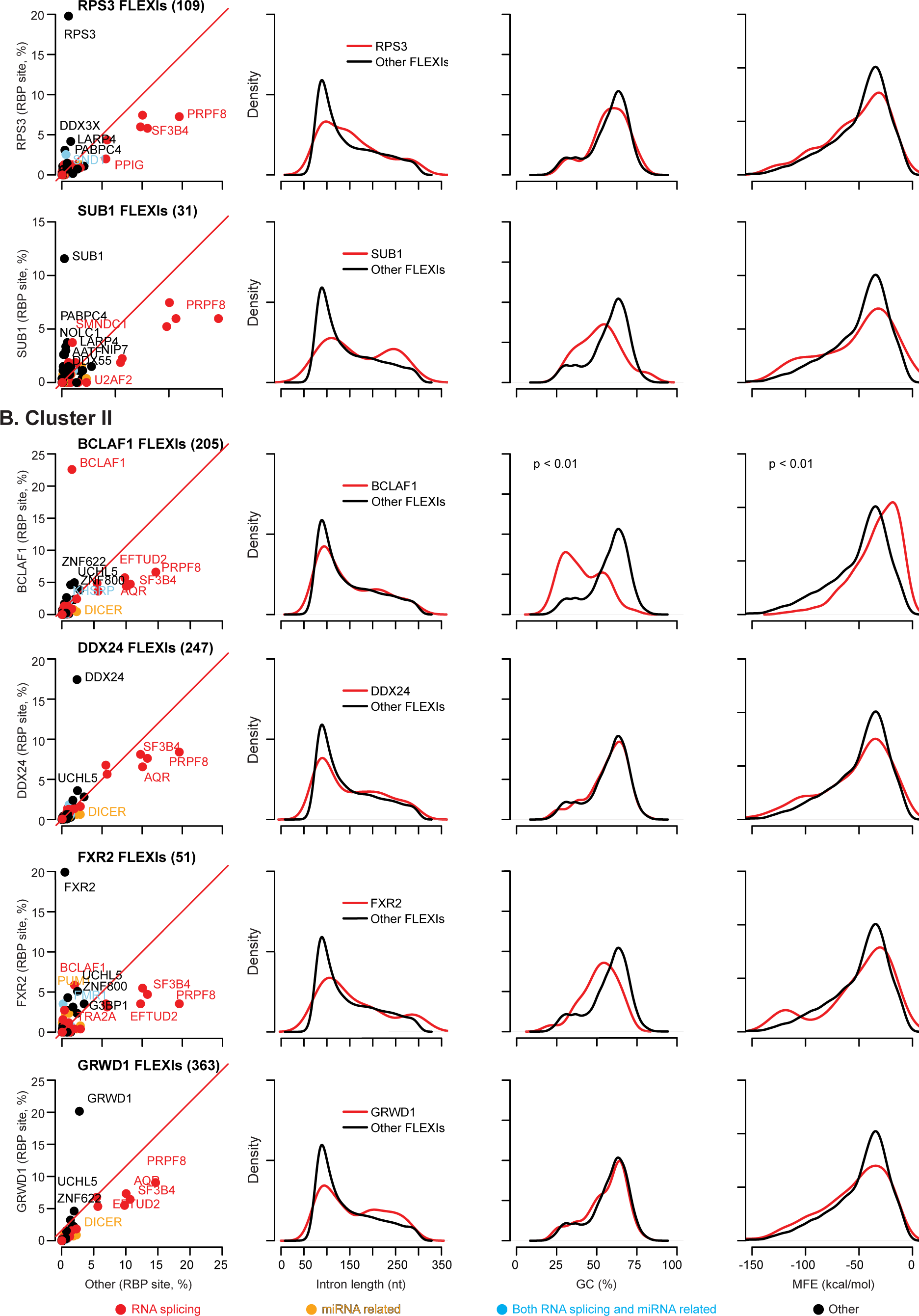
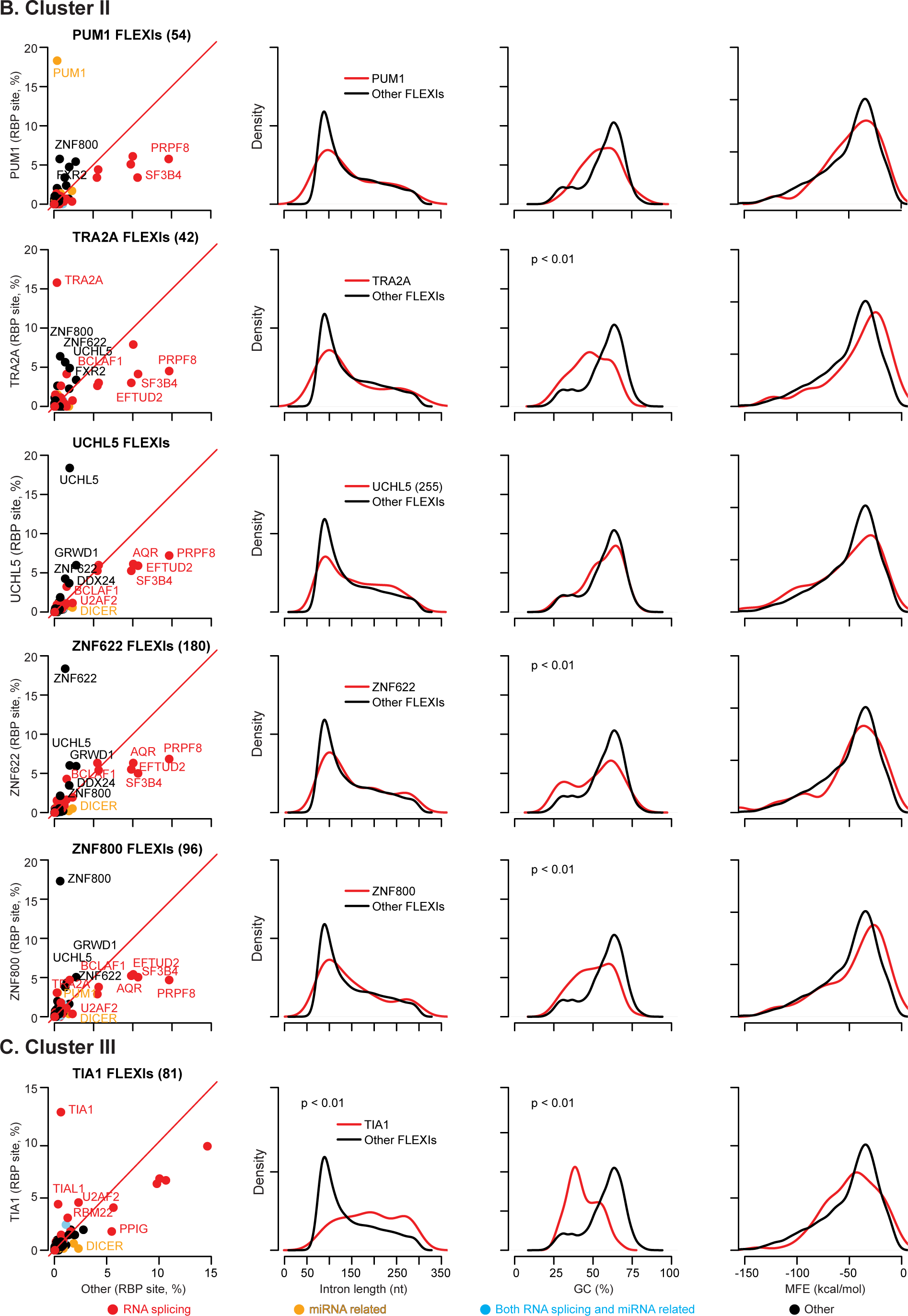
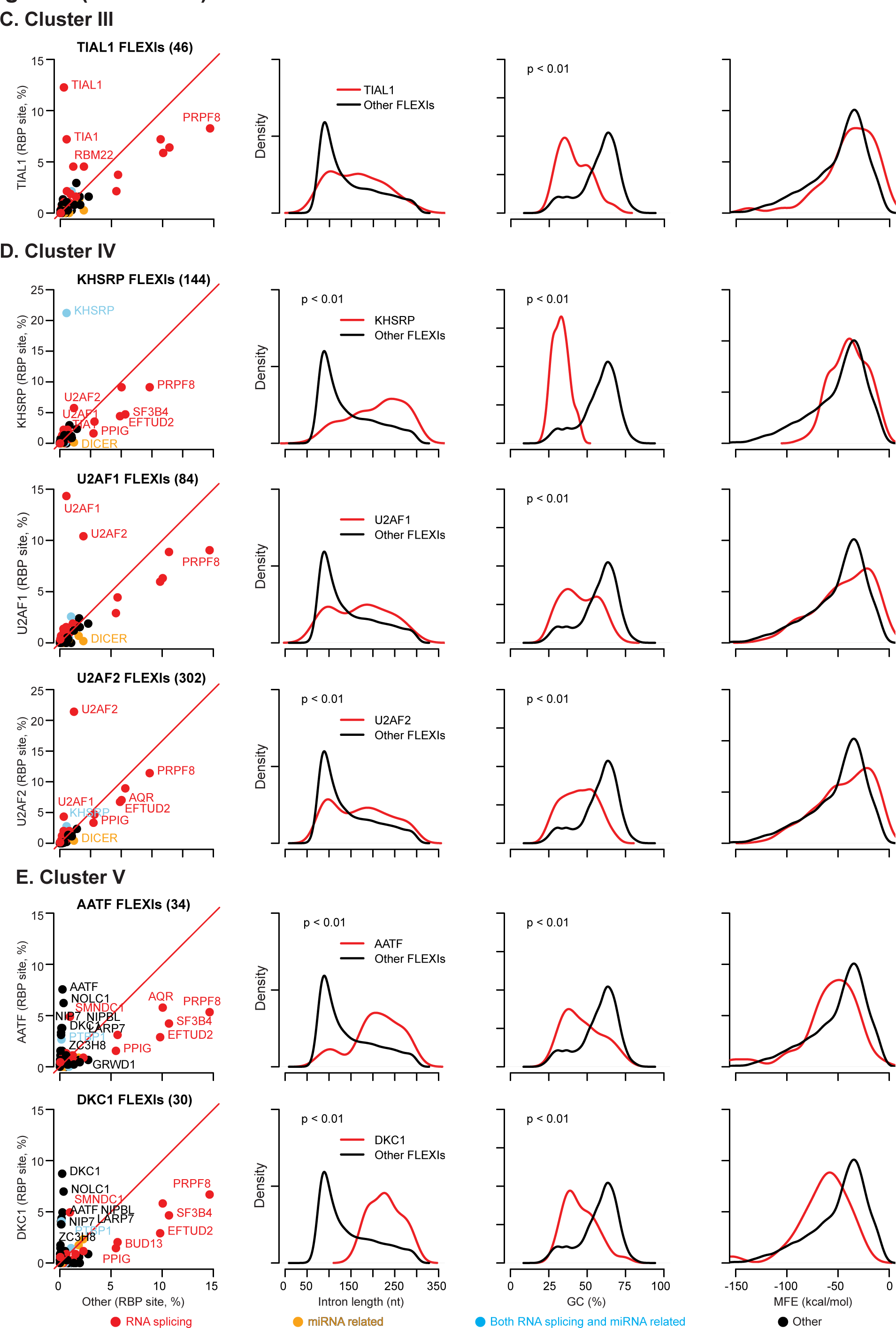
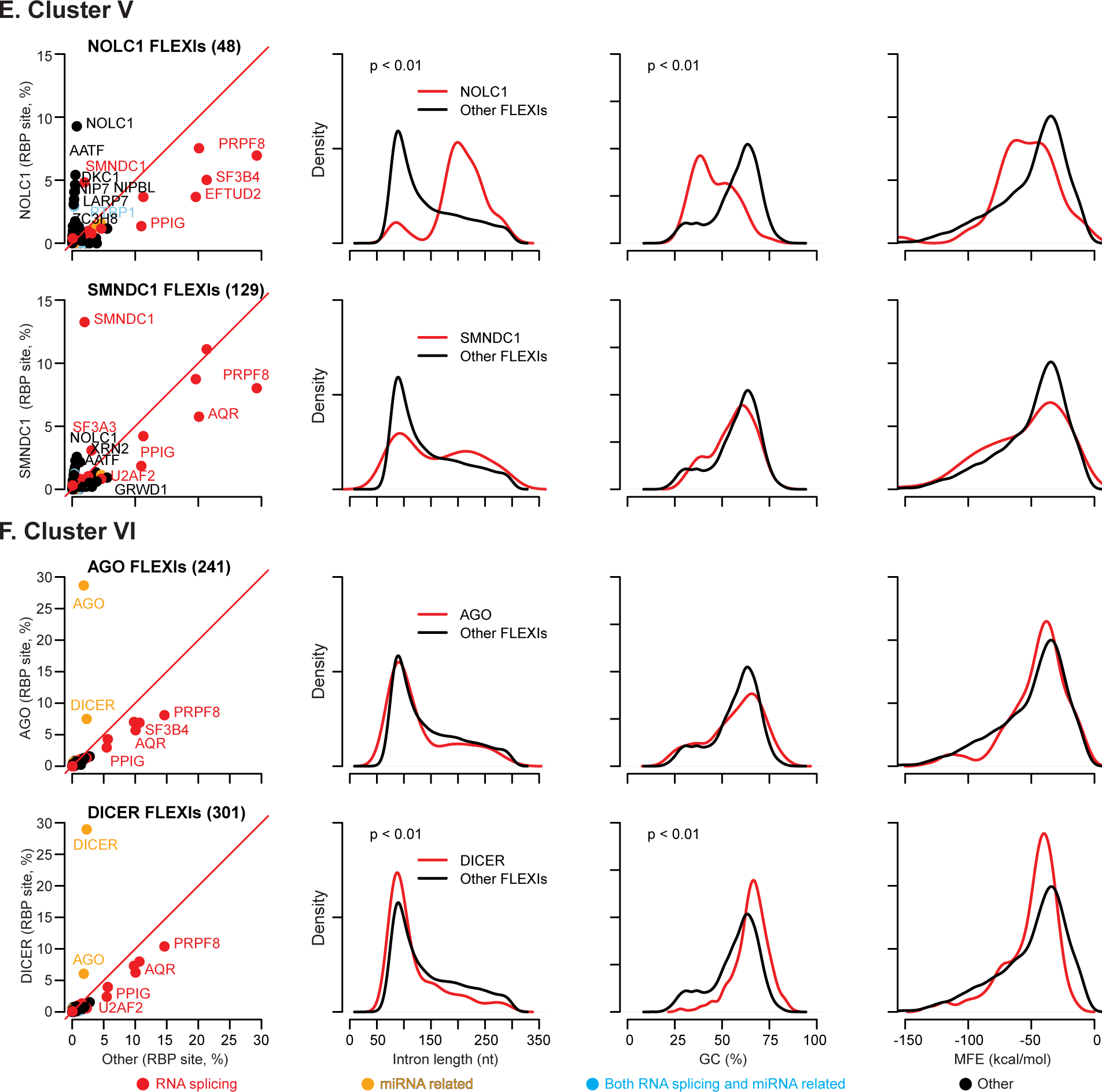
Enriched RBP-binding sites and characteristics of subsets of FLEXIs in Clusters I-VI. Scatter plots and density plots are shown for FLEXIs that bind each protein in (A) Cluster I, (B) Cluster II, (C) Cluster III, (D) Cluster IV, (E) Cluster V, and (F) Cluster VI. In the scatter plots (left), RBP-binding sites for other RBPs whose relative abundance was significantly over- or under-represented in the subset of FLEXIs compared to all other FLEXIs (≥2% abundance, p≤0.05 calculated by Fisher’s exact test and adjusted by the Benjamini-Hochberg procedure) are labeled by the name of the protein, color coded by protein function as indicated in the Figure. The density distribution plots (right) compare the length, GC content, and MFE for the most stable secondary structure predicted by RNAfold for subsets of FLEXIs bound by each RBP associated with Clusters I to VI (red) compared to all other FLEXIs (black). The number of FLEXIs comprising each subset is indicated in parentheses next to the name of the RBP. pvalues are shown at the top left of those density plots in which the distribution for the subset of FLEXIs differed significantly from other FLEXIs (p<0.01 and FDR≤0.05 as determined by 1,000 Monte-Carlo simulation).

**Figure S13.**
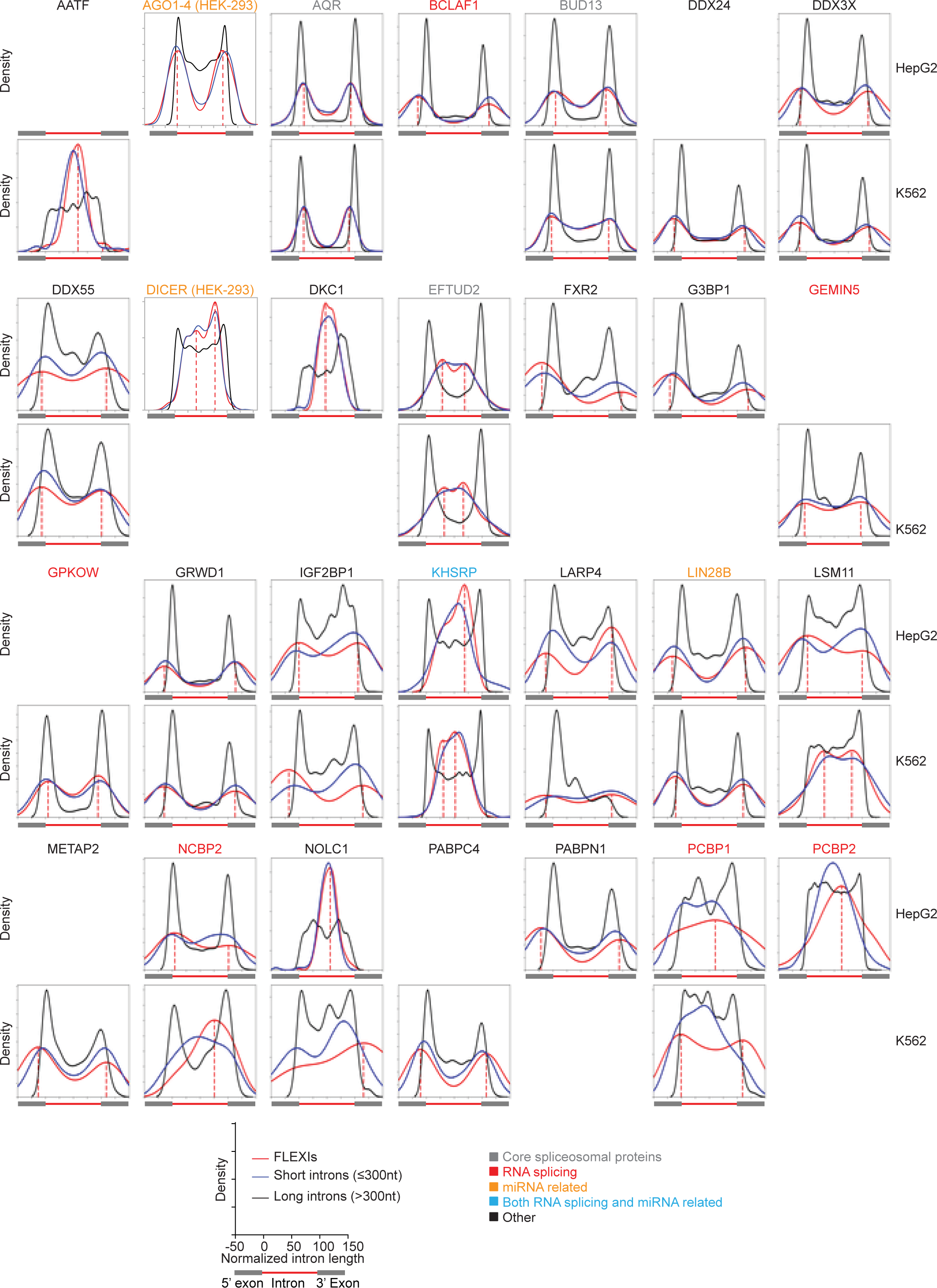
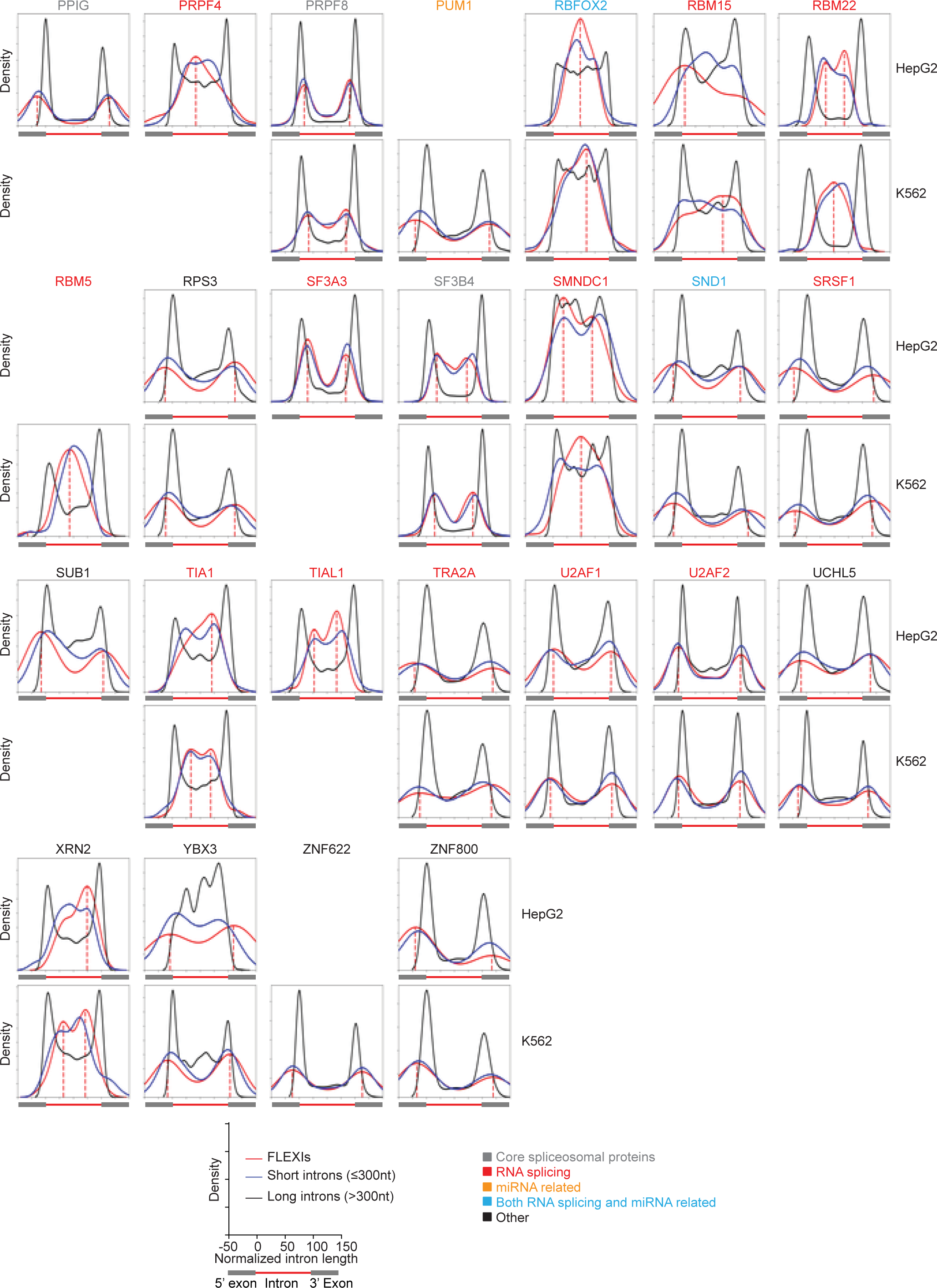
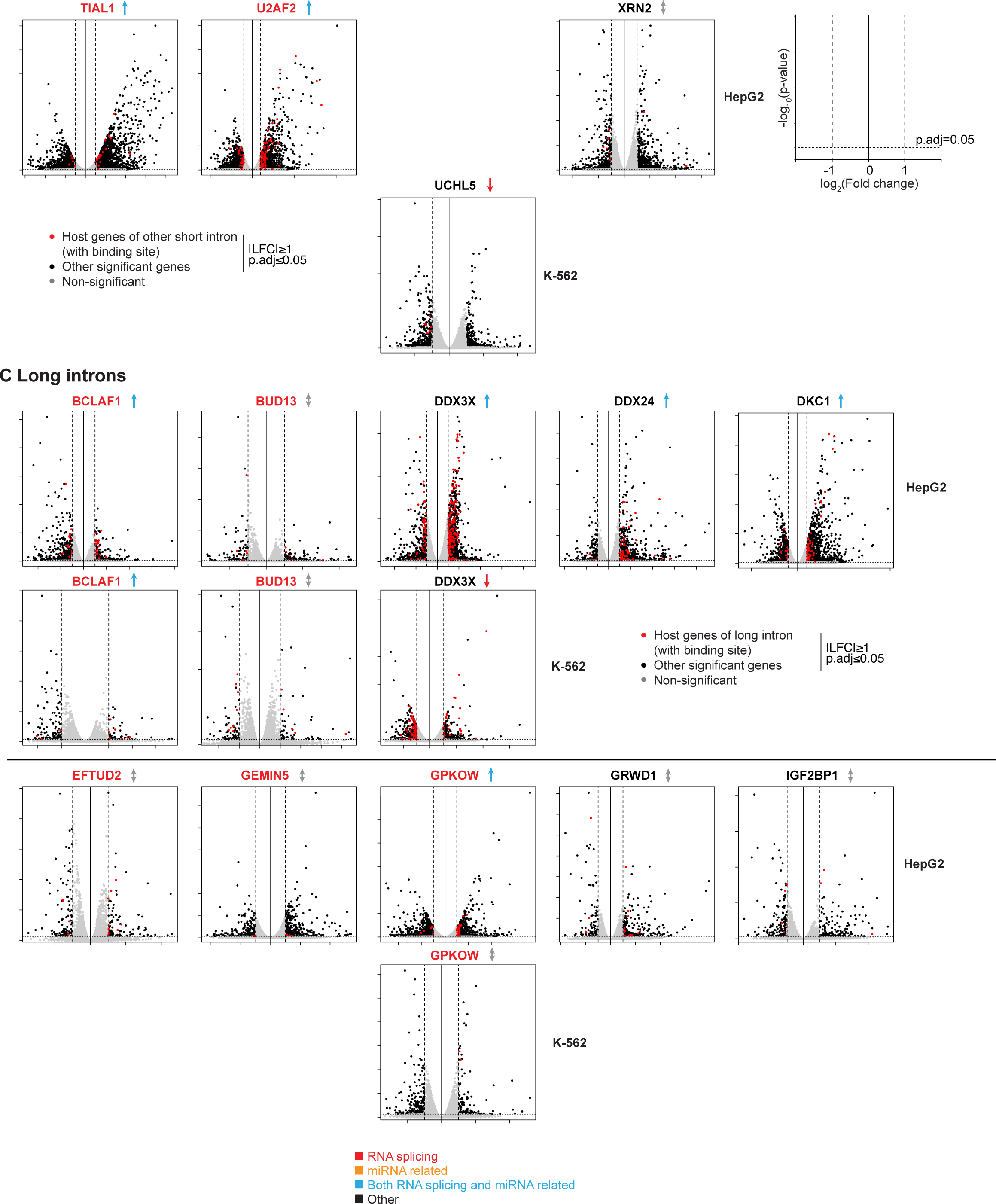
Relative positions of RBP-binding sites in FLEXIs and other classes of introns. Density plots showing the location of eCLIP-identified RBP-binding sites in K-562 or HepG2 cells within FLEXIs for the 53 RBPs that bind ≥30 FLEXIs in merged K-562, HEK-293T, HeLa S3, and UHRR datasets compared to other short and long introns. RBP-binding sites overlapping introns (≥1 nt) were identified by intersecting the annotated RBP-binding sites from the eCLIP datasets with the intron coordinates using BEDTools and plotted as the mid-point of the annotated RBP-binding sites normalized as a percentage of intron length, and with 0% and 100% corresponding to the 5’ and 3’ end of the intron, respectively. For each RBP, binding-site distributions are shown for FLEXIs (red), other short introns (≤300 nt; blue), and long introns (>300 nt; black). Vertical red dashed lines indicate the position of peaks in the density plots for FLEXI RNAs. RBP names are color coded by protein function as indicated at the bottom of the Figure. Blank spaces were left when datasets were not available for an RBP in one of the two cell lines used to obtain the eCLIP datasets.

**Figure S14.**
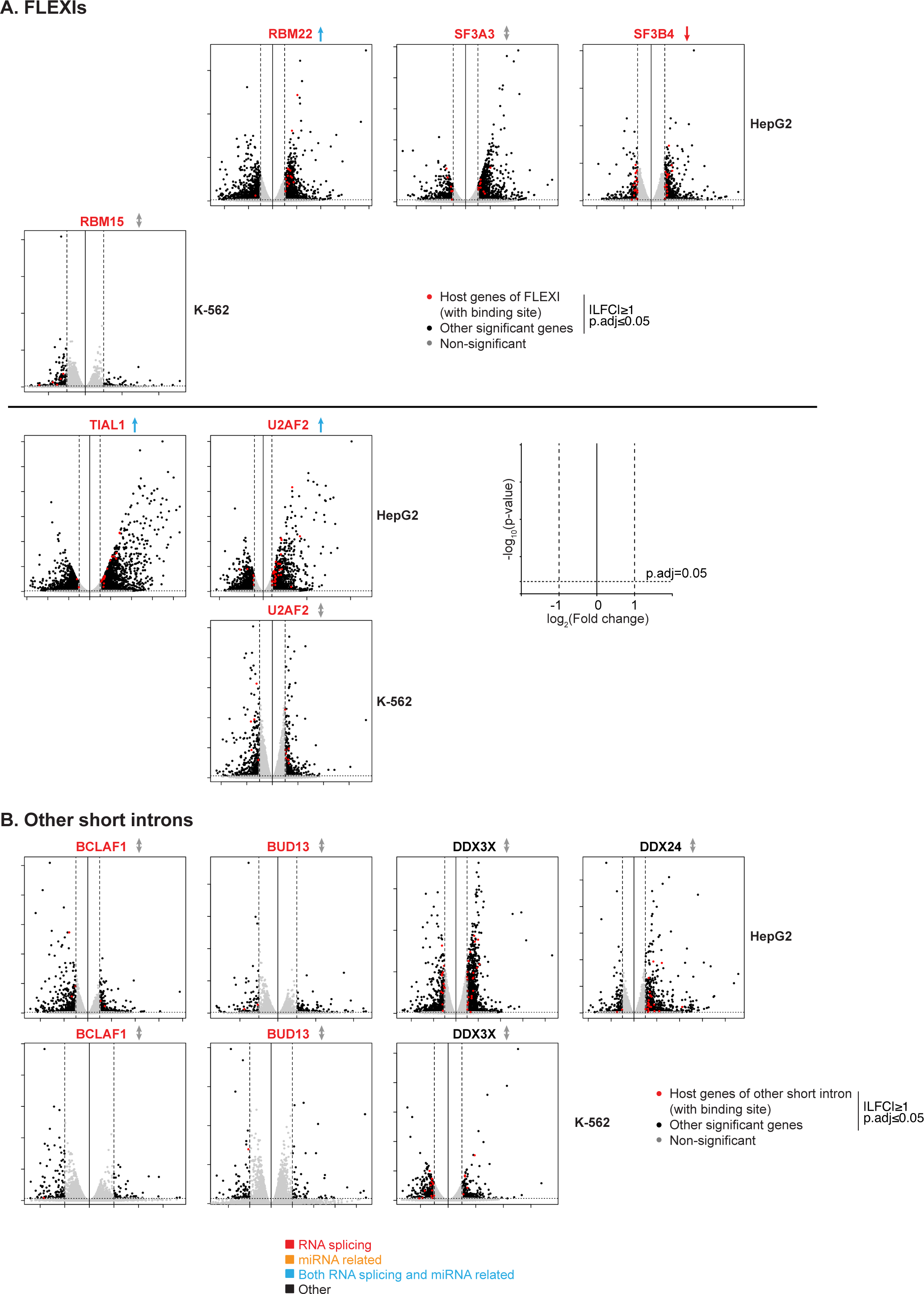
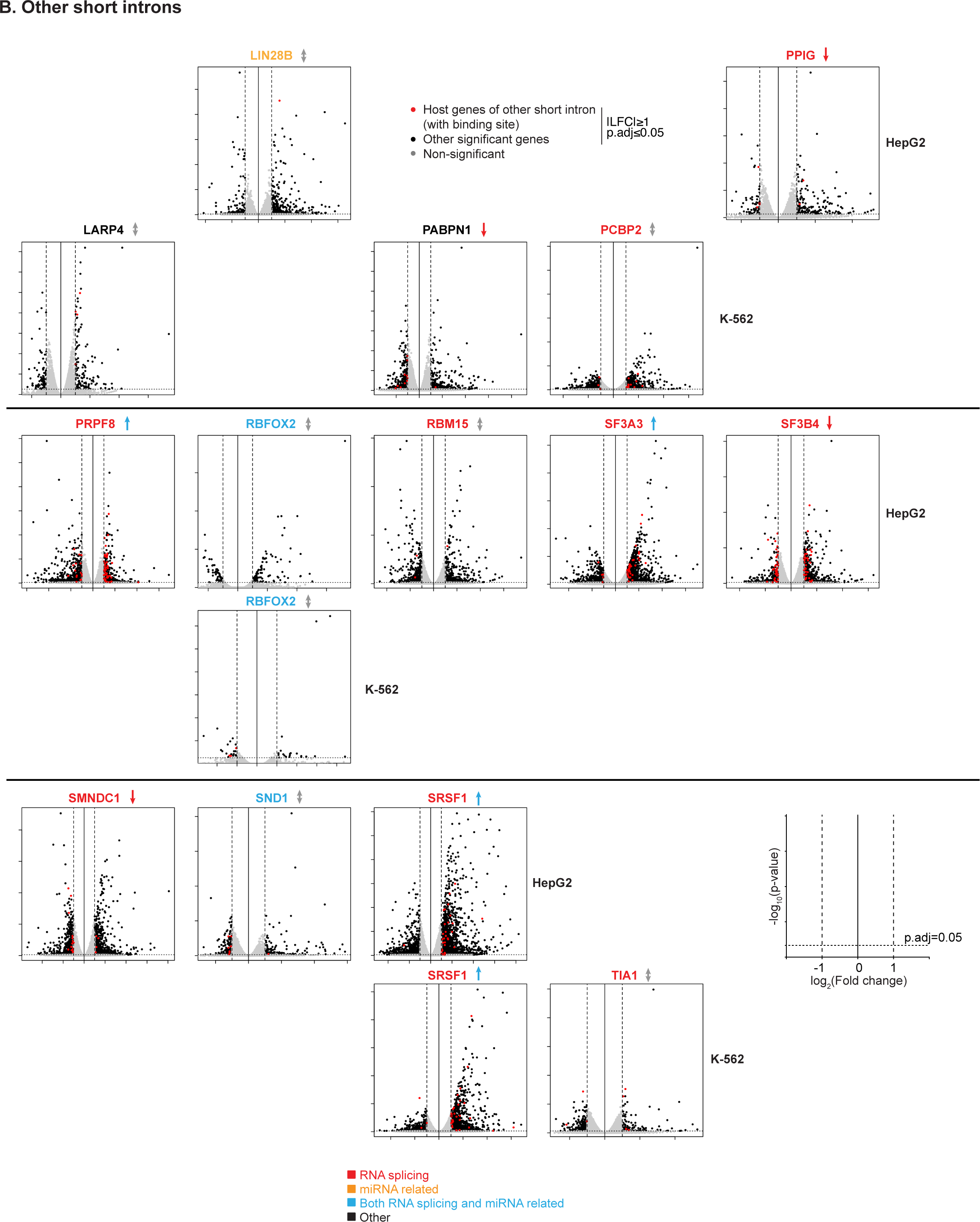
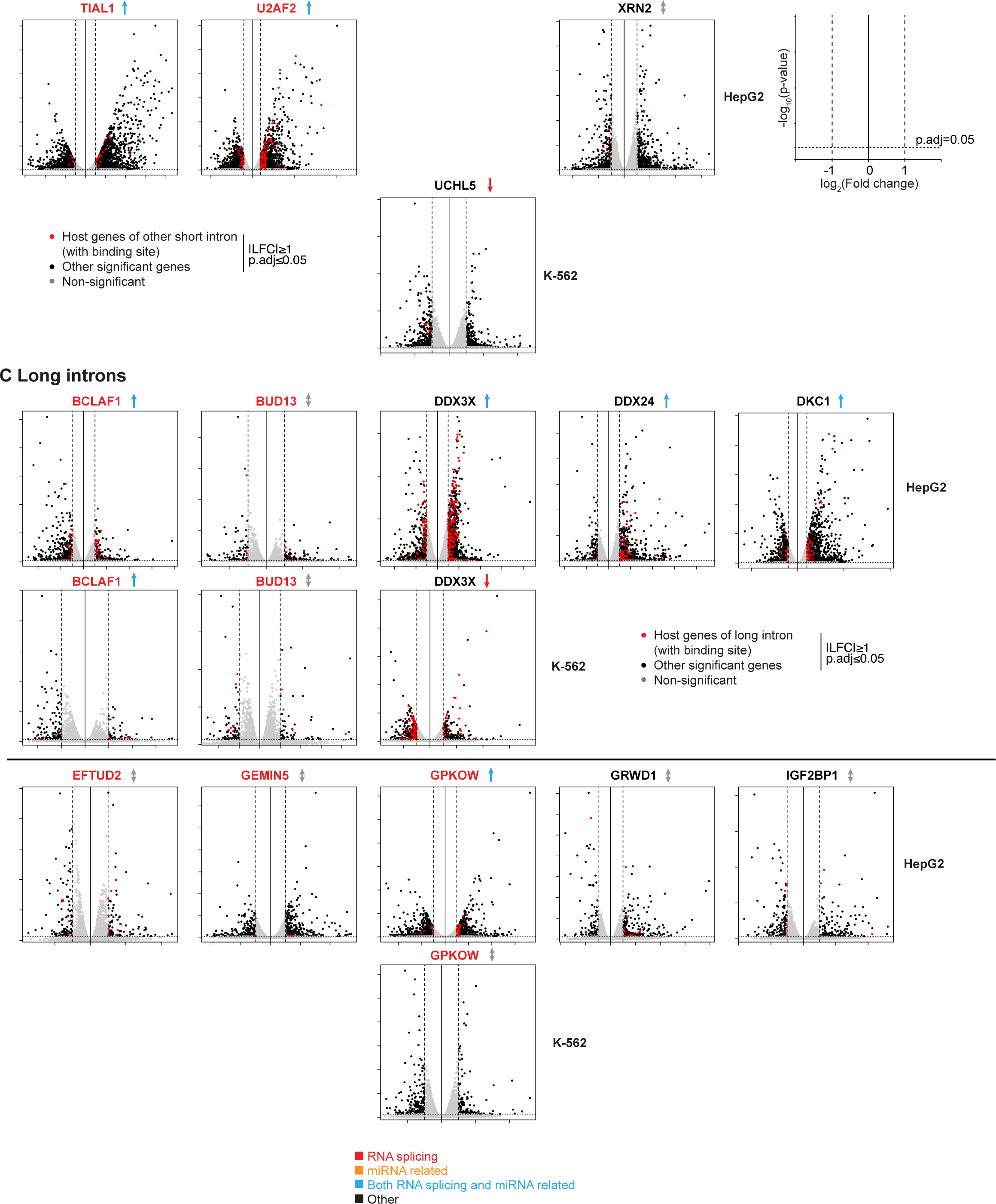
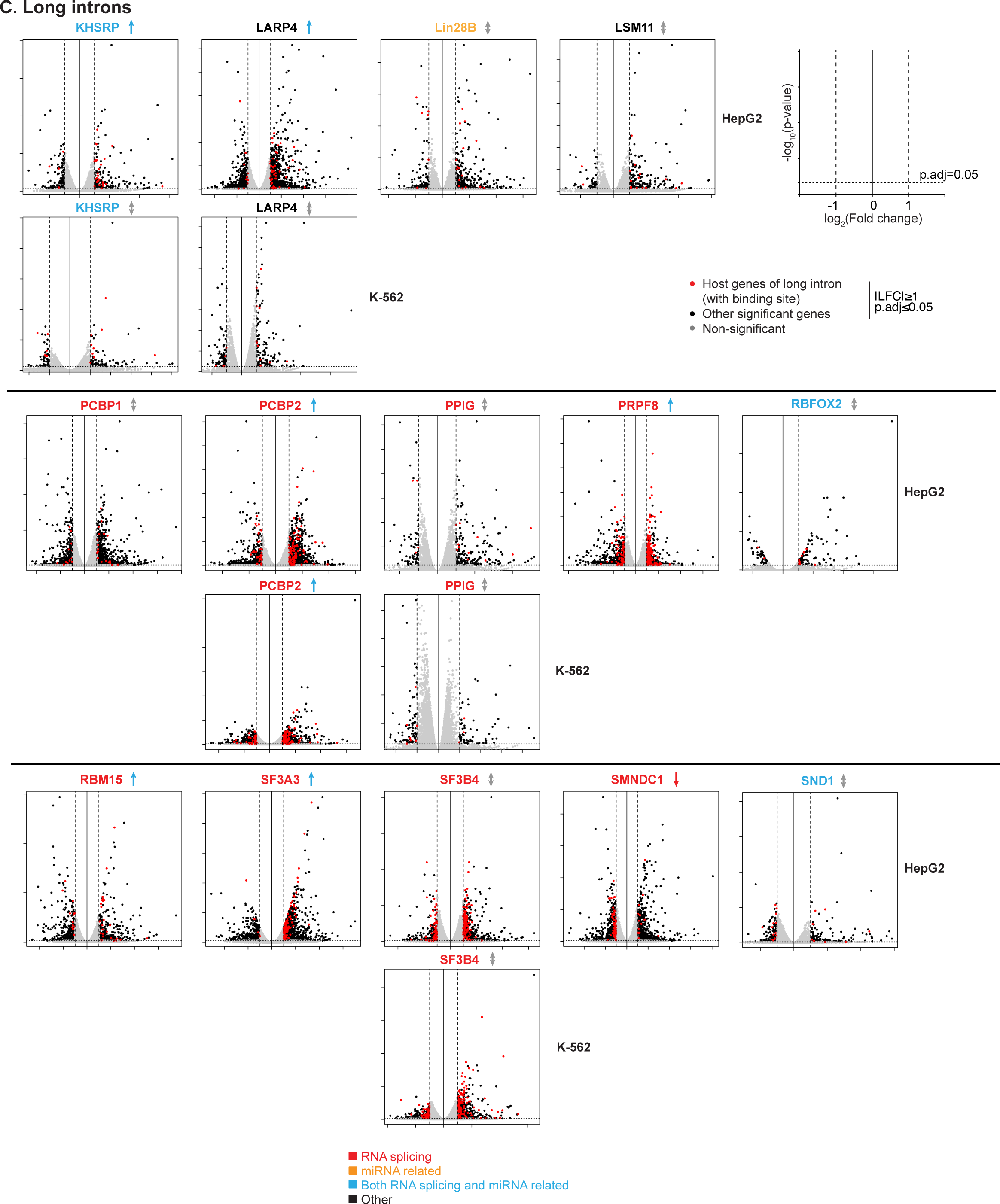
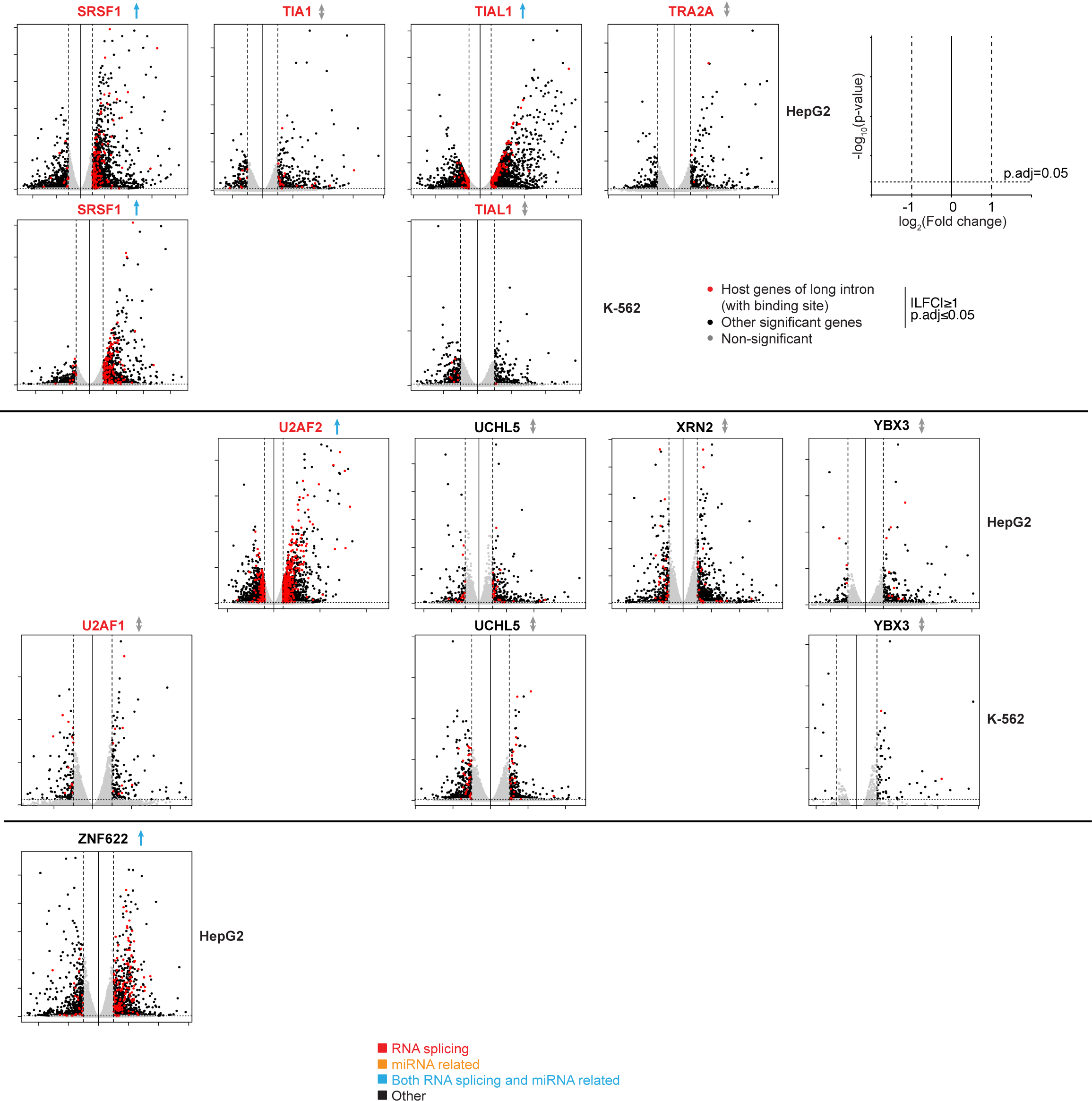
Changes in mRNA levels of host genes of FLEXIs, other short, and long introns with binding sites for different RBPs in RBP knockdown versus control datasets. Volcano plots show -log_10_-transformed adjusted p-values versus log_2_-transformed fold changes for ENSEMBL-annotated genes in ENCODE knockdown versus control datasets for the indicated RBPs in K-562 and HepG2 cells. Host genes of (A) FLEXIs, (B) other short introns, and (C) long introns that contain a binding site for the indicated RBP that have significant expression changes (adjusted p≤0.05, |LFC|≥1) in the knockdown datasets are shown as red dots. Other genes with or without significant expression changes are shown as black or gray dots, respectively. RBP knockdowns that had a significant bias towards increased or decreased mRNA levels in the group of host genes encoding FLEXIs, other short introns, and long introns with a binding site for the RBP are indicated by up (light blue) or down (red) arrows next to the RBP name (pvalue ≤0.05 determined by Fisher’s exact test comparing the ratio of significantly up (log_2_FC>0) or down (log_2_FC<0) regulated host genes containing a FLEXI with a binding site for the RBP to that in all significantly changed FLEXI host genes). RBP knockdowns that resulted in significant changes in mRNA levels from host genes containing a FLEXI, other short introns, and long introns with a binding site for the knocked down RBP, but no significant directional bias, are indicated by a (gray) bi-directional arrow next to the RBP name. Plots are shown only from those RBPs whose knockdown resulted in a significant change (p≤0.05). Datasets that were not available for an RBP in one of the two cell lines were left as a blank space. RBP names are color coded by protein function as shown at the bottom of the Figure.

**Figure S15.**
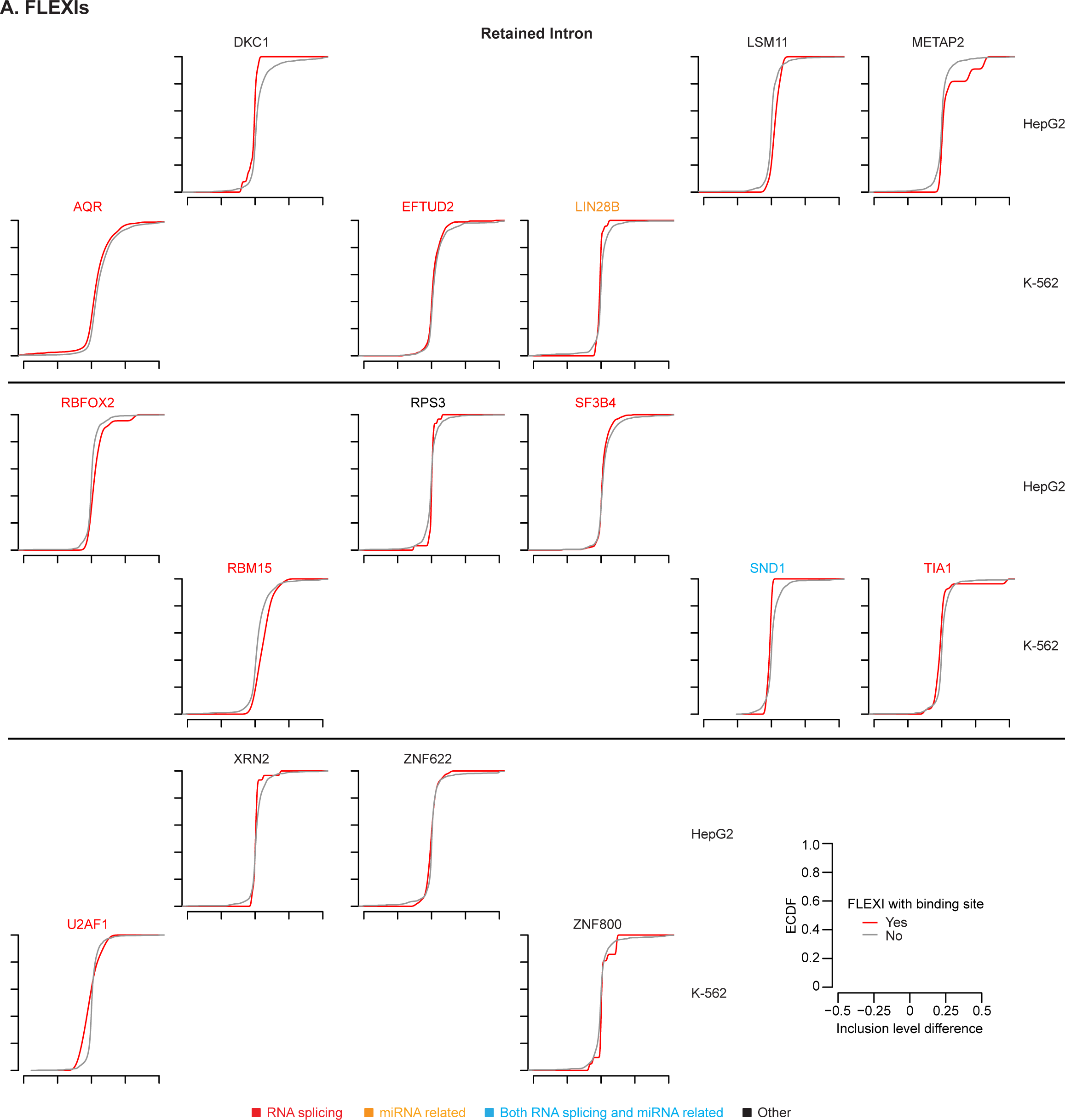
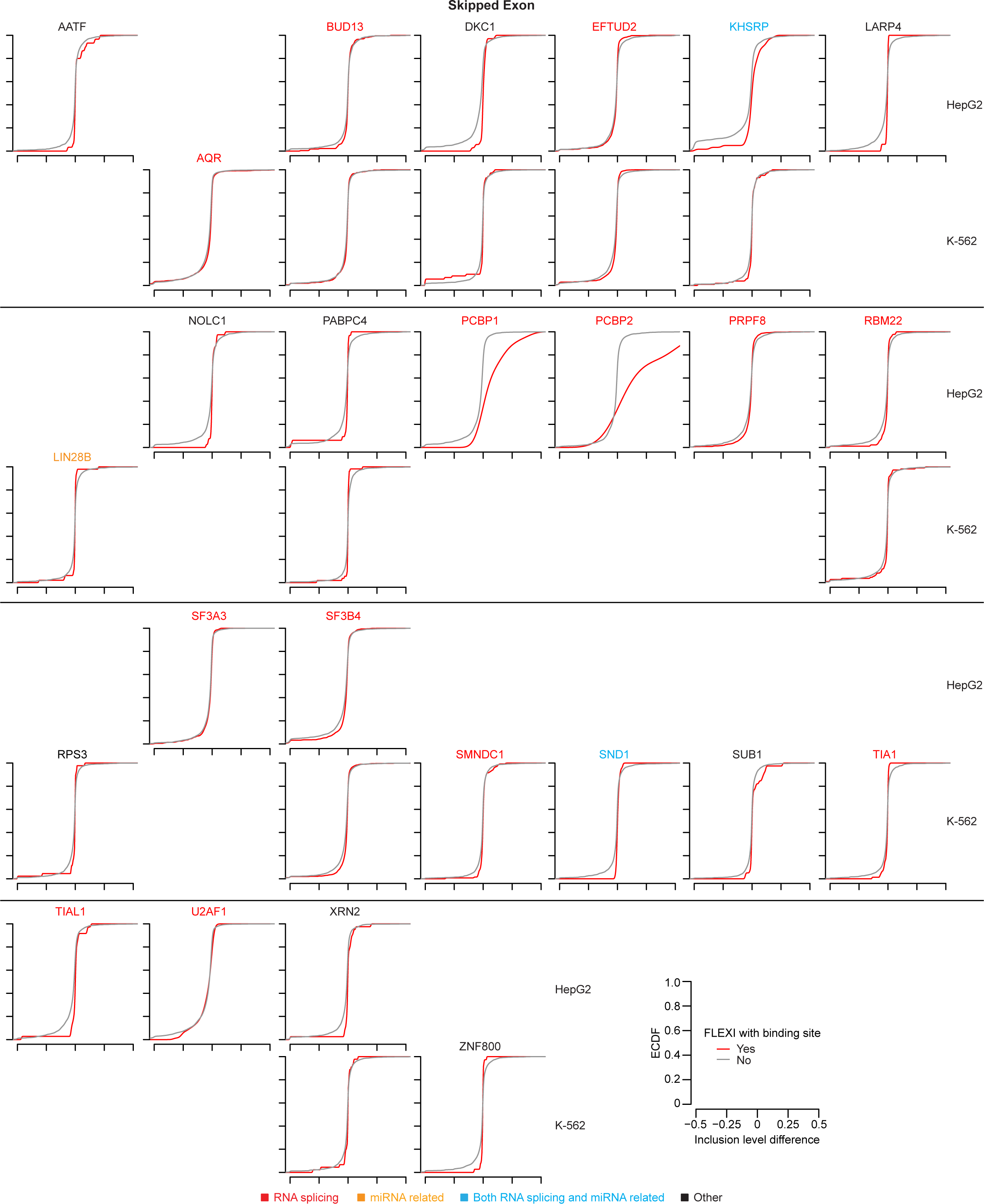
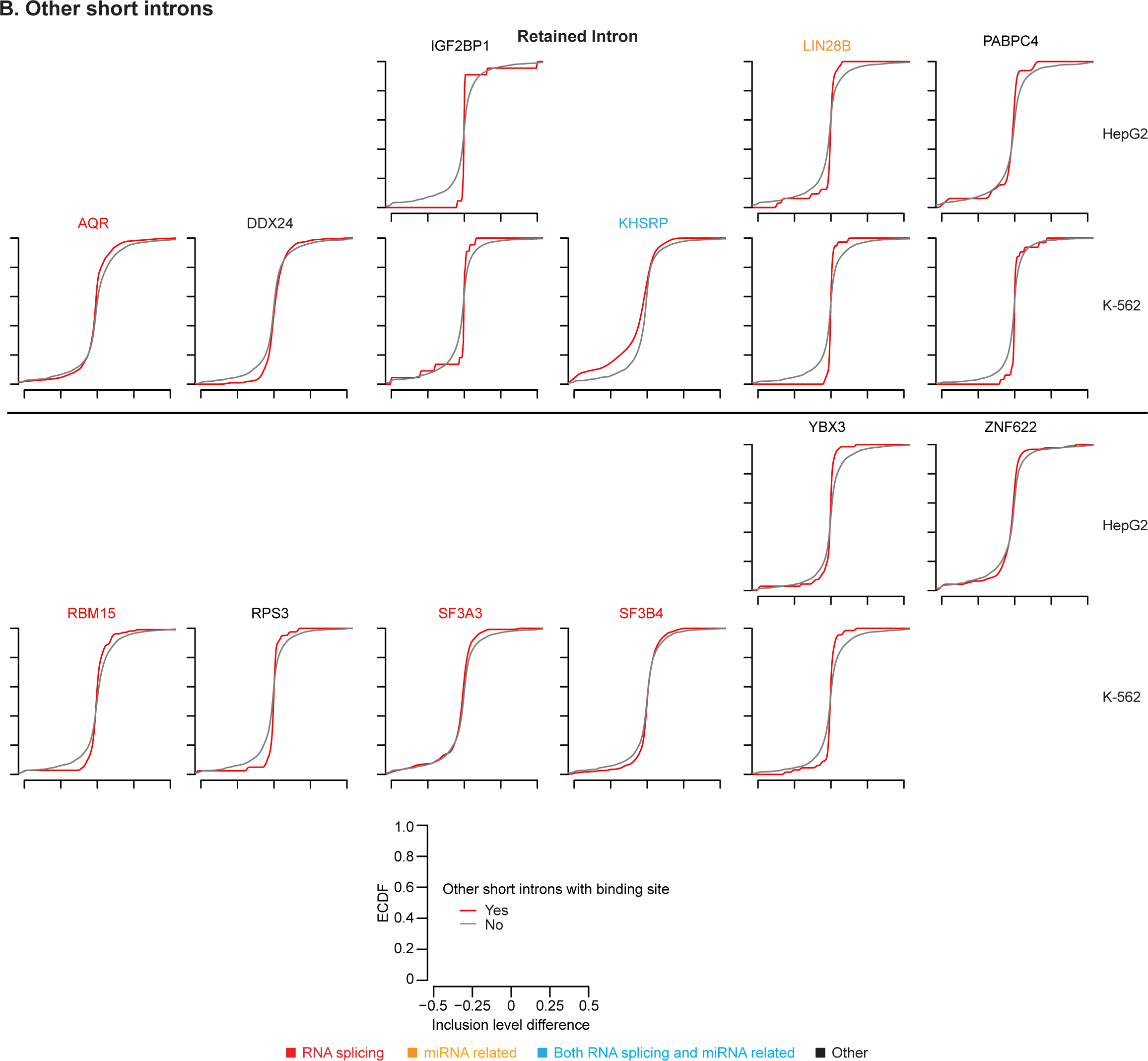
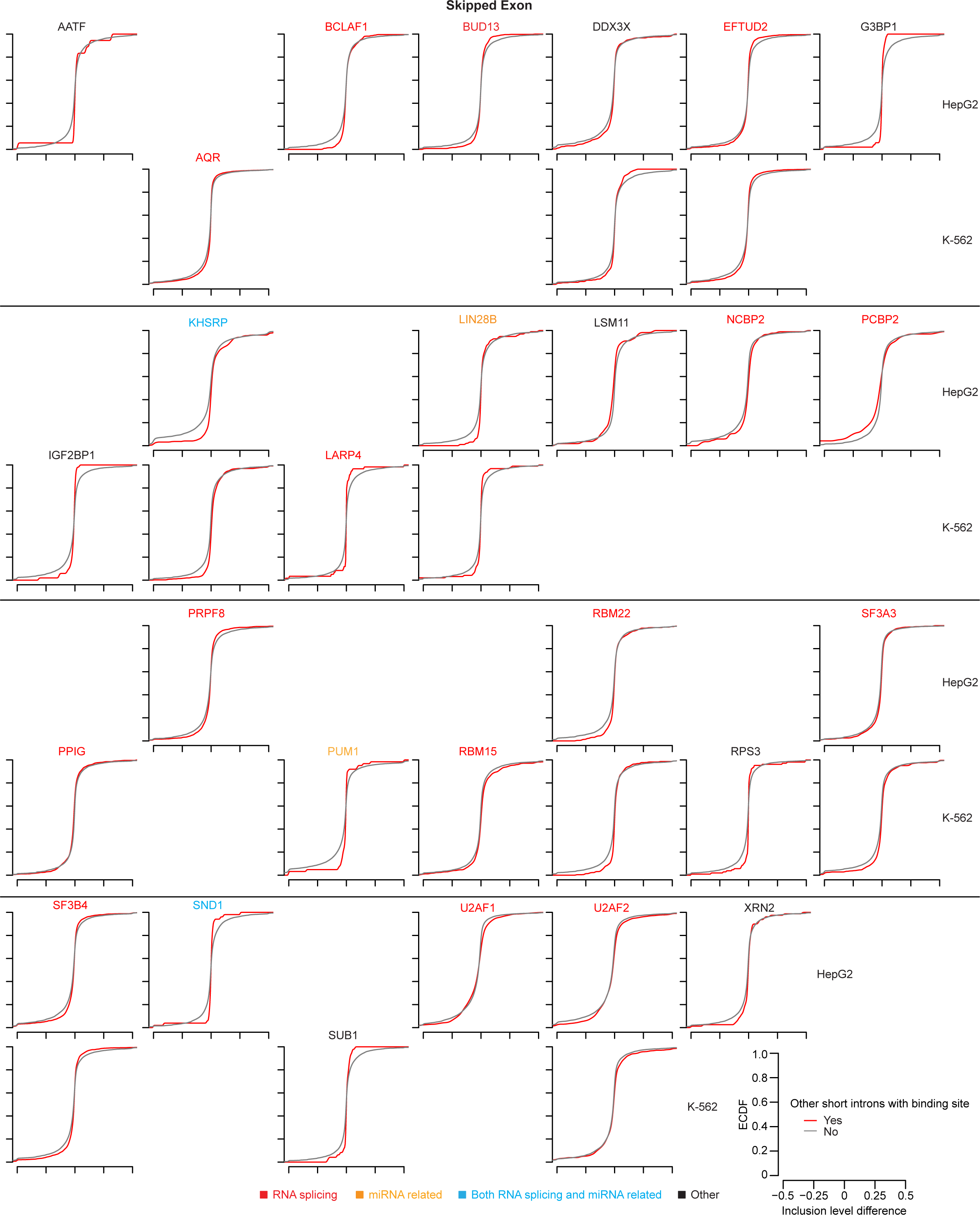
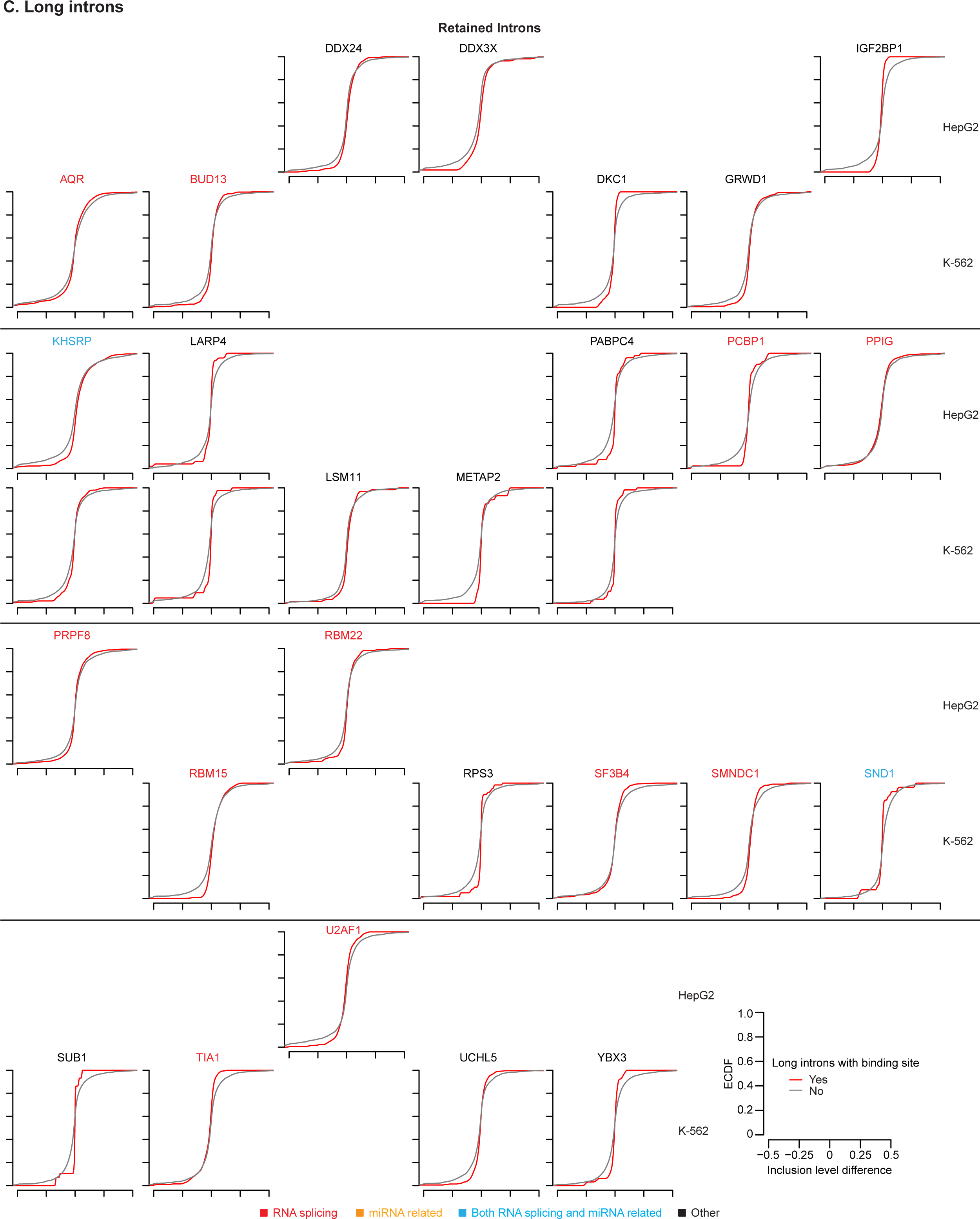
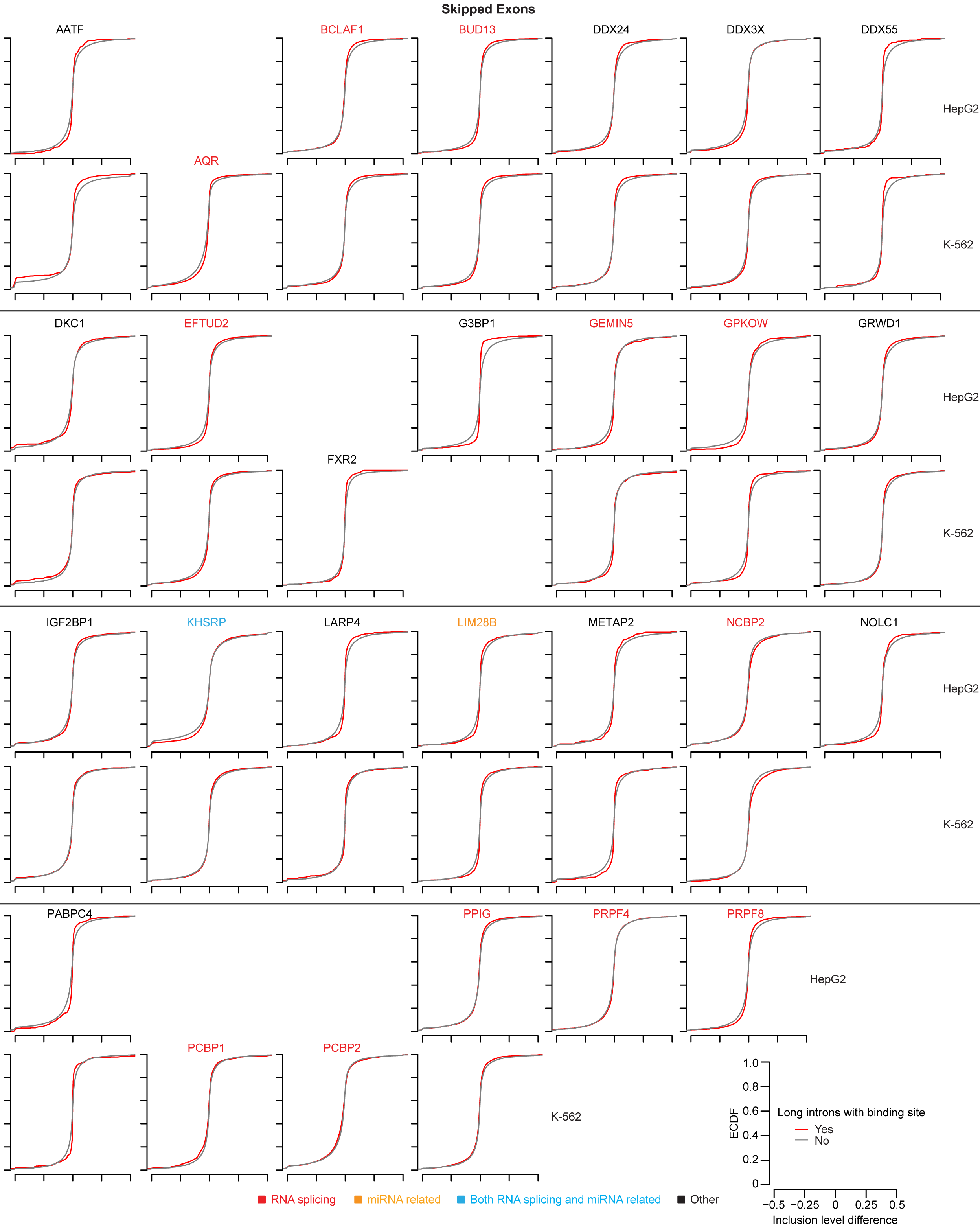
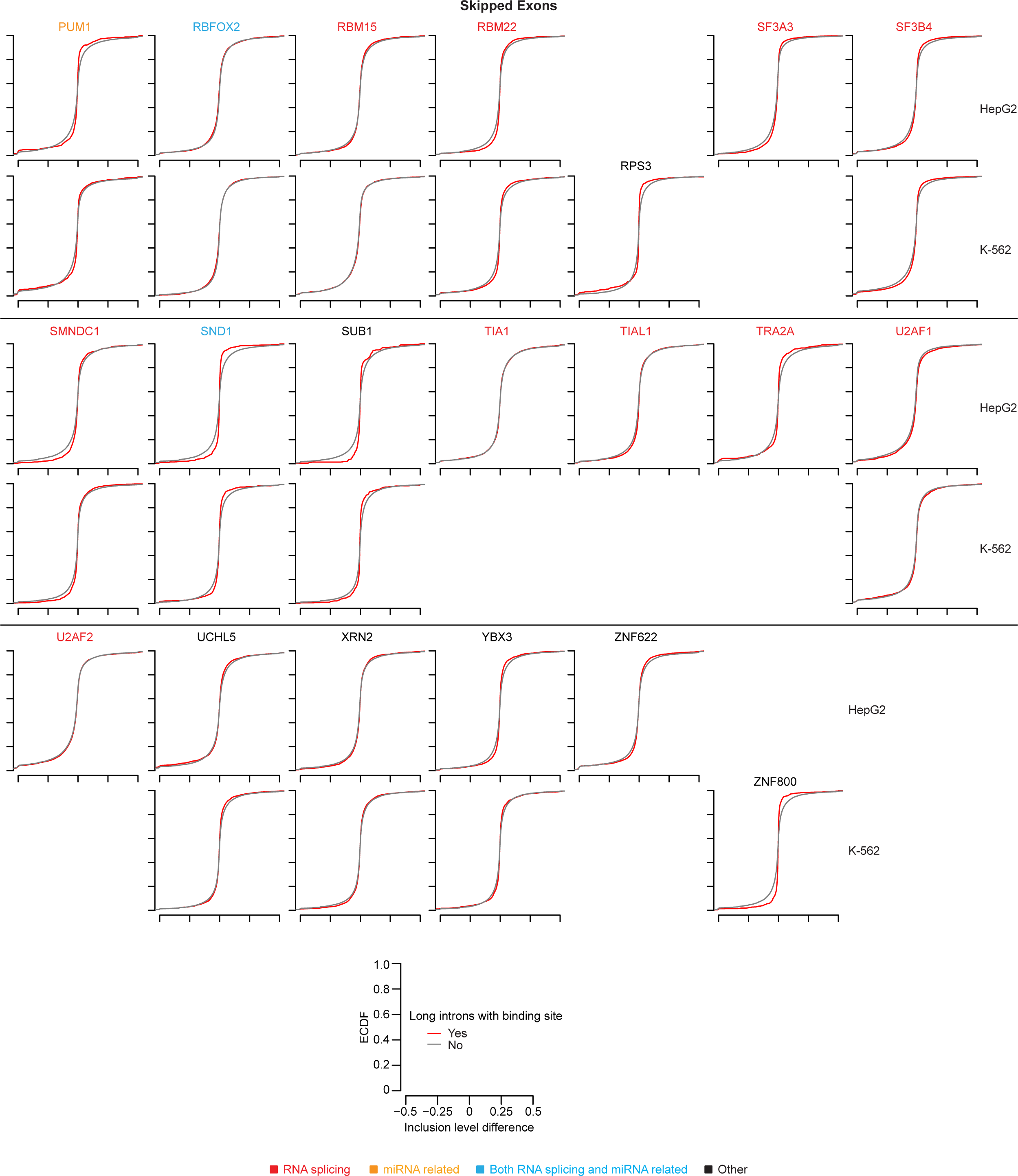
Splicing changes at or adjacent to different classes of introns in RBP-knockdown versus control datasets. Splicing changes for (A) FLEXIs, (B) other short introns, or (C) long introns in ENCODE knockdown datasets for HepG2 and K-562 cells were calculated using rMATS (https://rnaseq-mats.sourceforge.io). Empirical cumulative distribution function (ECDF) plots showing inclusion of retained introns (RI) or skipped exons (SE) adjacent to introns that have (red) or do not have (gray) a CLIP-seq-identified binding site for the indicated RBP in ENCODE knockdown datasets. Red curves shifting to the right or left of the control (gray) indicate an increase or decrease in retained introns and skipped exons as indicated at the top of each set of plots. Statistical significance was calculated by Kolmogorov-Smirnov test. Plots are shown only from those RBPs whose knockdown resulted in a significant change (p≤0.05). Names of RBPs are color coded by protein function as indicated in the Figure. Blank spaces were left for datasets that were not available for an RBP in one of the two cell lines. Axes labels are shown in the key at the bottom right.

**Figure S16.**
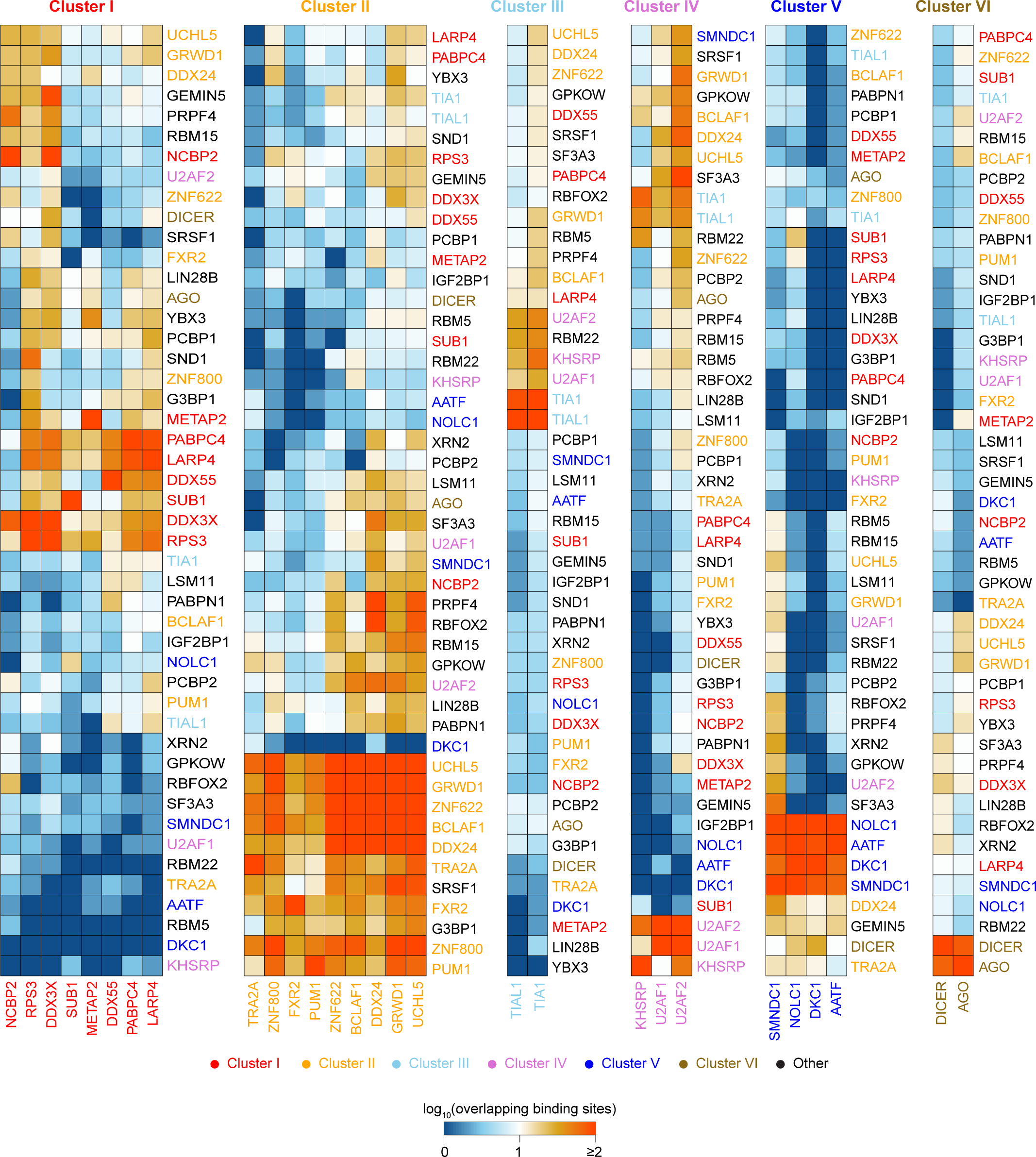
Patterns of overlapping RBP-binding sites in FLEXI RNAs. Heat map of FLEXI RNAs associated with Clusters I-VI that having overlapping binding sites for each of the 47 noncore spliceosomal RBPs that bind ≥30 FLEXIs (columns and rows, respectively). The number of FLEXIs containing overlapping binding sites for each RBP pair (columns and rows) were log_10_ transformed and clustered within boxes color coded by the number of different RBPs that bound the same FLEXIs. Names of RBPs primarily associated with Clusters I-VI at the significance levels used for hierarchical clustering and those not associated with Clusters I-VI (Other) are color coded as shown at the bottom of the Figure.

**Figure S17.**
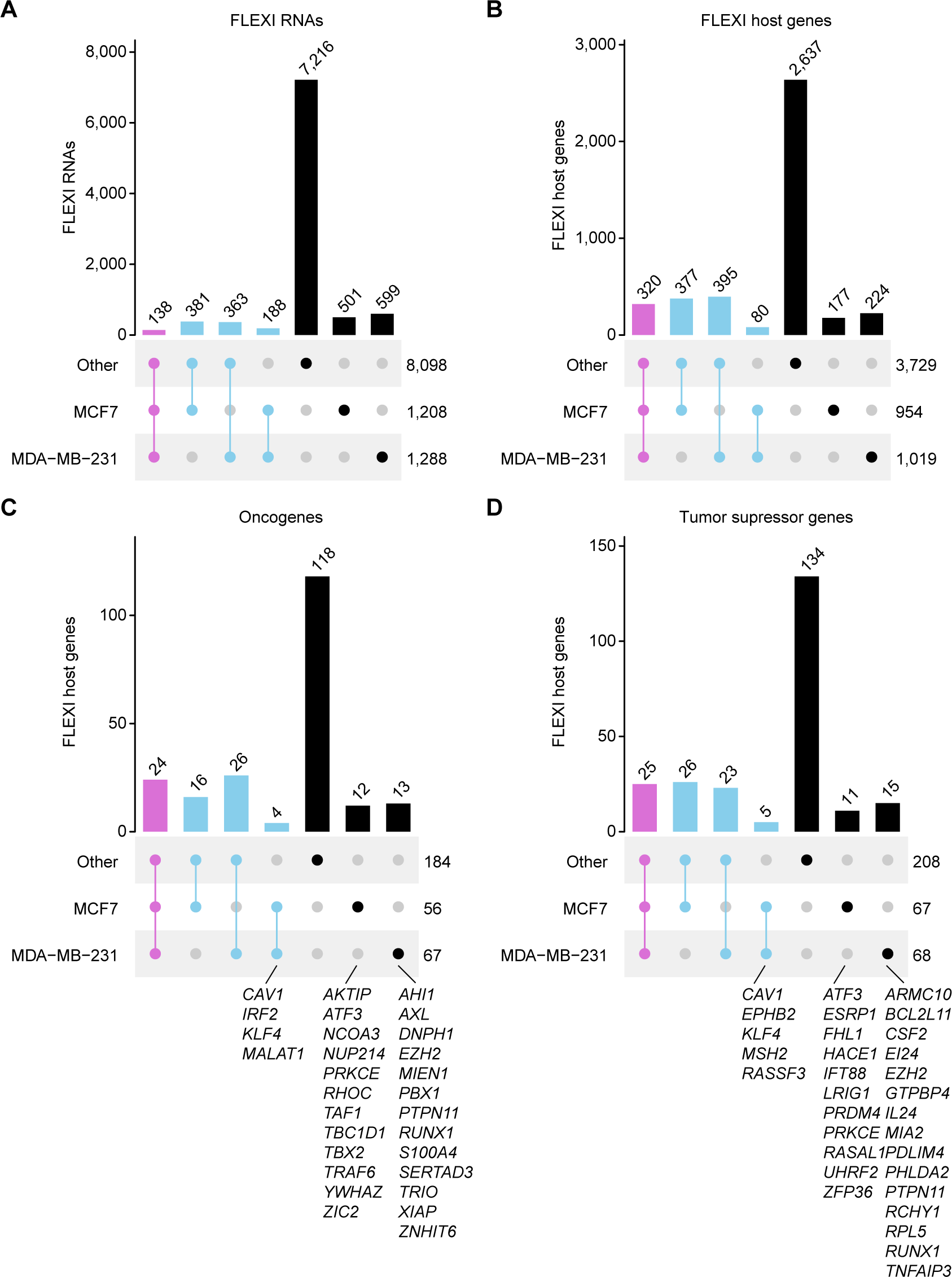
Cell-type specific FLEXI RNAs and host genes in MCF7 and MDA-MB-231 cancer cell lines. UpSet plots showing the distribution of (A) FLEXI RNAs, (B) FLEXI host genes, (C) FLEXI host oncogenes, and (D) FLEXI host tumor suppressor genes in MCF7 cells, MDA-MB-231 cells, and a combined dataset for 4 other cellular RNA samples (HEK-293T, HeLa S4, K-562 and UHRR). FLEXI RNAs from known oncogenes (73) or tumor suppressor genes (74) that were detected only in MCF7 and/or MDA-MB-231 are listed below the plots. ATF3 and KLF4 were annotated as both oncogenes and tumor suppressor genes (73, 74).

**Figure S18.**
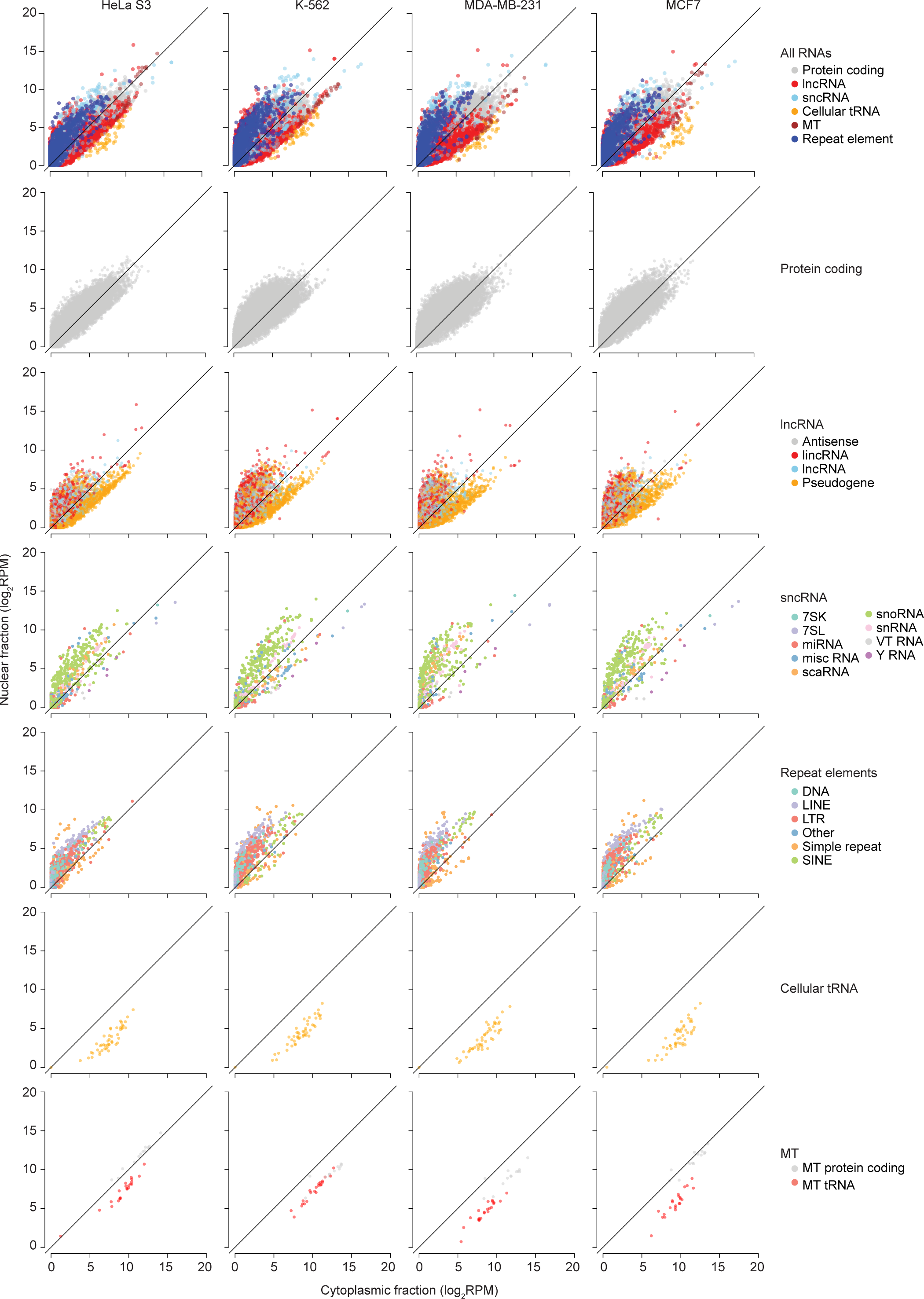
Analysis of RNAs in nuclear and cytoplasmic fractions from cultured cells. Scatter plots comparing enrichment of all and subtypes of cellular RNAs in cytoplasmic and nuclear fractions from HeLa S3, K-562, MDA-MB-231, and MCF7 cells. RNA subtypes are color coded as shown to the right of the scatter plots. Mitochondrial RNAs, tRNAs, Vault RNAs, Y RNA, and 7SL RNA were enriched in the cytoplasmic RNA fraction, while 7SK RNA and snRNAs were enriched in the nuclear RNA fraction. Most snoRNAs (357 to 457 in different cell types) had higher normalized counts in nuclear fractions than in cytoplasmic fractions with only 20-49 snoRNAs having higher normalized counts in cytoplasm fractions.

**Figure S19.**
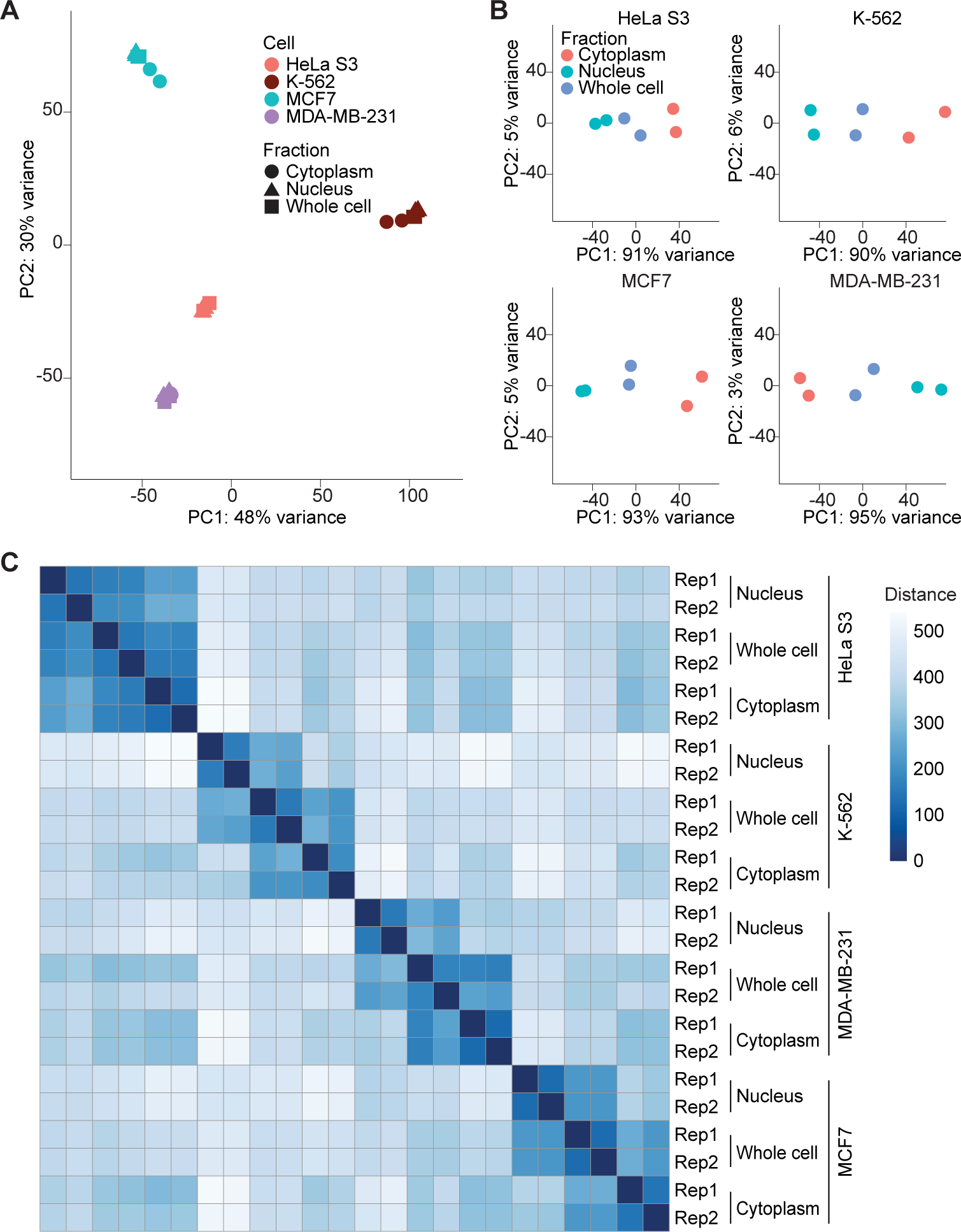
Clustering analysis of RNAs in nuclear and cytoplasmic fractions from cultured cells. (A) and (B) PCA plots for RNAs detected by TGIRT-seq in whole-cell (squares), nuclear fractions (Nucleus; triangles), and cytoplasmic fractions (Cytoplasm; circles) for HeLa S3, K-562, MCF7 and MDA-MB-231 cells color coded as shown in the Figure. (C) Heatmap of Euclidean distance between datasets shaded as being more (darker blue) or less (lighter blue) similar as shown by color scale in the Figure.

**Figure S20.**
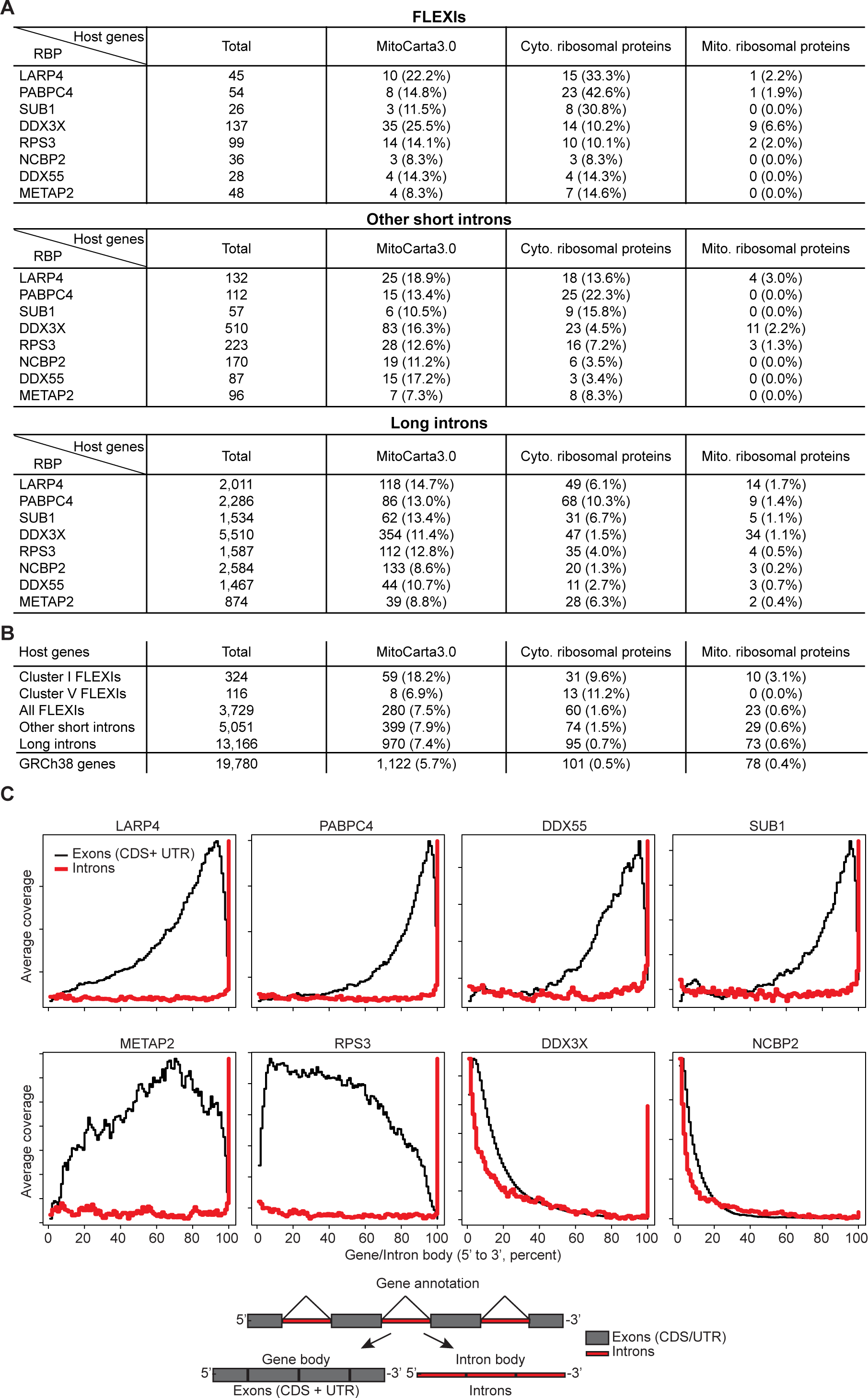
Characterization of RBP-binding sites in intron RNAs encoded by different categories of host genes. (A) Numbers and percentages of host genes encoding mitochondrial proteins annotated in MitoCarta3.0 or cytoplasmic or mitochondrial ribosomal proteins annotated in GRCh38 that contain FLEXIs (top), other short introns (middle), or long introns (bottom) with binding sites for Cluster I RBPs. (B) Numbers and percentages of all host genes for different subsets of introns compared to those for all GRCh38 annotated protein-coding genes. (C) Density plots showing the average coverage of RBP-binding sites across the gene body (Exons CDS + UTRs; black) or intron body (red) calculated using Picard tools normalized by percent length. In the gene body plots, 0% and 100% indicate the 5’ and 3’ ends the mRNAs, respectively. In the intron body plots, 0% indicates the 5’ end of the first intron and 100% indicates the 3’ end of the last intron of the same gene.

## Materials and Methods

### RNA samples and cultured cells

RNA samples used in this work are summarized in Table S1. Universal Human Reference RNA (UHRR) was purchased from Agilent under the product name Quantitative PCR Human Reference Total RNA). HeLa S3 and MCF7 RNAs used for initial FLEXI analysis (Figs. 1 to 9) were purchased from Thermo Fisher. All other cellular RNAs, including those for cell fractionation experiments (Fig. 10), were isolated from cultured cells by using a mirVana miRNA Isolation Kit (Thermo Fisher).

K-562 cells (ATCC CTL-243) were propagated from an ATCC source vial and maintained in Iscove’s Modified Dulbecco’s Medium (IMDM, Thermo Fisher 12440-053) supplemented with 10% Fetal Bovine Serum (FBS; Gemini Bioscience). HEK-293T/17 cells (ATCC CRL-11268) were propagated from an ATCC source vial and maintained in Dulbecco’s Modified Eagle Medium (DMEM, Thermo Fisher 11965-065) supplemented with 10% FBS. HeLa S3 cells (ATCC CCL-2.2) were propagated from an ATCC source vial and were maintained in F-12K (Thermo Fisher 21127-022) supplemented with 10% FBS. MDA-MB-231 cells (ATCC HTB-26) and MCF7 cells (ATCC HTB-22) were obtained from Dr. Blerta Xhelmaçe (University of Texas at Austin) and were maintained in DMEM (Thermo Fisher 11965-092) supplemented with 10% FBS. All cultures were grown in an Eppendorf CellXpert C170i incubator at 37°C with 5% CO_2_ atmosphere.

For RNA isolation, 2-4×10^6^ cells were harvested by centrifugation alone (K-562) or after trypsinization (all other cell lines) at 300 × g for 10 min at 4 °C and washed twice by centrifugation with cold Dulbecco’s Phosphate Buffered Saline (D-PBS; Thermo Fisher 14190-144). Cell pellets were then resuspended in 600 µL of mirVana Lysis Buffer and RNA was isolated according to the kit manufacturer’s protocol with elution in a final volume of 100 µL.

### DNA and RNA oligonucleotides

DNA and RNA oligonucleotides were purchased from Integrated DNA Technologies (IDT). The DNA and RNA oligonucleotides used for TGIRT-seq are shown in Table S4. R2R DNA oligonucleotides with 3’ A, C, G, and T residues were hand-mixed in equimolar amounts prior to annealing to the R2 RNA oligonucleotide to generate RNA/DNA heteroduplexes with a 1-nt 3’- DNA overhang for RNA-seq adapter addition by TGIRT template switching.

The oligonucleotides used as reverse transcriptase or PCR primers for the exonuclease and ddPCR experiments are listed in Table S5. The sequence specificity of all primer sets was confirmed by gel purifying a representative subset of qPCR amplicons and either Sanger sequencing amplicons directly or after TOPO-TA cloning (ThermoFisher). A circular form of the 3I_RAN FLEXI synthetic oligonucleotide control was generated by incubating the 5’ phosphorylated-3I_RAN FLEXI oligonucleotide (500 ng) with T4 RNA Ligase I (10 U; New England Biolabs) for 2 h at 25°C and 2 min at 95°C. The circularized oligonucleotide was cleaned up with an RNA Clean & Concentrator kit (Zymo Research), using 8 sample volumes of ethanol added during purification (denoted 8X ethanol method) to maximize the recovery of small RNAs concurrently with the longer RNAs.

### Exonuclease assay

Unfragmented UHRR or HEK-293T RNAs (5 μg) were DNased with 2 U TURBO DNase (Invitrogen) and purified using an RNA Clean & Concentrator kit (Zymo Research) following the manufacturer’s small-size selection method to separate RNAs ≤200 nt. The input of the small size-selected RNAs were scaled for optimal FLEXI detection in each sample (HEK-293T, 100 ng; UHRR, 70 ng). The size-selected RNAs or synthetic versions of FLEXIs were subjected to exonuclease treatment (1 U Terminator exonuclease (Lucigen), 20 U RNase R (Abcam), 20 U murine RNase Inhibitor (New England Biolabs), 1X RNase R reaction buffer (Abcam), supplemented with additional MgCl_2_ for a final concentration of 1 mm MgCl_2_) or mock-treated in the same reaction conditions without the enzymes, for 1 h at 42°C, with 3 replicates performed for each RNA type. The exonuclease reactions were cleaned up with the RNA Clean & Concentrator kit (8X ethanol method; Zymo Research). An aliquot of this material for synthesized FLEXIs was analyzed on a Novex 10% TBE-Urea polyacrylamide gel (Thermo Fisher) to separate linear and circular RNAs and imaged with SYBR Gold (Thermo Fisher). Reverse transcription was performed by using a gene specific primer (1 μM final concentration) located near the 3’ end of the FLEXI intron with 100 U Maxima H reverse transcriptase (ThermoFisher) in the manufacturer’s buffer and incubated at 25°C for 10 min, 50°C for 30 min, 65°C for 30 min and 85°C for 5 min. qPCR reactions (run in triplicate) were carried out with 1 μL cDNA, 0.3 μμ primers for all primer sets except 4I_JUP which used 0.8 μM, and PowerUp SYBR Green master mix (Life Technologies) using a ViiA 7 Real Time PCR system (Thermo Fisher). Matched control samples omitting the reverse transcriptase (RT-) were performed for every reaction. Mean delta cyclic threshold (CT) values were calculated to obtain a fold change of the RNA levels in the treated compared to the mock-treated samples and converted into a percentage of RNA remaining after exonuclease treatment. In two cases, the exonuclease treatment led to no measurable CT value and in these cases a pseudocount CT value of 41 was added to perform the calculation. Primer sequences are shown in Table S5.

### Droplet digital PCR

Unfragmented HeLa S3, UHRR or HEK-293T RNAs (5 μg) were DNased with 2 U TURBO DNase (Invitrogen) and purified using an RNA Clean & Concentrator kit (Zymo Research) following the manufacturer’s small-size selection method, to separate RNAs ≤200 nt. The inputs of the size-selected RNAs for reverse transcription reactions were scaled to have a similar SNORD14B control amplification when compared with total RNA inputs (HEK-293T and HeLa S3, 5 ng; UHRR, 1 ng). The RNAs were reverse transcribed by using 1 μM final concentration of a gene specific primer (Table S5) located at the 3’ end of the FLEXI intron with 100 U Maxima H Reverse Transcriptase (Thermo Fisher) in the manufacturer’s buffer and incubated at 25°C for 10 min, 50°C for 30 min, 65°C for 30 min and 85°C for 5 min. Matched control samples containing the small size-selected RNA, but omitting the RT (RT-) were performed for every reaction. Droplet digital PCR (ddPCR) was performed on 1 μL cDNA, 0.2 μμ primers for all primer sets except 4I_JUP which used 0.5 μM, and QX200 ddPCR EvaGreen Supermix (Biorad) using the QX200 Droplet Digital PCR system (BioRad). Primer sets (Table S5) were designed to encompass either the FLEXI intron sequence or to span the upstream 5’ exon-FLEXI junction in order to quantify the FLEXI RNA compared to any precursor or mature mRNA with retained FLEXI introns. The same gene-specific-generated cDNA was used as the PCR template for quantifying both the FLEXI and 5’ exon-FLEXI junctions. Due to the short length of the FLEXI sequences, primer design was constrained to the best available primer location that maximized length and sequence specificity. As a result of the primer constraints, some primers amplified a smaller region of the FLEXI sequence than the TGIRT-seq criteria (*i.e.,* 5’ and 3’ ends within 3 nt of the annotated splice sites), which could contribute to differences between ddPCR and TGIRT-seq quantification. 5’ exon-FLEXI junction primer pairs used a different reverse primer than the FLEXI, that was inset closer to the 5’ exon-FLEXI junction, so that the size of the amplicon would be more similar to the size of the compared FLEXI amplicon. As a consequence of these constraints, some primer pairs amplified background primer dimer artifacts in addition to verified products. For each reaction replicate, any RT-signal was subtracted, and three background-subtracted reaction replicates were averaged for each primer pair. The majority of the 5’ exon-FLEXI junction reactions either gave no product or amplified non-specific products. However, a few 5’ exon-FLEXI junction reactions produced validated products sequenced by direct Sanger sequencing and/or TOPO-TA cloning and Sanger sequencing of gel extracted, matched qPCR amplicons (HEK-293T, *RPS2*; HeLa S3, *ACTB*; UHRR, *POLG* and *ACTB*). These sequence-validated 5’ exon-FLEXI junction ddPCR signals were subtracted from the FLEXI ddPCR signals to obtain the final ddPCR estimate for abundance of the FLEXI. FLEXI copies per reaction were then normalized to the controls performed in parallel: U7 (chosen for similar abundance to FLEXIs), SNORD14B (75), and SNORD44 (76). Finally, the normalized FLEXI copies per reaction were converted to copy number per cell estimates by multiplying by respective control values generated by the linear regression model without or with TGIRT-seq bias correction.

### Cell fractionation

Cell cultures were grown to ∼80% confluency (HeLa S3, MDA-MB-231 and MCF7) or to a density of 9×10^5^ cells/mL (K-562) in T-75 flasks. To prepare cell pellets, adherent cells were either (1: HeLa S3) washed 3× with cold D-PBS and then scraped in D-PBS and pelleted by centrifugation at 1,000xg for 5 min at 4°C in an Eppendorf FA-24×2-PTFE rotor; or (2: MDA-MB-231 and MCF7) trypsinized and collected by centrifugation at 1,000 RPM for 5 min at room temperature in a Centra CL2 centrifuge with Aerocarrier rotor (Thermo Fisher) and then resuspended in cold D-PBS and washed 3× by centrifugation at 1,000xg for 5 min at 4°C in an Eppendorf FA-24×2-PTFE rotor. Suspension cells (K-562) were collected by centrifugation at 1,000 RPM for 5 min at room temperature in a Centra CL2 centrifuge with Aerocarrier rotor and then placed on ice and resuspended in cold D-PBS and washed 3× by centrifugation at 1,000xg for 5 min 4°C in an Eppendorf FA-24×2-PTFE rotor.

Cellular fractionation was performed as described in Rio *et al*. (77) . Cell pellets were resuspended in 1.0 mL of cell disruption buffer and then incubated for 10 min on ice and monitored for swelling by light microscopy. Swollen cells were transferred to a baked (400°C, 4 h) 2.0-mL glass Dounce homogenizer and broken with 10 strokes of a Type B pestle. Cell breakage was confirmed by microscopy. Cell homogenates were transferred to 1.7-mL tubes and brought to 0.1% Triton X-100 by the addition of 10% (w/v) Triton X-100 (Sigma). Samples were then centrifuged immediately at 1,500 × g for 5 min at 4°C in an Eppendorf FA-24×2-PTFE rotor.

Nuclear fraction pellets and cytoplasmic fraction supernatants were transferred to fresh tubes and placed on ice. RNA was extracted by using a mirVana RNA extraction kit following the manufacturer’s instructions. The nuclear fraction pellets were resuspended in 600 µL of kit lysis buffer and processed on a mirVana column. The cytoplasmic samples were split such that 200 µL of cytoplasm was solubilized in 400 µL of kit lysis buffer and then combined and processed together on the same column. RNA samples were evaluated on an Agilent 2100 Bioanalyzer with a Eukaryotic Total RNA Pico kit using the mRNA assay. RNAs were then processed for TGIRT-seq library construction as described below.

### Thermostable Group II Intron Reverse Transcriptase sequencing (TGIRT-seq)

TGIRT-seq libraries were prepared from unfragmented RNA as described (21, 22). To remove residual DNA, UHRR and HeLa S3 RNAs (1 μg) were incubated with 20 U exonuclease I (Lucigen) and 2 U Baseline-ZERO DNase (Lucigen) in Baseline-ZERO DNase Buffer for 30 min at 37 °C, and K-562, MCF7, MDA-MB-231 and HEK-293T RNAs (5 μg) were incubated with 2 U TURBO DNase for 30 min at 37 °C (Thermo Fisher). After DNase digestion, RNA was cleaned up with an RNA Clean & Concentrator kit (8X ethanol method; Zymo Research). The eluted RNAs were ribodepleted by using the rRNA removal section of a TruSeq Stranded Total RNA Library Prep Human/Mouse/Rat kit (Illumina), with the supernatant from the magnetic-bead separation cleaned up by using a Zymo RNA Clean & Concentrator kit with 8X ethanol. After checking RNA concentration and length by using an Agilent 2100 Bioanalyzer with a 6000 RNA Pico chip, RNAs were aliquoted into ∼20 ng portions and stored at -80 °C until use. TGIRT-seq libraries were prepared as described (21, 22) using 20-50 ng of ribodepleted unfragmented cellular RNAs. Template-switching and reverse transcription reactions were done with 1 μM TGIRT-III (InGex, currently available upon request from the Lambowitz lab) and 100 nM pre-annealed R2 RNA/R2R DNA starter duplex in 20 μL of reaction medium containing 450 mM NaCl, 5 mM MgCl_2_, 20 mM Tris-HCl, pH 7.5 and 5 mM DTT. Reactions were set up with all components except dNTPs, pre-incubated for 30 min at room temperature, a step that increases the efficiency of RNA-seq adapter addition by TGIRT template switching, and initiated by adding dNTPs (final concentrations 1 mM each of dATP, dCTP, dGTP, and dTTP). The reactions were incubated for 15 min at 60°C and then terminated by adding 1 μL 5 M NaOH to degrade RNA and heating at 95 °C for 5 min followed by neutralization with 1 μL 5 M HCl and one round of MinElute column clean-up (Qiagen). The R1R DNA adapter was adenylated by using a 5’ DNA Adenylation kit (New England Biolabs) and then ligated to the 3’ end of the cDNA by using Thermostable 5’ App DNA/RNA Ligase (New England Biolabs) for 2 h at 65°C. The ligated products were purified by using a MinElute Reaction Cleanup Kit and amplified by PCR with Phusion High-Fidelity DNA polymerase (Thermo Fisher; denaturation at 98°C for 5 s followed by 12 cycles of 98°C for 5 s, 60°C for 10 s, and 72°C for 15 s and then held at 4°C until removed. The PCR products were cleaned up by using Agencourt AMPure XP beads (1.4X volume; Beckman Coulter) and sequenced on an Illumina NextSeq 500 to obtain 2 × 75 nt pairedend reads or on an Illumina NovaSeq 6000 to obtain 2 × 150 nt paired-end reads at the Genome Sequence and Analysis Facility of the University of Texas at Austin.

### Sequencing data processing

Illumina TruSeq adapters and PCR primer sequences were trimmed from the reads with Cutadapt v2.8 (https://github.com/marcelm/cutadapt, sequencing quality score cut-off at 20; p-value <0.01) and reads <15 nt after trimming were discarded. To minimize mismapping, we used a sequential mapping strategy. First, reads were mapped to the human mitochondrial genome (Ensembl GRCh38 Release 93) and the *Escherichia coli* genome (GeneBank: NC_000913) using HISAT2 v2.1.0 (http://daehwankimlab.github.io/hisat2/) with customized settings (-k 10 --rfg 1,3 --rdg 1,3 --mp 4,2 --no-mixed --no-discordant --no-spliced-alignment) to filter out reads derived from mitochondrial and *E. coli* RNAs (denoted Pass 1). Unmapped read from Pass1 were then mapped to a collection of customized reference sequences for human sncRNAs (miRNA, tRNA, Y RNA, Vault RNA, 7SL RNA, 7SK RNA) and rRNAs (2.2-kb 5S rRNA repeats from the 5S rRNA cluster on chromosome 1 (1q42, GeneBank: X12811) and 43-kb 45S rRNA containing 5.8S, 18S and 28S rRNAs from clusters on chromosomes 13,14,15, 21, and 22 (GeneBank: U13369), using HISAT2 with the following settings -k 20 --rdg 1,3 --rfg 1,3 --mp 2,1 --no-mixed --no-discordant --no-spliced-alignment --norc (denoted Pass 2). Unmapped reads from Pass 2 were then mapped to the human genome reference sequence (Ensembl GRCh38 Release 93) using HISAT2 with settings optimized for non-spliced mapping (-k 10 --rdg 1,3 --rfg 1,3 --mp 4,2 --no-mixed --no-discordant --no-spliced-alignment) (denoted Pass 3) followed by splice aware mapping (-k 10 --rdg 1,3 --rfg 1,3 --mp 4,2 --no-mixed --no-discordant --dta) (denoted Pass 4). Finally, the remaining unmapped reads were mapped to Ensembl GRCh38 Release 93 by Bowtie 2 v2.2.5 (https://github.com/BenLangmead/bowtie2) using local alignment (-k 10 --rdg 1,3 --rfg 1,3 --mp 4 --ma 1 --no-mixed --no-discordant --very-sensitive-local) to improve mapping rates for reads containing post-transcriptionally added 5’ or 3’ nucleotides (poly(A) or poly(U)), short untrimmed adapter sequences, and non-templated nucleotides added to the 3’ end of the cDNAs by TGIRT-III during TGIRT-seq library preparation (denoted Pass 5). For reads that map to multiple genomic loci with the same mapping score in Passes 3 to 5, the alignment with the shortest distance between the two paired ends (*i.e.*, the shortest read span) was selected. In the case of ties (*i.e.*, reads with the same mapping score and read span), reads mapping to a chromosome were selected over reads mapping to scaffold sequences, and in other cases, the read was assigned randomly to one of the tied choices. The filtered multiply mapped reads were then combined with the uniquely mapped reads from Passes 3-5 by using SAMtools v1.10 (https://github.com/samtools/samtools).

To generate counts for different genomic features, the mapped reads were intersected with Ensembl GRCh38 Release 93 gene annotations plus the *RNY5* gene and its 10 pseudogenes, which are not annotated in this release. Coverage of each feature was calculated by BEDTools v2.29.2 (https://github.com/arq5x/bedtools2). To avoid miscounting reads with embedded sncRNAs that were not filtered out in Pass 2 (*e.g.,* snoRNAs), reads were first intersected with sncRNA annotations and the remaining reads were then intersected with the annotations for protein-coding gene RNAs, lincRNAs, antisense RNAs, and other lncRNAs to get the read count for each annotated feature. Coverage plots and read alignments were created by using Integrative Genomics Viewer v2.6.2 (IGV, https://igv.org). Genes with >100 mapped reads were down sampled to 100 mapped reads in IGV for visualization. Down sampling of reads mapping to FLEXI host genes was conducted by randomly picking mapped reads to match the read depth of FLEXI RNA reads.

### Identification and characterization of FLEXIs

To identify short introns that could give rise to FLEXI RNAs, intron annotations were extracted from Ensemble GRCh38 Release 93 gene annotations using a customized script and filtered to remove introns >300 nt as well as duplicate intron annotations from different mRNA isoforms. Mapped reads were then intersected with short intron annotations using BEDTools, and read pairs (Read 1 and Read 2) ending at or within 3 nucleotides of annotated 5’- and 3’-splice sites were identified as corresponding to FLEXI RNAs. Exon-junction reads were defined as those in which one of the paired-end reads mapped within the intron and the other within a flanking exon.

UpSet plots of FLEXI RNAs from different sample types were plotted by using the ComplexHeatmap package v2.2.0 in R (https://jokergoo.github.io/ComplexHeatmap-reference/book/). For UpSet plots of FLEXI host genes, FLEXI RNAs were aggregated by Ensemble ID, and different FLEXI RNAs from the same gene were combined into one entry. Density distribution plots and scatter plots were plotted by using R.

FLEXI RNAs corresponding to annotated mirtrons or agotrons were identified by intersecting FLEXI RNA coordinates with those of annotated mirtrons (58) and agotrons (19). FLEXI RNAs containing embedded snoRNAs were identified by intersecting the FLEXI RNA coordinates with those of annotated snoRNAs and scaRNAs from Ensembl GRCh38 annotations.

5’- and 3’-splice sites (SS) and branch-point (BP) consensus sequences of human U2- and U12-type spliceosomal introns were obtained from previous publications (29, 30). FLEXI RNAs corresponding to U12-type introns were identified by searching for: (i) FLEXI RNAs with AU-AC ends and (ii) the 5’-splice site consensus sequence of U12-type introns with GU-AG ends (29) using FIMO (https://meme-suite.org/meme/tools/fimo) with the following settings: FIMO --text - -norc <GU_AG_U12_5SS motif file> <SEQUENCE file>. Splice-site consensus sequences of FLEXI RNAs were calculated from nucleotide frequencies at the 5’ and 3’ ends of the introns. The U2-type intron branch-point consensus sequence of intron RNAs was identified by searching for U2-type branch-point consensus using FIMO. The U12-type intron branch-point consensus was determined by searching for motifs enriched within 40 nt of the 3’ end of the introns using MEME (https://meme-suite.org/meme/tools/meme) with settings: meme <SEQUENCE file> -rna - oc <OUTPUT folder> -mod anr -nmotifs 100 -minw 6 -minsites 100 -markov order 1 -evt 0.05. The branch-point consensus sequence of U12-type FLEXI RNAs (2 with AU-AC ends and 34 with GU-AG matching the 5’ sequence of GU-AG U12-type introns) was determined by manual sequence alignment and calculation of nucleotide frequencies. Motif logos were plotted from the nucleotide frequency tables of each motif using Ceqlogo from the MEME suite (https://meme-suite.org/meme/doc/ceqlogo.html).

The length and GC content of each FLEXI was calculated from its GRCh38-annotated intron sequence. Minimum free energy (MFE) was calculated for the most stable secondary structure predicted by RNAfold for each FLEXI RNA. The distribution of length, GC content and MFE of FLEXI subsets compare to those of all other detected FLEXIs were compared by Kolmogorov– Smirnov (KS) test. To test if the significance in distribution was due to the small sample size in FLEXI subsets, those with p-values <0.01 (denoted test subset) were further tested by 1,000 Monte-Carlo simulations. In each simulation, a random selected FLEXI subset with the same number of FLEXIs as the test subset was compared to all other FLEXIs by the same KS test. If more than 50 of the 1,000 simulations (≥95%) yield a calculated p<0.01, the significance in the original test subset was considered to be a false hit.

### PCA, t-SNE, and ZINB-WaVE analysis

PCA analysis of cell-type specific FLEXI RNA profiles in replicate cellular RNA datasets was plotted using R, and PCA initialized t-SNE and ZINB-WaVE analyses of these datasets were plotted using the Rtsne (https://github.com/jkrijthe/Rtsne) and zinbwave (https://github.com/drisso/zinbwave) packages in R. Batch effect correction was done by using ZINB-WaVE to include the batch information as a sample-level covariate. The normalized counts outputted by ZINB-WaVE were used for PCA analysis, and the *W* matrix outputted by ZINB-WaVE was used for visualization directly or by t-SNE.

### RBP-binding site analysis

FLEXI RNAs containing RBP-binding sites were identified by intersecting the FLEXI RNA coordinates with ENCODE annotated binding sites of 150 RBPs from eCLIP-seq with irreproducible discovery rate analysis (36) and with crosslink-centered regions identified from the DICER PAR-CLIP (38) and AGO1-4 PAR-CLIP (37) datasets by using BEDTools. Multiple binding sites for the same RBP in an intron were collapsed and counted as one binding site for that RBP. Functional annotations, localization patterns, and predicted RNA-binding domains of RBPs in the ENCODE eCLIP dataset were based on Table S2 of ref. (68). RBPs found in stress granules were as annotated in the RNA Granule and Mammalian Stress Granules Proteome (MSGP) databases (56, 69). The functional annotations, localization patterns, and RNA-binding domains of AGO1-4 and DICER were from the UniProt database (https://www.uniprot.org). Near full length FLEXI or other short intron reads were identified in ENCODE eCLIP datasets by intersecting mapped eCLIP reads with FLEXI or other short intron coordinates using BEDTools.

Gene expression changes in RBP knockdown datasets were calculated from changes in mRNA abundance measured by using DESeq2 (https://github.com/thelovelab/DESeq2) after GC correction with Salmon (https://github.com/COMBINE-lab/salmon) and conditional quantile normalization (CQN) (78). The number of significant gene expression changes (DESeq2-calculated adjusted p-value<0.05) for FLEXI host genes was compared to 3 gene sets containing the same number of randomly selected genes, and ANOVA was used to identify knockdowns that resulted in significantly more gene expression changes among FLEXI host genes than expected by random chance. To identify FLEXI host genes whose changes in mRNA levels were significantly biased in one direction, the numbers of upregulated and downregulated host genes containing a FLEXI with a binding site for the knocked-down RBP (defined as log_2_-transformed fold change>0, adjusted p-value<0.05) were compared to those for all significantly changed genes by Fisher’s exact test, and those with p-value <0.05 for difference in one or the other direction were considered significantly biased in that direction. Volcano plots were generated by plotting log_2_-transformed fold changes in the mRNA levels between the knockdown and control datasets versus -log_10_ adjusted p-values.

Changes in alternative splicing were calculated by using rMATS (https://rnaseq-mats.sourceforge.io). Coordinates of adjacent introns or of retained introns from splicing change files (MATS output) were intersected with coordinates of bound FLEXIs (*i.e*., FLEXIs that contain a binding site for a RBP), unbound FLEXIs (*i.e*., FLEXIs that do not contain a binding site for the same RBP), and other short introns, and the results were displayed as an empirical cumulative density plot (ECDF) showing the inclusion level difference for the same intron between the knockdown and control datatsets.

To map RBP-binding sites within introns, RBP-binding sites from ENCODE eCLIP datasets were intersected with FLEXI, other short intron, or long intron coordinates (BEDTools). Any binding site that overlapped with any portion of the intron was used. The location of the binding site with respect to normalized intron length was calculated from the distance between the midpoint of the annotated binding site and the beginning or end of the intron. Data were displayed as a density plot of the mid-point of the binding sites normalized by intron length.

For hierarchical clustering of the RBP-binding site co-enrichment, subsets of FLEXIs that have a binding site for each of 47 RBPs that have binding sites in ≥30 different FLEXI RNAs were extracted from datasets for each of the cellular RNA samples. Over- and under-represented RBPs in each of the 47 subsets for each cellular RNA sample were identified by generating a contingency table comparing the frequency of binding sites for the 126 RBPs that have an identified binding site in a FLEXI RNA (Fig. S9) in the tested subset to its frequency in all FLEXIs. Multiple binding sites for the same RBP in the same FLEXI were counted as one binding site. p-values were calculated using Fisher’s exact test and then adjusted by the Benjamini-Hochberg procedure and log_10_-transformed. RBP-binding sites were then key-coded as not significantly different or as significantly over- or under-represented (≥2% abundance, p≤0.05) in the tested subset of FLEXIs compared to all FLEXIs, and matrices of the Gower’s distance (79) were calculated using the key-coded information from the above analyses and used as input for hierarchical clustering by the complete linkage clustering method (72). The results were displayed as a heat map plotted using the R package Cluster and ComplexHeatmap, respectively. For hierarchical clustering analysis of the RBP-binding site co-enrichment in other short introns and long intron, 2,000 randomly selected other short or long introns for each category of intron were analyzed as above, with the process was repeated 10 times for each of the cellular RNA samples.

### Analysis of RBP-binding site in FLEXIs and RNA fragments of other short and long introns in nuclear and cytoplasmic fractions

FLEXIs detected by TGIRT-seq in nuclear and cytoplasmic fractions were analyzed by DESeq2. FLEXIs that have a fold change (FC) value > 1.5 in one fraction compared to the other were considered as differentially enriched in the corresponding fraction, and were denoted as nuclear or cytoplasmic FLEXIs, respectively. RNA fragments derived from other short and long introns were identified by intersecting mapped reads with other short intron or long intron coordinates extracted from Ensemble GRCh38 Release 93 gene annotations using BEDTools. Read pairs (Read 1 and Read 2) within the annotated introns were identified as intron RNA fragments and merged into a single bed file using BEDTools. RBP-binding in these RNA fragments were identified by intersecting the coordinates of FLEXI RNA, merged short intron, or long intron RNA fragments, with ENCODE annotated binding sites of 150 RBPs from eCLIP-seq with irreproducible discovery rate analysis (36) and with crosslink-centered regions identified from the DICER PAR-CLIP (38) and AGO1-4 PAR-CLIP (37) datasets by using BEDTools. Multiple binding sites for the same RBP in an intron were collapsed and counted as one binding site for that RBP.

### GO term enrichment analysis

GO term enrichment analysis of host genes encoding FLEXIs bound by different RBPs was performed by using DAVID bioinformatics tools (https://david.ncifcrf.gov) with all FLEXI host genes as the background. Hierarchical clustering was performed based on p-values for GO term enrichment of FLEXI host genes bound by the same RBPs using the gplots package in R (https://www.rdocumentation.org/packages/gplots/versions/3.1.3.1).

### Protein function and network analysis

Proteins identified as oncogene, tumor suppressor gene, or mitochondrial localized protein were based on Liu *et al*. (73), Zhao. *et al*.(74), and human MitoCarta3.0 (https://www.broadinstitute.org/mitocarta/mitocarta30-inventory-mammalian-mitochondrial-proteins-and-pathways). Protein network analysis was done using the STRING database (https://string-db.org) with the default setting.

## Data Availability

All TGIRT-seq datasets analyzed in this manuscript have been deposited in the National Center for Biotechnology Information Sequence Read Archive. These include datasets for: (i) commercial human plasma pooled from multiple healthy individuals from a previous study (20) (accession number PRJNA640428); (ii) biological replicates of unfragmented cellular RNAs from HEK-293T cells from a previous study (26) (samples WC and YC; accession number (PRJNA389469); (iii) TGIRT-seq datasets for biological replicates of unfragmented cellular RNAs from K-562, HEK-293T, HeLa S3, MCF7, MDA-MB-231 and UHRR (accession number PRJNA648481); (iv) K-562 cells, samples UFC1-7; MDA-MB-231, samples MDA1-5; and UHRR, samples UHRR0-9 (accession number PRJNA722757); (v) TGIRT-seq datasets of whole-cell RNA and nuclear and cytoplasmic fractions from K-562, HeLa S3, MD-MBA-231, andMCF7 (accession number PRJNA648481).

---

**Table S1.** Summary of datasets obtained in this study.

| Cell / Tissue | Raw reads<br>(x10<sup>6</sup>) | Trimmed<br>reads (x10<sup>6</sup>) | Mapped<br>reads (x10<sup>6</sup>) | Mapped to<br>feature (x10<sup>6</sup>) |
| --- | --- | --- | --- | --- |
| HEK-293T (Bio. 1)<sup>*</sup> | 741.6 | 726.5<br>(98.0%) | 715.2<br>(98.5%) | 690.9<br>(96.6%) |
| HeLa S3<sup>†</sup> | 851.0 | 803.3<br>(94.4%) | 768.4<br>(95.7%) | 705.6<br>(92.0%) |
| K-562 (Bio. 1)<sup>§</sup> | 744.2 | 725.0<br>(97.4%) | 713.8<br>(98.4%) | 698.5<br>(97.9%) |
| UHRR (Bio. 1)<sup>‡</sup> | 712.0 | 682.2<br>(95.8%) | 666.3<br>(97.7%) | 630.9<br>(94.7%) |
| Plasma<sup>¶</sup> | 134.5 | 122.7<br>(91.3%) | 71.1<br>(57.9%) | 61.7<br>(87.2%) |
| MCF7 | 757.8 | 703.7<br>(92.9%) | 692.1<br>(98.4%) | 673.7<br>(97.3%) |
| MDA-MB-231 (Bio. 1)<sup>#</sup> | 226.1 | 211.3<br>(93.5%) | 207.5<br>(98.2%) | 202.2<br>(97.5%) |
| HEK-293T (Bio. 2)<sup>‡‡</sup> | 224.1 | 211.0<br>(94.2%) | 180.8<br>(85.7%) | 162.6<br>(89.9%) |
| K-562 (Bio. 2)<sup>¶¶</sup> | 59.4 | 54.6<br>(91.9%) | 47.9<br>(87.7%) | 45.5<br>(95.1%) |
| MDA-MB-231 (Bio. 2)<sup>##</sup> | 314.8 | 274.3<br>(87.1%) | 259.1<br>(94.5%) | 243.7<br>(94.1%) |
| UHRR (Bio. 2)<sup>§§</sup> | 416.4 | 397.3<br>(95.4%) | 359.4<br>(90.5%) | 293.0<br>(81.5%) |
| HeLa S3 (Total RNA)<sup>**</sup> | 46.8 | 27.2<br>(58.2%) | 20.5<br>(75.1%) | 13.3<br>(65.0%) |
| HeLa S3 (Nucleus RNA)<sup>**</sup> | 37.4 | 27.5<br>(73.5%) | 22.4<br>(81.4%) | 15.3<br>(68.5%) |
| HeLa S3 (Cytoplasm RNA)<sup>**</sup> | 49.7 | 29.3<br>(59.1%) | 21.1<br>(71.9%) | 13.2<br>(62.7%) |
| K-562 (Total RNA)<sup>**</sup> | 66.2 | 49.7<br>(75.1%) | 37.5<br>(75.4%) | 24.3<br>(64.8%) |
| K-562 (Nucleus RNA)<sup>**</sup> | 75.5 | 66.7<br>(88.4%) | 56.5<br>(84.6%) | 34.4<br>(60.9%) |
| K-562 (Cytoplasm RNA)<sup>**</sup> | 78.4 | 59.4<br>(75.8%) | 40.9<br>(68.8%) | 25.4<br>(62.0%) |
| MDA-MB-231 (Total RNA)<sup>**</sup> | 82.0 | 74.1<br>(90.3%) | 57.4<br>(77.5%) | 38.7<br>(67.3%) |
| MDA-MB-231 (Nucleus RNA)<sup>**</sup> | 108.6 | 102.3<br>(94.2%) | 87.7<br>(85.7%) | 54.3<br>(61.9%) |
| MDA-MB-231 (Cytoplasm RNA) ** | 108.1 | 92.6<br>(85.7%) | 65.2<br>(70.4%) | 42.0<br>(64.4%) |
| MCF7 (Total RNA) ** | 70.7 | 63.8<br>(90.2%) | 49.3<br>(77.3%) | 33.3<br>(67.6%) |
| MCF7 (Nucleus RNA) ** | 89.7 | 85.0<br>(94.8%) | 73.7<br>(86.7%) | 48.2<br>(65.5%) |
| MCF7 (Cytoplasm RNA) ** | 109.9 | 94.9<br>(86.3%) | 68.3<br>(72.0%) | 43.6<br>(63.9%) |

\*HEK-293T cell RNA, combined datasets from 8 technical replicates (SRA BioProject accession number PRJNA648481, HEK-rep1 to 8).
†HeLa S3 cell RNA, combined datasets from 10 technical replicates (SRA BioProject accession number PRJNA648481, HeLa-rep1 to 10).
§K-562 cell RNA, combined datasets from 8 technical replicates (SRA BioProject accession number PRJNA648481, K-562-rep1 to 8).
‡Universal human reference RNA, combined datasets from 8 technical replicates (SRA BioProject accession number PRJNA648481, UHRR-rep1 to 8, lot# 0006390163).
¶Commercial human plasma pooled plasma from healthy individuals. Fifteen combined datasets (SRA BioProject accession number PRJNA640428, samples DNaseI\_1 to 12, ExoI\_1 to 3) (Yao *et al.* 2020).
||MCF7 RNA, combined datasets from 8 technical replicates (SRA BioProject accession number PRJNA648481, MCF-rep1 to 8).

**Table S2.** Abundance of sncRNAs (RPM) detected by TGIRT-seq in cellular RNA samples compared to literature copy number per cell values for these RNAs.

| Name | RPM |  |  |  | Copies/cell* | Type |
| --- | --- | --- | --- | --- | --- | --- |
|  | K-562 | HEK-293T | HeLa S3 | UHRR |  |  |
| U1 | 1,385/6,057 | 1,266/4,756 | 1,337/5,732 | 2,647/7,977 | 1x10<sup>6</sup> | Major spliceosomal snRNA |
| U2 | 5,764/9,970 | 9,476/1,1977 | 28,774/26,539 | 9,787/12,135 | 5x10<sup>5</sup> | Major spliceosomal snRNA |
| U4 | 65/371 | 34/172 | 540/1,815 | 76/280 | 2x10<sup>5</sup> | Major spliceosomal snRNA |
| U5 | 1,146/5,476 | 2,703/9,161 | 13,241/26,325 | 2,715/8,247 | 2x10<sup>5</sup> | Major spliceosomal snRNA<br>Minor spliceosomal snRNA |
| U6 | 974/3,474 | 1,766/5,204 | 923/4,863 | 1,355/4,947 | 4x10<sup>5</sup> | Major spliceosomal snRNA |
| U7 | 3/22 | 4/27 | 49/197 | 3/9 | 4x10<sup>3</sup> | U7 |
| U11 | 2/32 | 3/33 | 11/108 | 8/44 | 1x10<sup>4</sup> | Minor spliceosomal snRNA |
| U12 | 36/157 | 24/78 | 192/433 | 41/110 | 5x10<sup>3</sup> | Minor spliceosomal snRNA |
| U4ATAC | 37/183 | 21/101 | 1,291/2,883 | 32/173 | 2x10<sup>3</sup> | Minor spliceosomal snRNA |
| U6ATAC | 204/1,104 | 286/1,481 | 699/4,236 | 268/1,350 | 2x10<sup>3</sup> | Minor spliceosomal snRNA |
| RN7SK | 1,368/3,585 | 2,771/4,828 | 5,043/11,318 | 7,174/6,428 | 2x10<sup>5</sup> | 7SK |
| RN7SL | 3,445/10,723 | 8,080/16,276 | 4,689/13,770 | 19,801/17,395 | 5x10<sup>5</sup> | 7SL |
| RPPH1 | 165/474 | 245/643 | 257/713 | 1,293/2,212 | 2x10<sup>5</sup> | RNase P RNA component |
| SNORD3 | 7,637/13,928 | 12,700/18,193 | 23,204/29,379 | 38,531/30,293 | 2x10<sup>5</sup> | C/D box snoRNA, U3 |
| SNORD118 | 222/758 | 985/1,617 | 552/948 | 745/1,178 | 4x10<sup>4</sup> | C/D box snoRNA, U8 |
| SNORD13 | 436/2,572 | 367/2,129 | 817/3,477 | 1,223/4,786 | 1x10<sup>4</sup> | C/D box snoRNA, U13 |
| SNORD14 | 4,278/6,920 | 3,887/5,651 | 9,979/9,149 | 11,073/9,899 | 1x10<sup>4</sup> | C/D box snoRNA, U14 |
| SNORD22 | 362/1,600 | 240/963 | 748/2,526 | 947/2,547 | 1x10<sup>4</sup> | C/D box snoRNA, U22 |
| RMRP | 669/1,907 | 1,776/3,781 | 2,211/7,294 | 3,276/6,048 | 1x10<sup>5</sup> | MRP |

\*Copy number per cell values for sncRNAs in vertebrate cells (Tycowski *et al.*, 2006) used in the linear regression analysis of Fig. S6.

**Table S3.**
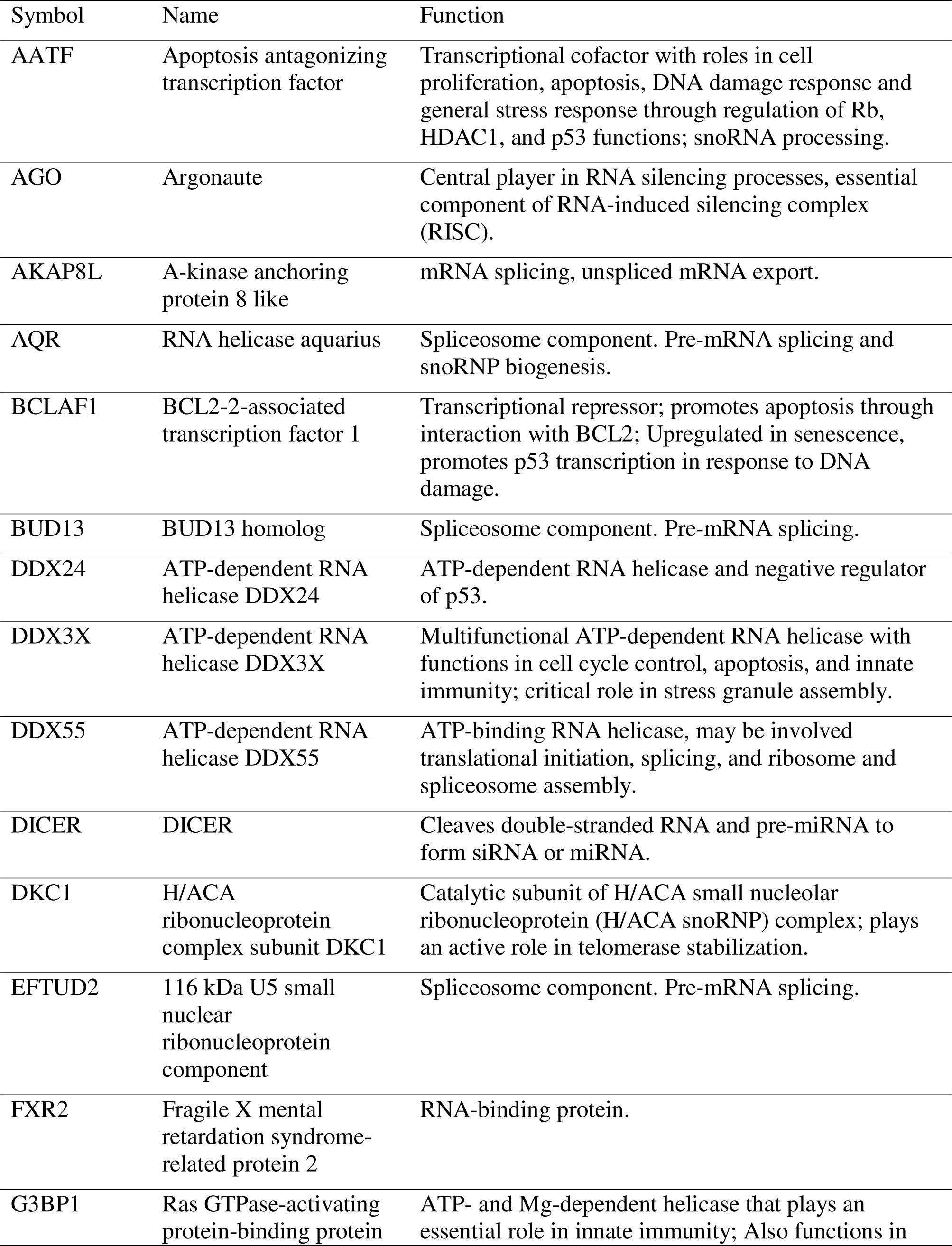
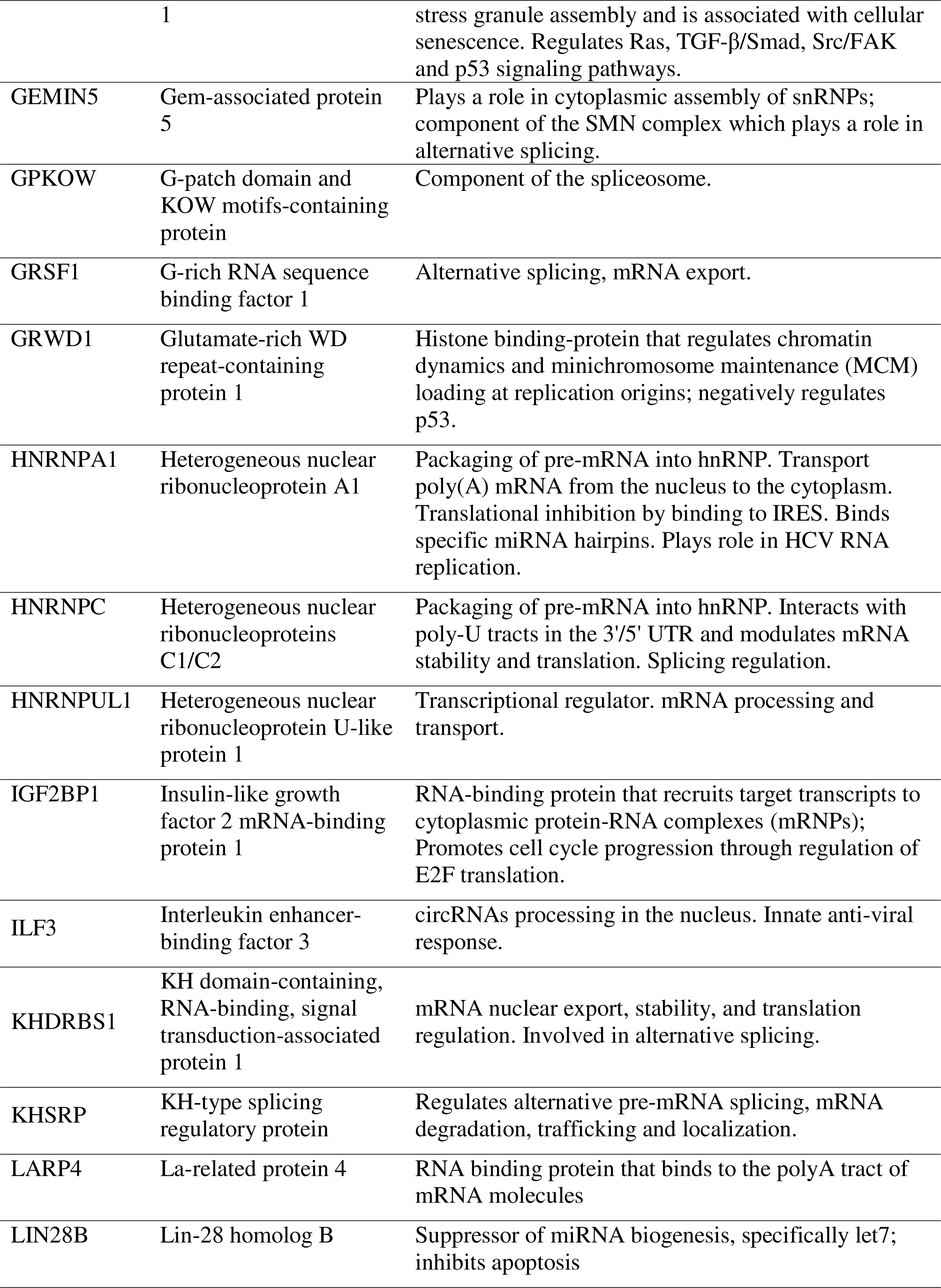
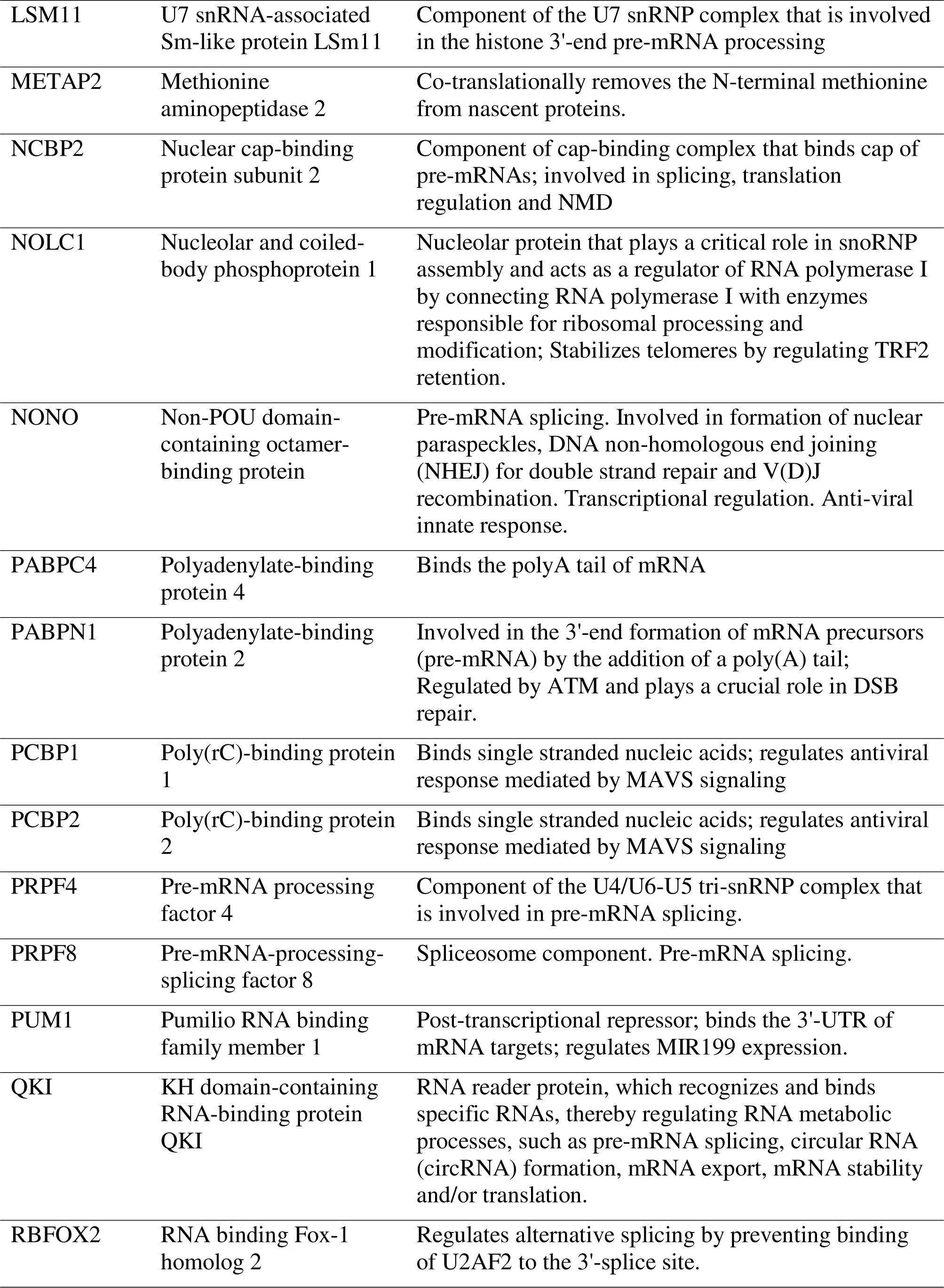
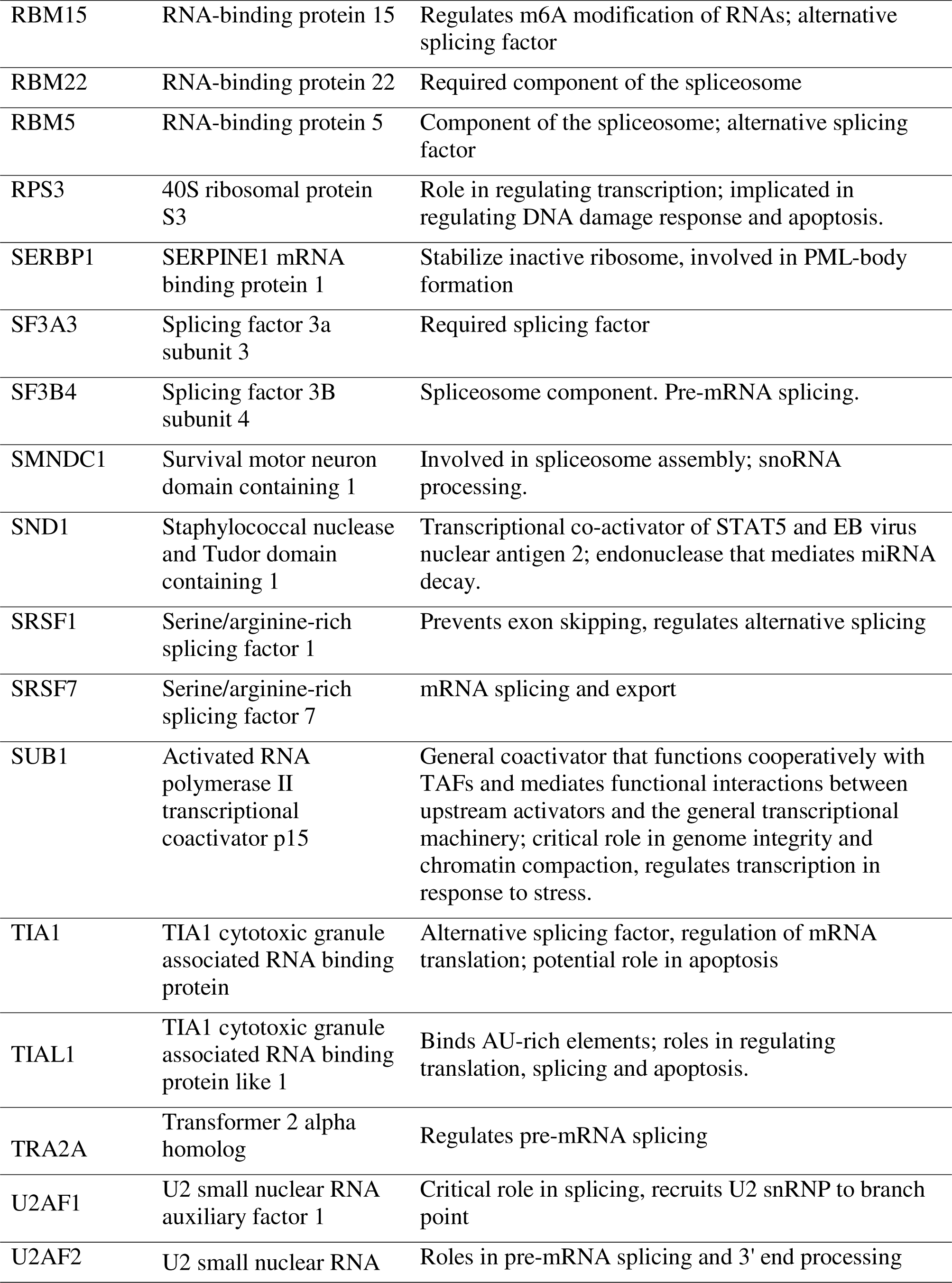
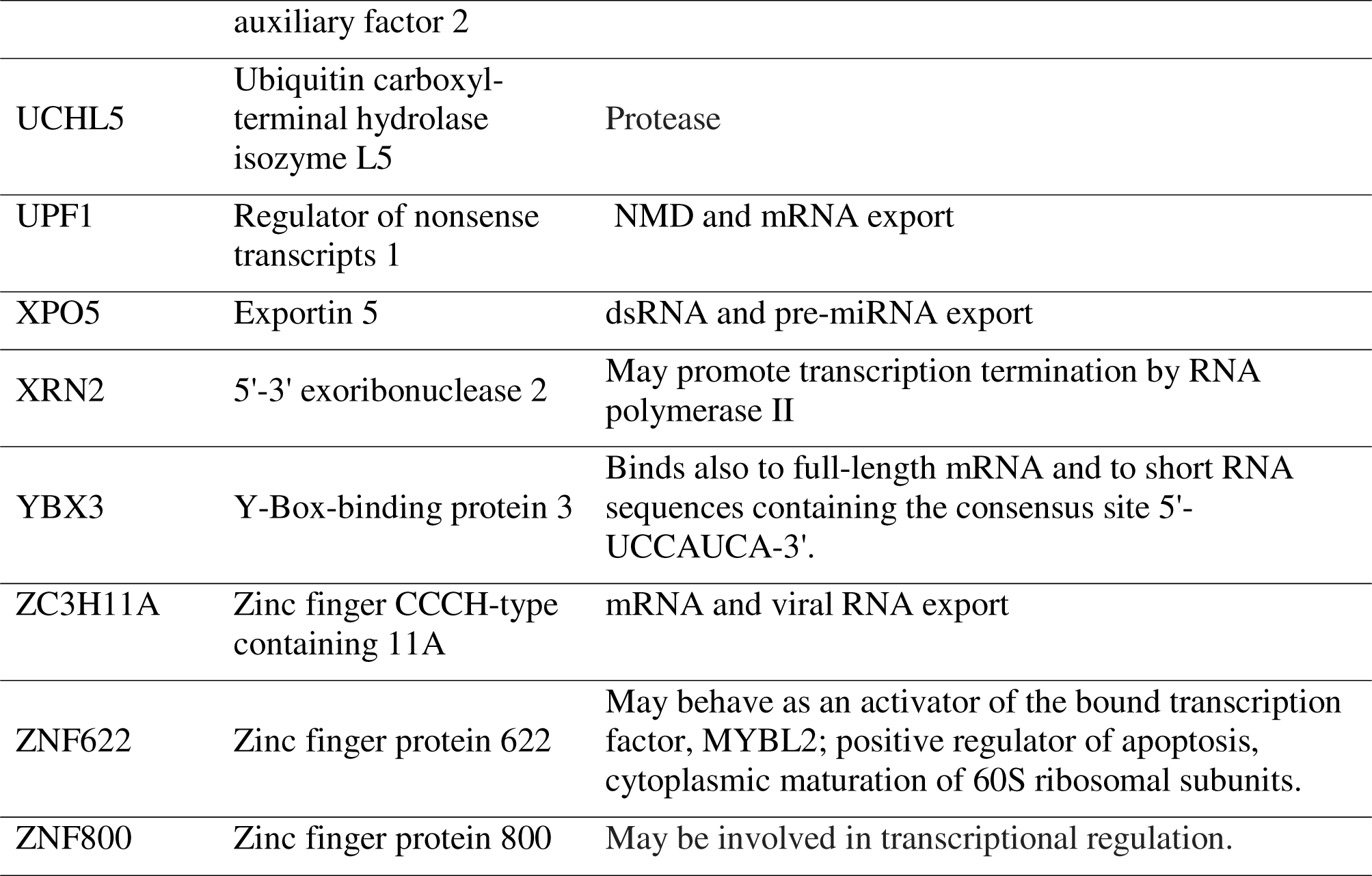
Proteins that bind FLEXIs.

**Table S4.**
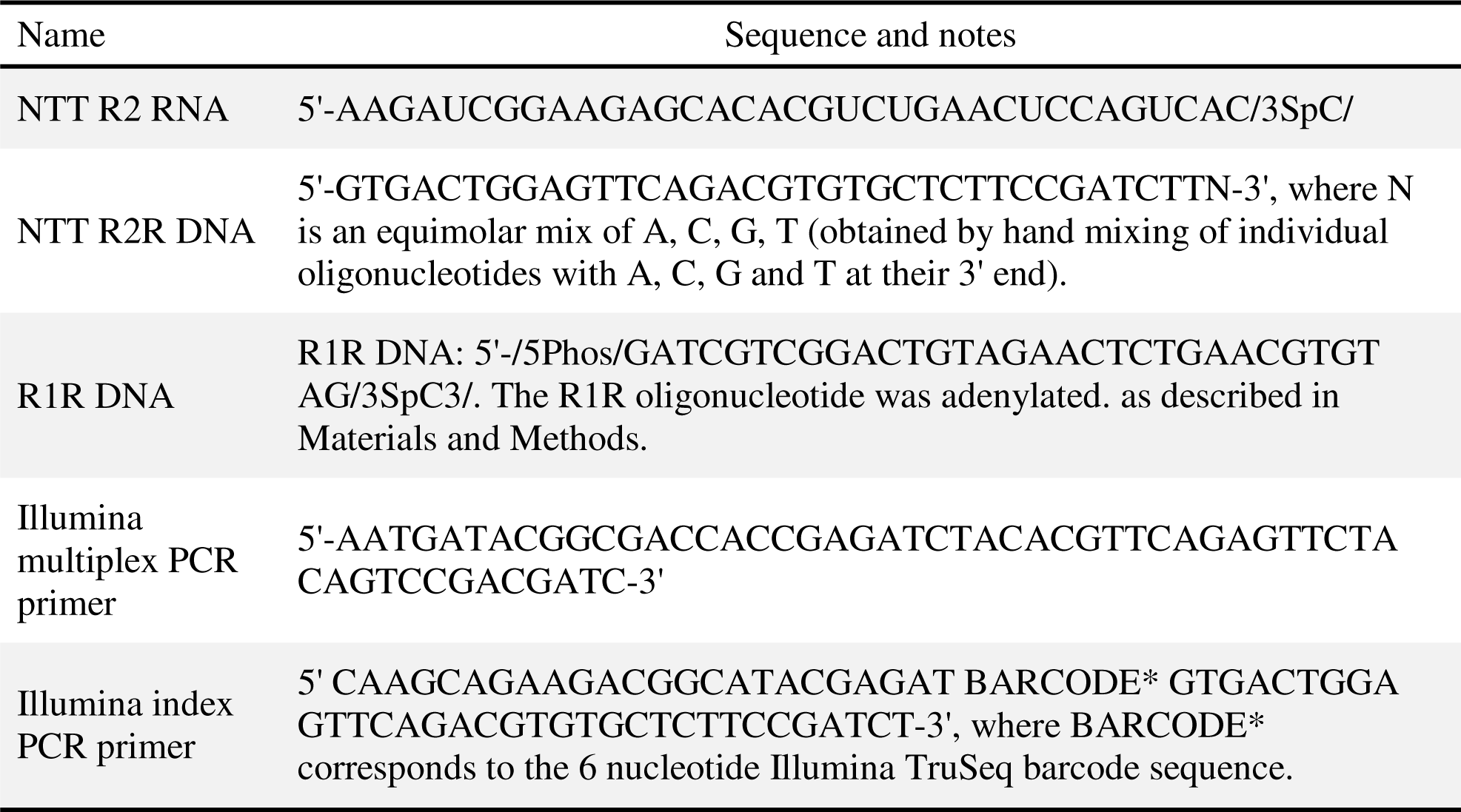
Oligonucleotides used for construction of TGIRT-seq libraries.

**Table S5.**
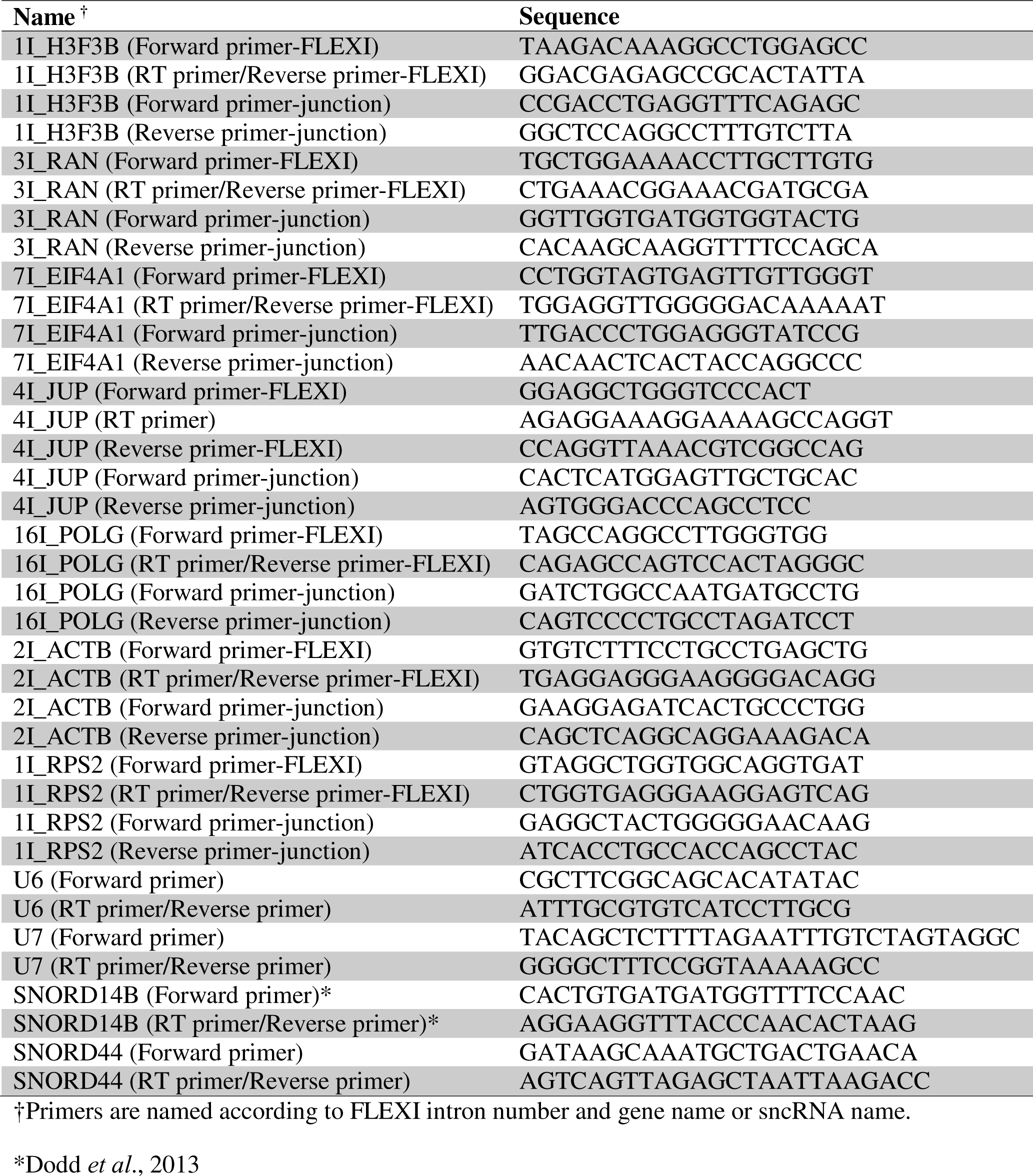
Oligonucleotides used for ddPCR and exonuclease digestion assays.

